# Exploiting the inherent promiscuity of the acyl transferase of the stambomycin polyketide synthase for the mutasynthesis of analogues

**DOI:** 10.1101/2024.10.29.620781

**Authors:** Li Su, Yaouba Souaibou, Laurence Hôtel, Christophe Jacob, Peter Grün, Yan-Ni Shi, Alicia Chateau, Sophie Pinel, Helge B. Bode, Bertrand Aigle, Kira J. Weissman

## Abstract

The polyketide specialized metabolites of bacteria are attractive targets for generating analogues, with the goal of improving their pharmaceutical properties. Here, we aimed to produce C-26 derivatives of the giant anti-cancer stambomycin macrolides using a mutasynthesis approach, as this position has been shown previously to directly impact bioactivity. For this, we leveraged the intrinsically broad specificity of the acyl transferase domain (AT_12_) of the modular polyketide synthase (PKS), which is responsible for the alkyl branching functionality at this position. Feeding of a panel of synthetic and commercially available dicarboxylic acid ‘mutasynthons’ to an engineered strain of *Streptomyces ambofaciens* (Sa) deficient in synthesis of the native α-carboxyacyl-CoA extender units, resulted in six new series of stambomycin derivatives as judged by LC-HRMS and NMR. Notably, the highest product yields were observed for substrates which were only poorly accepted when AT_12_ was transplanted into a different PKS module, suggesting a critical role for domain context in the overall functioning of PKS proteins. We also demonstrate the superiority of this mutasynthesis approach − both in terms of absolute titers and yields relative to the parental compounds − in comparison to the alternative precursor-directed strategy in which monoacid building blocks are supplied to the wild type strain. We further identify a malonyl-CoA synthetase, MatB_Sa, with specificity distinct from previously identified promiscuous enzymes, making it a useful addition to a mutasynthesis toolbox for generating atypical, CoA activated extender units. Finally, we show that two of the obtained (deoxy)-butyl-stambomycins exhibit antibacterial and antiproliferative activities similar to the parental stambomycins, while an unexpected butyl-demethyl congener is less potent. Overall, this works confirms the interest of biosynthetic pathways which combine a dedicated route to extender unit synthesis and a broad specificity AT domain for producing bioactive derivatives of fully-elaborated complex polyketides.

## Introduction

The reduced polyketide specialized metabolites of bacteria exhibit stunning structural diversity. One source of this architectural variety is the multiple acyl-CoA building blocks incorporated during chain building. This construction process is carried out by mega-enzyme assembly lines called type I polyketide synthases (PKSs), which comprise a series of functional modules distributed among multiple polypeptide subunits. The modules are composed of sets of catalytic domains (minimally ketosynthase (KS) and acyl transferase (AT)) which act in partnership with non-catalytic acyl carrier proteins (ACPs) to elongate the chains, followed by optional modification of each newly-incorporated monomer. Both the ‘starter units’ which initiate the biosynthesis and the ‘extender units’ used in chain elongation are selected by the AT domains. These domains thus govern to a large extent the branching functionality within polyketide structures. Correspondingly, manipulation of AT domains using genetic engineering offers a powerful potential means for the regioselective diversification of polyketide skeletons, especially in the case of ATs which exhibit unusual substrate specificity relative to the typical choice of malonyl-CoA or methylmalonyl-CoA^1^ (e.g. for propyl-, butyl-, allyl-malonyl-CoA, etc.^2-4^). To date, a variety of approaches including AT domain exchange, *in trans* AT complementation, and active site-directed mutagenesis have been applied to polyketide structural engineering with varying levels of success^5^.

Experimentally, AT swapping consists of exchanging only the AT domain within a module of interest for an AT of different substrate specificity from another module, either sourced from the same or a different PKS^6-10^. This approach has resulted in various hybrid metabolites, with recent work revealing highly effective domain boundaries for such exchanges^11, 12^. Concerning the complementation strategy, it involves selectively inactivating the native AT within a particular PKS module and rescuing its activity via a co-expressed *trans*-acting AT^4, 13, 14^. Drawbacks of this approach include that it depends on the ability of the *trans-*ATs to recognize noncognate ACPs as partners^14^, and the typically limited substrate (for malonate)^4^ of these enzymes. Finally, AT active site mutagenesis has been used in efforts to redirect and/or broaden innate AT substrate selectivity^15-17^. Despite the availability of several AT crystal structures^18-21^, this approach remains hampered by the paucity of co-complex structures in the presence of native extender units. Indeed, all such attempts to date to switch AT specificity have only resulted in a shift in substrate preference^4, 15^-^17, 22^, a result potentially explained by the currently insufficient predictive power of the identified AT substrate-binding motifs^12^.

An alternative approach towards polyketide structural diversification is to exploit the intrinsically broad substrate specificity of certain ATs, and challenge them with alternative building blocks via precursor-directed biosynthesis (PDB) and/or mutasynthesis^23-31^. In the case of PDB (**Figure 1a**), the supplied ‘mutasynthons’ must compete with the native substrates, while in a mutasynthetic strategy (**Figure 1b**), precursor biosynthesis is disabled^2^, and thus the pathway depends, at least in principle, on the presence of the exogenous building block in order to function. To date, multiple functional groups including several not found naturally in polyketides have been incorporated into polyketides in this manner, including alkyne, azide, and halogen moieties^24-31^.

**Figure 1.**
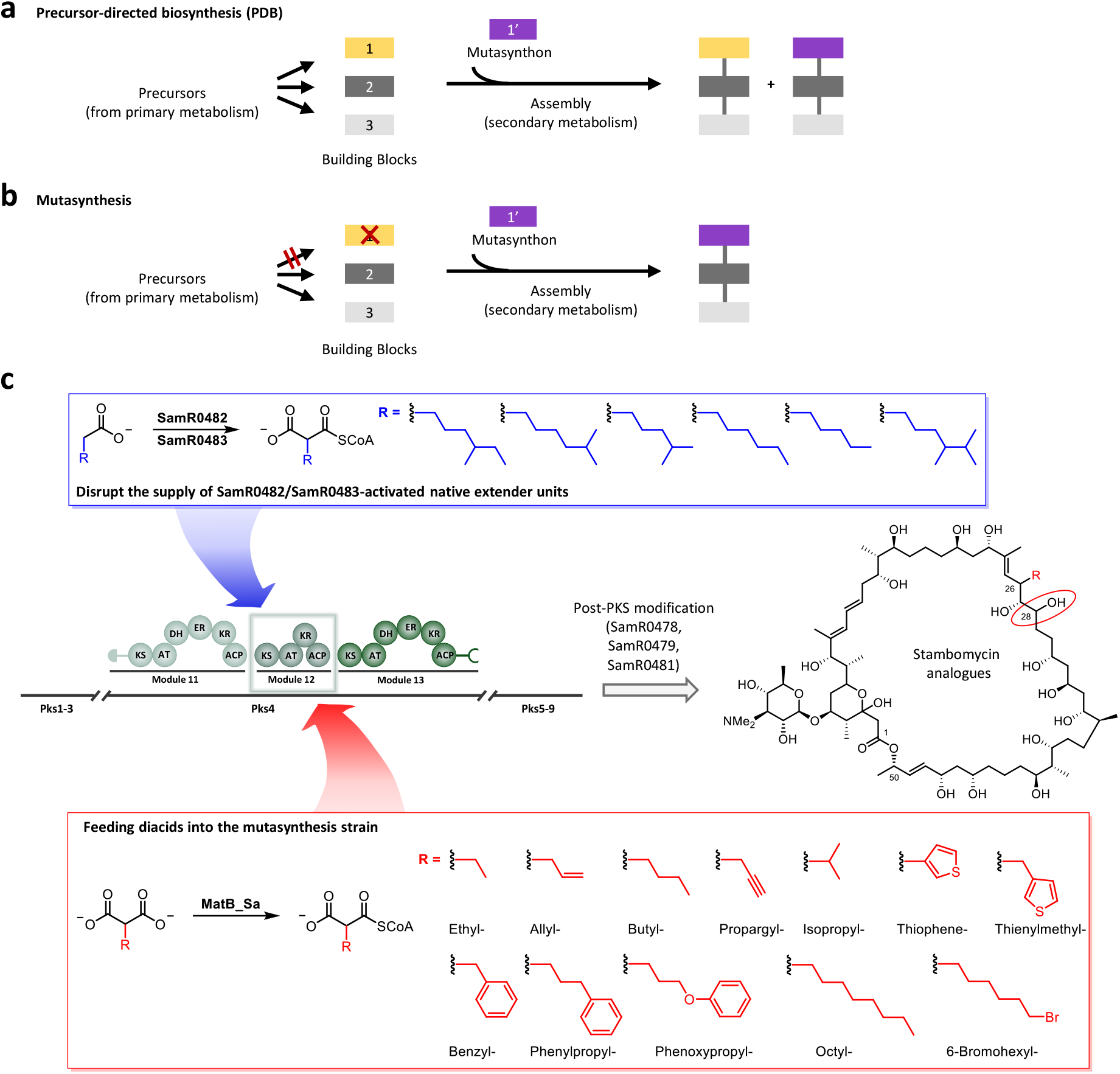
Schematic overview of the (a) Precursor-directed biosynthesis (PDB) and (b) mutasynthesis approaches^23^, and the (c) mutasynthesis strategy applied to stambomycin biosynthesis. Commercially sourced and synthetic malonic acids incorporating a wide variety of side chains were fed to the production culture of mutasynthesis mutant ATCC/OE484/Δ483. At least six of these alternative extender units were then successfully activated to their corresponding CoAs by the promiscuous MatB_Sa of *S. ambofaciens*, and incorporated into the pathway by AT_12_, resulting in stambomycin derivatives bearing modifications at C-26, a position which directly impacts the bioactivity^42^. The hydroxylation at C-28 (red circle) was found to occur to variable extents, but the hydroxylation at C-50 and mycaminosylation went to completion.

Evidently, a critical parameter for successful PDB and/or mutasynthesis is exogenous precursor supply. Previous *in vitro* studies have shown that commercially available malonyl-CoA derivatives can be used as mutasynthons^32^. However, this method is not suitable for *in vivo* feeding studies due to poor acyl-CoA membrane permeability^33^. This problem can be addressed by the use of synthetic *N*-acetylcysteamine thioesters (SNACs) which mimic the distal end of CoA, but leveraging them in large-scale applications is complicated by the need to synthesize the compounds, as well as their cellular toxicity^34^. A promising alternative is a chemoenzymatic approach, in which a fed monomer (mono-or di-acid) is activated to its corresponding malonyl-

CoA derivative *in cellulo* by an appropriate enzyme − either a crotonyl-CoA reductase/carboxylase (CCR)^35-38^ homologue or a broad-specificity malonyl-CoA synthetase (MatB) ^18, 34, 39-41^ homologue (**Figure S1a, b**).

ATs which recognize atypical extender units typically show broad substrate specificity relative to their malonate-or methylmalonate equivalents^12, 38^. Even among such domains, a particularly promiscuous AT participates in the biosynthesis of the stambomycin family of 51-membered macrolides in *S. ambofaciens* ATCC23877^42-44^. This domain (AT_12_) naturally recruits at least six atypical alkyl malonyl-CoA-derived extender units: (4-methylhexyl)malonyl-CoA, (5-methylhexyl)malonyl-CoA, (4-methylpentyl)malonyl-CoA, *n*-hexylmalonyl-CoA, *n*-pentylmalonyl-CoA and (4,5-dimethylhexyl)malonyl-CoA (**Figure 1c**). This broad specificity results in six stambomycins (A−F, respectively) bearing distinct side chains at C-26. Notably, modifications at this position directly impact the biological properties of the metabolites, with stambomycins C/D showing improved antiproliferative activity towards a range of human cancer cells^42^. In contrast to the typical CCR-mediated synthesis of polyketide extender units via reductive carboxylation of α,β-unsaturated acyl-thioesters^35-38^, the stambomycin precursors are generated by direct carboxylation of medium chain acyl-CoA substrates catalyzed by the ATP-dependent enzyme SamR0483 (although SamR0483 was renamed MccB for medium chain acyl-CoA carboxylase β-subunit, we will refer to it here as SamR0483 in line with the gene nomenclature)^44^ (**Figure S1c**). The acyl-CoAs derive either from the corresponding amino acids via the consecutive action of the branched-chain α-keto acid dehydrogenase complex and primary metabolic fatty acid synthase enzymes (FabH and FabF), or via direct conversion of the corresponding fatty acids to their CoA thioesters by SamR0482, a medium chain fatty acyl-CoA ligase homologue^44^. Notably, SamR0482 and SamR0483 together were shown to generate the corresponding extender units from 6-azidohexanoic acid and 8-nonyoic acid, resulting in stambomycin C-26 analogues bearing azide and alkyne groups. Thus, in addition to the intrinsically broad specificity of AT_12_ and all twelve downstream PKS modules, Sam0482 and SamR0483 exhibit useful substrate tolerance.

Here we aimed to further exploit the intrinsic flexibility of the stambomycin pathway to access a larger panel of analogues by mutasynthesis for biological testing (**Figure 1c**). For this, we first disrupted the supply of the native extender units by mutational inactivation of *samR0483*. We then fed the resulting strains with 12 commercially-available or synthetic malonic acid derivatives, relying either on the intrinsic promiscuity of the *S. ambofaciens* MatB (MatB_Sa) or heterologously-expressed MatB from *S. cinnamonensis*^45^ for activation to their respective CoA thioesters. Six of the derivatives resulted in stambomycin analogues incorporating or lacking hydroxylation at C-28, including four at yields sufficient for structure elucidation by NMR and for biological activity tests. We further show that MatB_cinna is more effective than the native *S. ambofaciens* homologue at monomer activation, while direct comparison of the yields obtained by PDB and mutasynthesis confirms the relative utility of the latter approach for analogue generation.

## Results

### Construction of a mutant for stambomycin mutasynthesis

In order to disable the native pathway to the six extender units, the genes encoding SamR0482 and SamR0483 were deleted using PCR-targeting, resulting in the mutants ATCC/Δ482^46^ and ATCC/Δ483 (**Tables S1**−**S4**). Subsequently, the LAL regulator overexpressing plasmid pOE484 was introduced into the mutants as previously described to trigger stambomycin biosynthesis^42^, affording the mutants ATCC/OE484/Δ482^46^ and ATCC/OE484/Δ483 (**Table S1**). Extracts of mutants ATCC/OE484/Δ482 and ATCC/OE484/Δ483 were then analyzed by LC-HRMS in comparison to the control strain ATCC/OE484. This analysis revealed that while production by ATCC/OE484/Δ483 was almost completely abolished (**Figure 2**), the ATCC/OE484/Δ482 mutant continued to produce stambomycins at approximately 51% yield relative to the wild type strain (**Figure 2**). BlastP analysis^47^ of the *S. ambofaciens* genome using SamR0482 as a query revealed several homologues, two of which are located in the antimycin and congocidine clusters: AntF (acyl-CoA synthetase, 29.8% amino acid identity to SamR0482) and Cgc22 (acyl-CoA synthetase, 34.5% amino acid identity to SamR0482). We thus propose that in addition to provision of the necessary acyl-CoAs via core metabolism, the loss-of-function of SamR0482 may be compensated by either AntF, Cgc22 or both (**Table S1**). BlastP analysis using SamR0483 also revealed several homologues, notably including the duplicated enzyme AlpX (proposed to be involved in kinamycin biosynthesis^48, 49^) and two propionyl-CoA carboxylase (PCC) β-subunits which show 59% and 52% amino acid identity to SamR0483 and high mutual similarity (**Table S1**). However, PccBs typically act on short chain acyl-CoAs^50^, potentially explaining why only a low level of SamR0483 complementation was observed (0.3% stambomycin yield relative to wild type).

**Figure 2.**
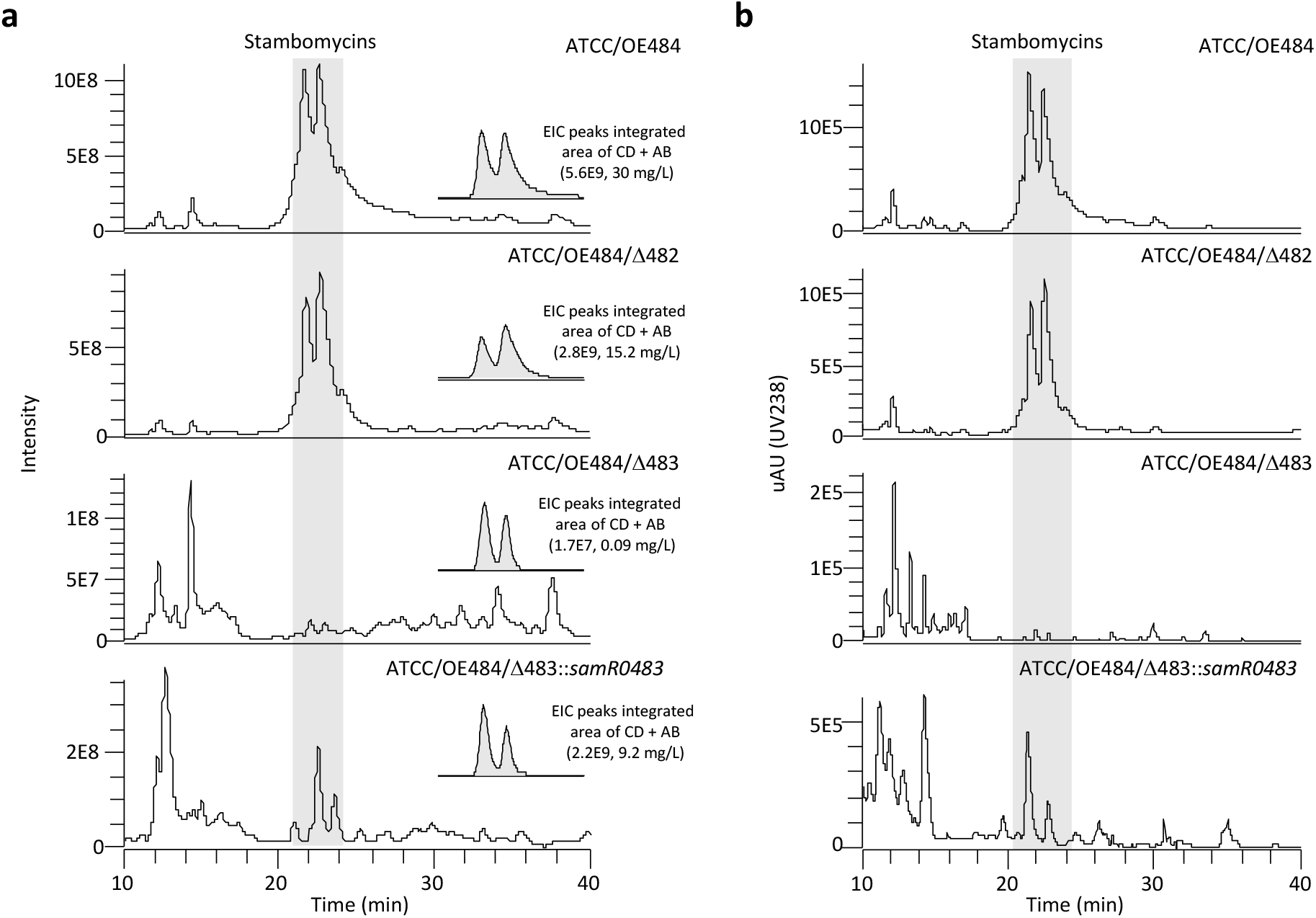
Analysis by (a) mass spectrometry and (b) UV at λ_max_ 238 nm of ATCC/OE484 (control) and mutants for mutasynthesis. The TIC (total ion chromatogram) and UV spectrum of ATCC/OE484/Δ483 clearly show that production by the mutant of the stambomycins (peaks highlighted in grey) is almost abolished. The indicated yields of stambomycins (AB and CD) in the wild type and mutant strains are based on integration of the EIC (extracted ion chromatogram) peaks and a previously generated standard curve for stambomycin AB^43^. The calculated areas represent the average of the analysis of at least two independent clones of each strain.

To confirm that the substantial decrease in stambomycin production in the ATCC/OE484/Δ483 mutant was due to the loss function of SamR0483, we carried out a complementation experiment by integrating the *samR0483* gene into the genome of ATCC/OE484/Δ483 mutant under the control of the constitutive promoter *ermEp*\*^51^. Although the observed complementation was incomplete (31% stambomycin yields relative to wt (**Figure 2**)), restoration of biosynthesis nonetheless supports the proposed essential role for SamR0483 in the pathway^44^. We therefore used mutant ATCC/OE484/Δ483 as our base strain for the mutasynthesis experiments (**Figure 2**).

### Mutasynthesis of stambomycin analogues

BlastP analysis against the genome sequence of *S. ambofaciens* using the MatB_Sc (*S. coelicolor*)^40^ as the query revealed an enzyme (referred to here as MatB_Sa) with amino acid identity of 90% to MatB_Sc. Sequence alignment also showed that MatB_Sa shares 75% and 39% amino acid identity with MatB_cinna (*S. cinnamonesis*)^34^ and MatB_Rt (*Rhizobium trifolii*)^41^, respectively (**Figure S2**). The high homology between MatB_Sa and the demonstrably promiscuous MatB_cinna^34^ suggested that MatB_Sa might also possess the necessary broad specificity to efficiently activate non-native malonate derivatives to their corresponding CoAs. On this basis, individual cultures of the ATCC/OE484/Δ483 mutant strain were supplemented with a wide panel of commercially sourced and synthetic diacids, including ethyl-, isopropyl-, allyl-, butyl-, benzyl-, octyl-, phenylpropyl-, phenoxypropyl-, thiophene-, thienylmethyl-, propargyl-, and 6-bromohexyl-malonic acids (**Figure 1c**) at a final concentration of 10 mM, in parallel with a control (*i*.*e*. a culture of the mutant strain lacking added malonic acid derivatives).

Analysis of cell pellets following extraction with methanol by LC-HRMS showed that the supplementation with ethyl-(giving rise to compound **1**), allyl-(**2** and **3**), butyl-(**4** and **5**), benzyl-(**6** and **7**), phenoxypropyl-(**9** and **10**), and 6-bromohexyl-malonic acids (**11**) yielded new peaks relative to the control, all with masses consistent with the expected C26-modified stambomycin analogues (**Table 1** and **Figure S3a, c, d, e, h, l**). It must nonetheless be noted that the exact mass of the 6-bromohexyl-malonic acid analogue corresponds to a structure lacking two of the expected hydrogens (**Figure S3l**). This structural variation may correspond to a ketone at position C-28 instead of the expected hydroxyl functionality, due to over-oxidation by the P450 hydroxylase SamR0478^52^. Indeed, SamR0478 belongs to the CYP107H family whose members have been shown to catalyse ketonization of C-H bonds, but this hypothesis remains to be verified). In contrast, the phenylpropyl-, thiophene-, thienylmethyl-, and propargyl-malonic acids failed to be incorporated (**Table 1, Figure S3b, g, i, j, k** and **Tables S5−S7**). A peak at 1390.9546 ([M+H]^+^, C_74_H_130_NO_22_^+^; r.t. 25.4 min) found in the octyl-fed extract (**Table 1**) requires further investigation, in order to determine whether it corresponds to a novel octyl-stambomycin (**8**) or the native stambomycin F^53^ (r.t. 24.2 min), as the two masses are identical (**Table S5**). Arguing for the formation of octyl-stambomycin, however, the observed yield of the metabolite is 9.2% relative to the stambomycins A−D (**Figure S3f**), while the typical titers of stambomycin F are only 2.5% (**Figure S4a**).

**Table 1.** Production of stambomycins and stambomycin analogues (the observed compounds are numbered) resulting from mutasynthesis strain ATCC/OE484/Δ483 in the presence of supplemental malonic acid derivatives (final concentration of 10 mM). The amount of stambomycins (A–D) produced by ATCC/OE484/Δ483 in the absence of supplementation was set to 100%. For details, see **Figure S3**.

| Malonic acid derivatives | Production relative to stambomycins (A–D) by the control |  |  |
| --- | --- | --- | --- |
|  | Stambomycins<br>(A–D) | Analogue<br>(with C-28 OH) | Deoxy analogue<br>(lacking C-28 OH) |
| Ethyl- | 853% | n.d. | 0.86% (**1**) |
| Isopropyl- | 361% | n.d. | n.d. |
| Allyl- | 189% | 2.05% (**2**) | 378% (**3**) |
| Butyl- | 192% | 4326% (**4**) | 20619% (**5**) |
| Benzyl- | 478% | 27.5% (**6**) | 54.8% (**7**) |
| Octyl- | 845% | 77.5% (**8**) | n.d. |
| Phenylpropyl- | 591% | n.d. | n.d. |
| Phenoxypropyl- | 233% | 10.9% (**9**) | 13.5% (**10**) |
| Thiophene- | 83.9% | n.d. | n.d. |
| Thienylmethyl- | 99.4% | n.d. | n.d. |
| Propargyl- | 458% | n.d. | n.d. |
| 6-Bromohexyl- | 611% | 27.5% (**11**) | n.d. |

Biosynthesis of the parental stambomycins involves C-28 hydroxylation by the P450 hydroxylase SamR0478, as well as on-line C-50 hydroxylation by a second P450 SamR0479 to generate the nucleophile required for macrolide formation^54^. For five of the successfully incorporated precursors (ethyl-, allyl-, butyl-, benzyl- and phenoxypropyl-malonate), the macrocyclic stambomycin analogues lacking C-28 hydroxylation (the ‘deoxy’ forms, **1, 3**, **5, 7** and **10**) were the main or even sole products (*i*.*e*. no ethyl-stambomycin was detected), accounting for 55.3−99.5% of the total yields based on their relative abundance in the mass spectra. In contrast, 6-bromohexyl-stambomycin (**11**) is the sole analogue observed in the 6-bromohexyl feeding experiments. This observation suggests that SamR0478 strongly prefers its native substrates or at minimum closely similar chains (**Figure 1**). In contrast, in no case were aglycone analogues detected in the fed extracts (data no shown), showing that the post-PKS glycosylase SamR0481, exhibits broad substrate tolerance, at least towards substrates of comparable size to the parental stambomycins^43^ (**Figure 1**).

Unexpectedly, for all fed extracts with the exception of thiophene-malonic acid and thienylmethyl-malonic acid for which there was no change, peaks corresponding to stambomycins A/B and C/D were always present, and at yields 2−8 times higher than from the unsupplemented ATCC/OE484/Δ483 mutasynthesis mutant (**Table 1** and **Figure S3**). To explain this surprising observation, we propose that the fed malonic acid derivatives somehow induce expression of the identified SamR0483 homologues (**Table S1**), which can then more substantially compensate for the loss of SamR0483. However, this mechanism remains to be verified directly by targeted inactivation.

### Quantitative analysis of the stambomycin derivatives produced by the mutasynthesis mutants ATCC/OE484/Δ483 and ATCC/OE484/Δ483/MatB_cinna

We next considered whether the lack of production of stambomycin derivatives in the presence of isopropyl-, phenylpropyl-, thiophene-, thienylmethyl-, and propargyl-malonic acid, might have resulted from intolerance to the modified polyketide chains by modules downstream of AT_12_. As such a mechanism could have resulted in release of stalled intermediates, we analyzed the extracts for the presence of intermediates corresponding to the products of modules 12 and 13. However, no convincing signal was detected (data not shown). Furthermore, we did not detect any of these predicted shunt products in extracts from feeding experiments where the stambomycin analogues were obtained, consistent with the idea that the modules downstream of module 12 are broadly tolerant to alternative chains. In light of this evidence, we hypothesized that the major blocks to incorporation of diacids were failure to efficiently activate the fed substrates to their CoA forms by MatB_Sa, or the inability of AT_12_ to recognize the modified malonates.

To directly address the first possibility, we introduced promiscuous MatB_cinna^45^ into mutant ATCC/OE484/Δ483, resulting in mutant ATCC/OE484/Δ483/MatB_cinna. Mutants ATCC/OE484/Δ483 and ATCC/OE484/Δ483/MatB_cinna were then fed with ethyl-, allyl-, butyl- and benzyl-malonic acids, followed by quantification of the resulting analogues produced by the two strains. For this, erythromycin (1 mM added during the work-up step) was used as an internal standard for the LC-HRMS analyses to control for any differences in extraction efficiency. The obtained data demonstrated that the presence of MatB_cinna improved the incorporation of ethyl-, allyl- and benzyl-malonic acids (**Figure 3a, b, d**), resulting in a nearly 2-fold increase relative to the parental strain which contains only MatB_Sa. Indeed, recent work has demonstrated that MatB_cinna efficiently converts ethyl-, allyl- and benzyl-malonic acid to their corresponding CoA forms^34^, which is consistent with the results obtained here. In contrast, levels of incorporation of butyl-malonic acid by the two mutants was essentially identical (88% vs 71%, **Figure 3c**), reflecting the fact that MatB_cinna is poorly active with butyl-diacid^34^.

**Figure 3.**
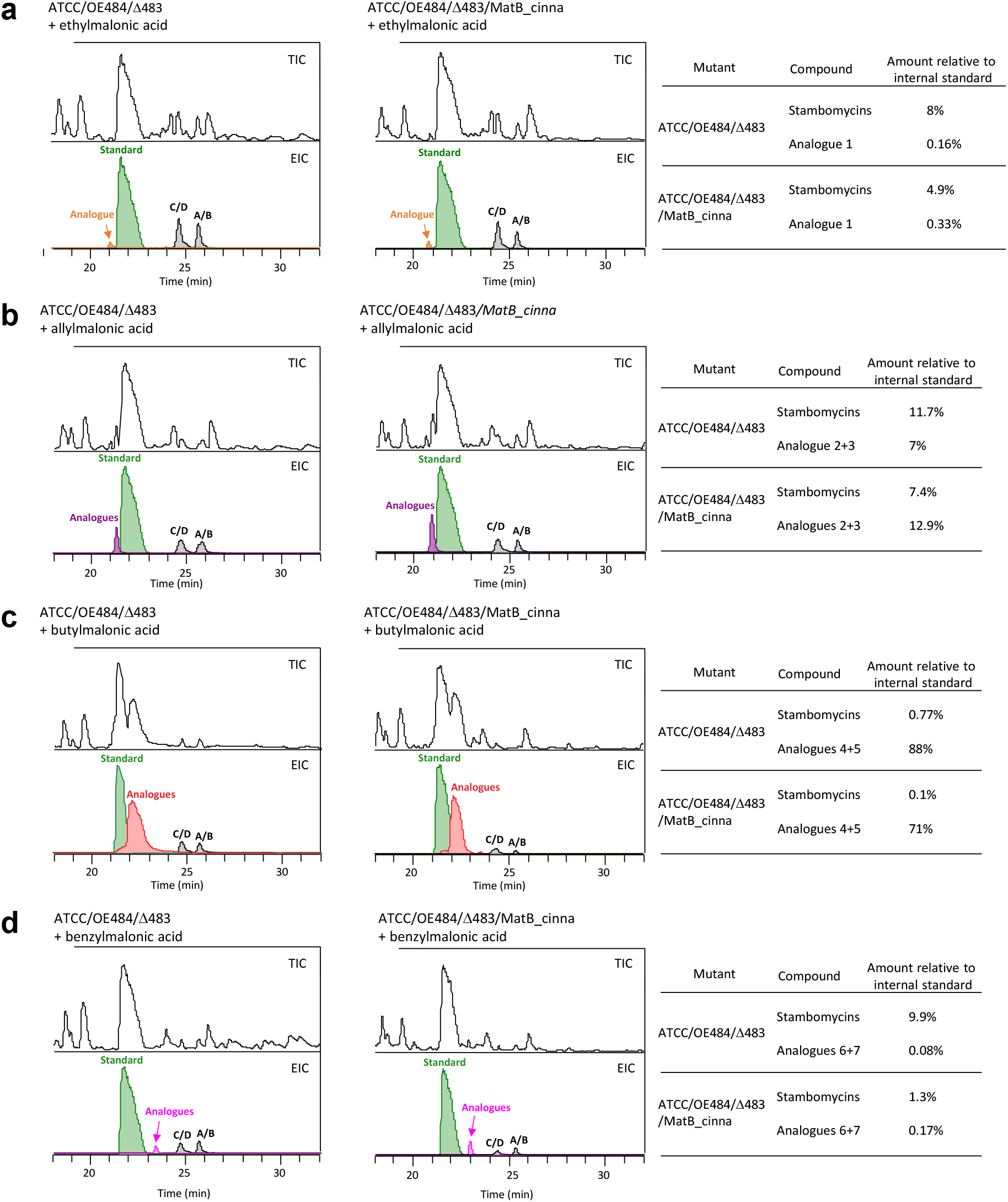
Quantitative analysis of the mutasynthesis mutants ATCC/OE484/Δ483 and ATCC/OE484/Δ483/ MatB_cinna by LC-HRMS using erythromycin as the internal standard. a) Comparison of the two mutants when supplied with ethylmalonic acid. b–d) Analysis following supplementation with allyl-, butyl- and benzylmalonic acids. TIC (total ion chromatogram) of the crude extract is shown in the top panel in each case. The amount of stambomycins (A–D) and stambomycins analogues relative to the internal standard erythromycin A were determined using the integrated area of EIC peaks (mean value of three individual fermentation results), with the peak area of the standard set to 100%.

With the aim of obtaining definitive structural proof for certain analogues, we therefore used ATCC/OE484/Δ483/ MatB_cinna to obtain allyl-incorporated stambomycin, and the original mutant ATCC/OE484/Δ483 to produce the butyl variant. This approach resulted in sufficient quantities of deoxy-allyl-stambomycin (**3**) (0.7 mg/L) to allow for its purification (the yields of allyl-stambomycin (**2**) were inadequate), while three compounds (22.5 mg/L combined yield of the two most abundant compounds) were purified from extracts of ATCC/OE484/Δ483 fed with butyl-malonic acid. The structures of these four compounds were elucidated by comprehensive NMR and HRMS analysis (**Figure S5**–**S29**), yielding data consistent with deoxy-allyl-stambomycin (**3**), deoxy-butyl-stambomycin (**5**) and butyl-stambomycin (**4**). Concerning the third butyl derivative, comparison of the obtained NMR data with that of butyl-stambomycin (**4**) revealed a methyl signal missing from the ^1^H NMR spectrum (δ_H_ 1.64) corresponding to the methyl at C-24, and the shift of an olefinic proton from δ_H_ 5.22 to δ_H_ 5.43. Additionally, coupling constants of 15.4 and 8.0 were observed for H-25 (**Figures S17**–**22**). These data are all consistent with a C-24-demethyl derivative of butyl stambomycin (**12**). Such a compound would arise from alternative incorporation of malonyl-CoA instead of methymalonyl-CoA by AT_13_, a reasonable hypothesis given the intrinsically broad specificity of AT_12_. In support of this hypothesis, reinspection of wild type extracts revealed a peak with a mass of 1334.8911 (C_70_H_128_NO_22_^+^; *m*/*z* [M+H]^+^ = 1334.8922, Δppm = −0.8), which matches that of a demethyl derivative of stambomycin E (**Figure S4b**) (note, it was not possible to conclusively identify analogous demethyl analogues of stambomycins AB, CD and F, as C-24-demethyl-stambomycins AB would have the same exact masses as stambomycins CD, C-24-demethyl-stambomycins CD, the same masses as stambomycin E, and C-24-demethyl-stambomycin F the same as stambomycins AB).

### Quantitative comparative analysis of the PDB and mutasynthesis approaches

As we observed production of the parental stambomycins in the mutasynthesis strains which is the classical outcome of PDB experiments, it was of interest to directly compare the relative efficiency of mutasynthesis to PDB. In contrast to mutasynthesis, PDB in *S. ambofaciens* relies on fed mono-acids, which are then activated by the SamR0482/SamR0483 pair to generate the corresponding extender unit CoA thioesters. Thus, to judge the relative efficacies of the two approaches, we supplemented strain ATCC/OE484 with 3-butenoic acid, butyric acid and phenyl propanoic acid (**Figure 4a**), and in parallel, mutant ATCC/OE484/Δ483 with the corresponding allyl-, butyl- and benzyl-malonic acids (**Figure 4b**). Erythromycin (1 mM) was again used as internal standard in the LC-HRMS analyses to permit accurate quantification. The obtained data showed that butyric acid was activated by the SamR0482/SamR0483-mediated pathway in strain ATCC/OE484, resulting in butyl-incorporated stambomycin analogues (**Figure 4d**), but that incorporation of 3-butenoic acid and benzyl-malonic acid failed (**Figure 4c, e**). Moreover, the yield of butyl-incorporated analogues from mutasynthesis with the ATCC/OE484/Δ483 strain relative to PDB based on ATCC/OE484 (88% vs 11%, **Figure 4d**), confirmed that mutasynthesis was the superior approach, despite the continued biosynthesis of the wild type stambomycins.

**Figure 4.**
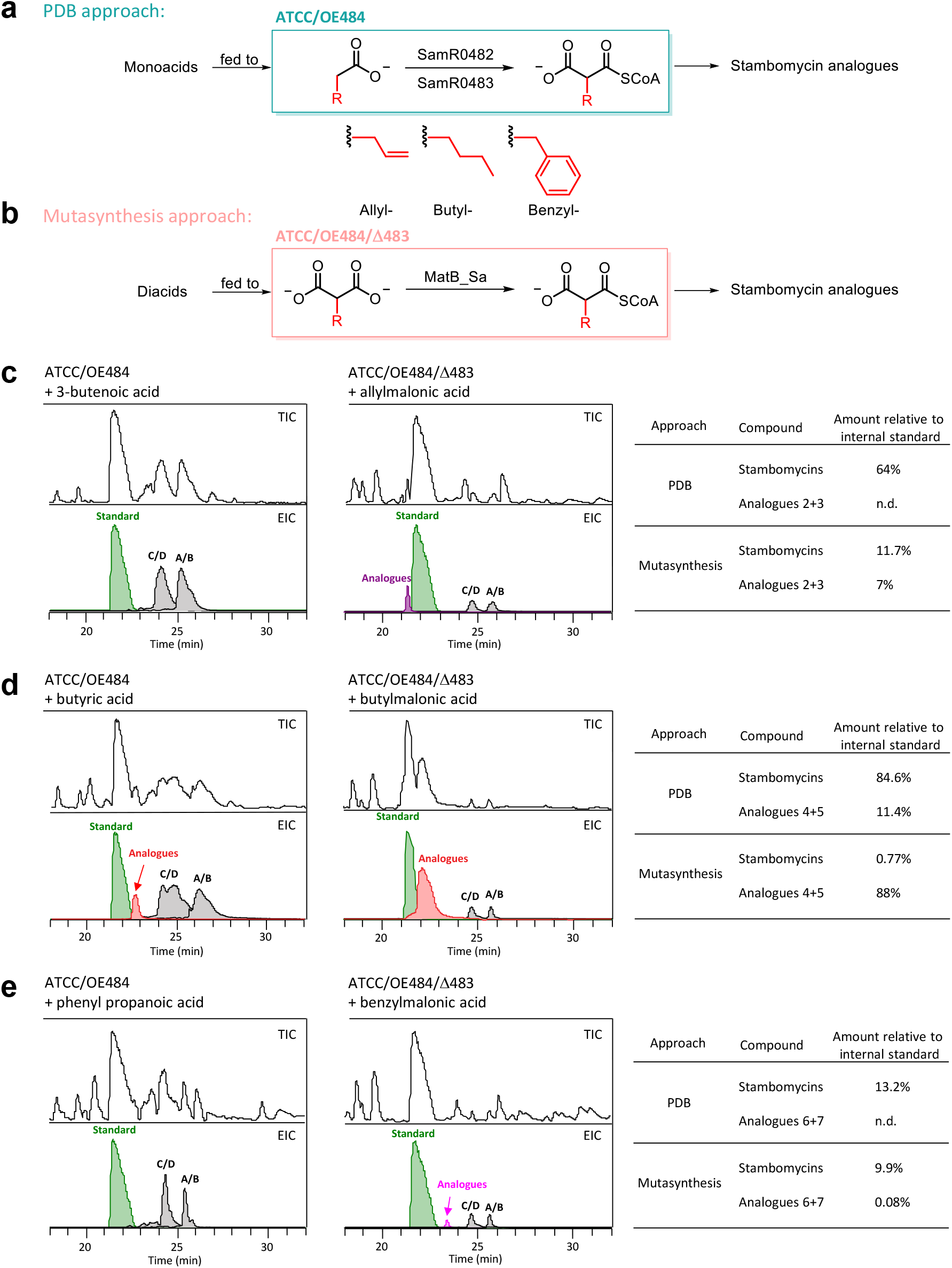
Quantitative analysis of PDB and mutasynthesis-mediated biosynthesis of stambomycin analogues. a) PDB: the culture medium is supplemented with monoacids, which are converted to extender units for AT_12_ by the two-enzyme pathway involving SamR0482 and SamR0483. b) Mutasynthesis: the culture medium is supplemented with diacids, which are converted to acyl-CoA extender units for AT_12_ by MatB_Sa. c−e) Quantitative analysis of crude extracts of the PDB and mutasynthesis experiments. The amount of stambomycins (A–D) and stambomycin analogues relative to the internal standard (erythromycin A) were determined via the integrated area of the EIC peaks (mean value of three individual fermentation results), with the peak area of the standard set to 100%.

### Biological activities of the stambomycin analogues

The stambomycins have moderate activity against Gram-positive bacteria but none against Gram-negative bacteria^42^. More interestingly they were also shown to possess antiproliferative activity towards a range of human cancer cells, with potency comparable to several clinically-used compounds ^42^. We therefore tested the bioactivity of the four purified stambomycin analogues (**3, 4, 5** and **12**). All four exhibited anti-bacterial activity, albeit variable, against both Gram-positive *Bacillus subtilis* and *Micrococcus luteus*, but not against the Gram-native *Escherichia coli* (**Figure S30**). The butyl- and deoxy-butyl-stambomycins (**4** and **5**) were the most active (with an efficiency similar to the original stambomycins), while the allyl form (**3**) showed only weak activity. Interestingly, the C-24-demethyl derivative of butyl-stambomycin (**12**) was less active relative to the (deoxy-)butyl-stambomycins.

Next, we tested the antiproliferative activities against two human cancer cell lines, U87-MG glioblastoma cells (brain cancer) and MDA-MTB-231 cells, which are commonly used to model late-stage breast cancer. MTT assays revealed significant and similar antiproliferative activities against the two lines for the butyl-stambomycin and its deoxy form (**Figure S31**) with IC_50_ values around 1 μM for both the U87 and the MDA-MB-231 cells (**Table S9**). These values are in the range of those observed for the parental stambomycins^42^ and are better than those observed for the clinical anticancer agent doxorubicin used as a control (**Table S9**). Cell counting further confirmed that the inhibition observed in the MTT assays was related to cell death (**Figure S31**). As observed with the anti-bacterial assays, the C-24-demethyl derivative (**12**) of butyl-stambomycin and the allyl analogue (**3**) were significantly less active against the two cell lines (**Table S9, Figure S31**).

## Discussion and Conclusions

In contrast to many PKS AT domains which exhibit strict substrate specificity^4^, the AT_12_ domain of the stambomycin biosynthetic pathway naturally recognizes at least six atypical malonate extender units, attesting to its innate broad substrate tolerance. This promiscuity in fact extends to all 12 downstream PKS modules and post-PKS enzymes, as the system generates a family of fully-elaborated stambomycins^42^. These observations motivated us to try to expand the library of available stambomycin C26 analogues^44^ for biological evaluation using a mutasynthesis approach. For this, we aimed to disable production of the native extender units by targeting the enzymes (SamR0482 and SamR0483) involved in their biosynthesis from medium-chain fatty acids^44^. By deletion of the individual genes, we demonstrated that SamR0483 was critical for stambomycin biosynthesis in the absence of additional growth modification, but that SamR0482 could be complemented by other enzymes encoded in the genome. We therefore carried out mutasynthesis using the SamR0483 inactivation mutant.

Supplementation of ATCC/OE484/Δ483 with suitable precursors yielded 6 novel series of stambomycin analogues at varying yields. Derivatives obtained in this way incorporated ethyl, butyl, allyl, benzyl, phenoxypropyl and 6-bromohexyl groups at C-26 instead of the native chains (**Table 1**) (initial evidence was also obtained for the C-26 octyl analogue). For each of the analogues except that bearing the 6-bromohexyl group which was fully modified, a major proportion of the compounds lacked the C-28 hydroxylation catalyzed by the P450 enzyme SamR0478. It is also notable that feeding of butylmalonic acid resulted in almost exclusive production of butyl-stambomycin relative to residual stambomycins A−D, whereas the native metabolites dominated in previous precursor-directed biosynthesis experiments^44^. Indeed, in our hands, the PDB approach only yielded a single novel derivative when a limited series of monoacid substrates was fed. Overall, successful production of this panel of analogues depended on the intrinsic substrate tolerance of three distinct enzymatic events in the pathway: the MatB_Sa-catalyzed activation of the diacids as their CoA thioesters, substrate selection and ACP loading by AT_12_, and recognition of the modified intermediates by the downstream PKS modules, the C-50 hydroxylase SamR0479^54^, and SamR0481-mediated mycaminosylation. It is also worth noting that producing comparable analogues using synthetic chemistry would require their total synthesis.

Nonetheless, the intrinsically broad specificity of AT_12_ and the overall PKS system did not translate into universal acceptance of all alternative building blocks. Intriguingly, despite the bulky nature of the (methyl)pentyl/(methyl)hexyl side chains of the native extender units, the non-native analogues showing the highest level of incorporation were significantly shorter (*i*.*e*. butyl and allyl). The efficient incorporation of butyl-malonic acid is also notable for a second reason, as it suggests that MatB_Sa has unusually high activity towards this substrate, which was not seen with MatB_cinna^34^ and MatB_Rt^41^ despite their intrinsic promiscuity. This hypothesis is also supported by the fact that expression of MatB_cinna alongside MatB_Sa did not improve incorporation of butyl-malonate by the strain. Concerning the substrates which were not accepted (isopropyl, phenylpropyl-, thiophene-, thienylmethyl- and propargylmalonic acid), it is not possible to conclusively identify the enzymatic bottleneck(s), as none of these substrates were directly evaluated in previous work with MatB_cinna^34^, but the absence of prematurely released chain extension intermediates points to either AT_12_ or MatB.

Our positive incorporation results notably contrast with recent data obtained from an engineered swap of AT_12_ into module 6 of the erythromycin (DEBS) PKS^12^. In this case, analysis *in vitro* with purified hybrid protein and a series of α-carboxyacyl-CoA substrates revealed essentially no activity towards ethyl- and allylmalonyl-CoA, comparable but low activity with propyl-, butyl-, pentyl-, hexylmalonyl-CoA, and a pronounced preference for isopentyl-CoA. This result taken alongside other obtained data was interpreted to show that atypical AT domains can incorporate extender units which are equal in size or slightly larger than their native substrates^12^. Indeed, AT_12_ was predicted by AlphaFold2^12, 55^ to possess one of the largest active site cavity volumes among the evaluated ATs, in accord with the high steric demands of its native substrates.

At this stage it is difficult to reconcile the two sets of data, particularly as incorporation of butyl-malonyl-CoA in our hands resulted in mature analogues at yields comparable to that of the parental stambomycins (20–30 mg/L)^42^. Either the AT in its native context exhibits a more relaxed specificity, or the specificity of AT_12_ was unchanged, but the tolerance of the other DEBS module 6 domains was negatively impacted by introduction of this heterologous AT (in the presence of alternative non-native ATs, DEBS module 6 accepted both allyl- and butylmalonyl-CoA with good efficiency^12^). Another possible explanation, however, is that insertion of AT_12_ into the module caused structural perturbation, which impacted turnover. Indeed, the AT swapped modules which gave the lowest purification yields relative to wild type also exhibited the most restricted substrate specificities^12^. In any case, the cumulative results point to a complex interplay between the AT and the remaining domains of the PKS in determining the efficacy with which alternative building blocks can be incorporated into engineered polyketides.

Our observation that the butyl (C_4_) congeners (**4** and **5**) retained good activity relative to the native metabolites (linear and branched C_5_ and C_6_ side chains) demonstrates a certain tolerance to the chain length and functionality at the C-26 position. However, the allyl analogue (C_3_) (**3**) was markedly less potent, an effect which may be due to the further reduced length, or the presence of the double bond. Our data also confirm an earlier observation made with the parental stambomycins^46^ that the hydroxylation at C-28 has no significant effect on the biological activities of the analogues. Interestingly, the methyl group at C-24 appears to contribute significantly to both the antibacterial and antiproliferative properties of the derivatives, as its absence (*i*.*e*. **12**) results in a sharp decrease in bioactivity. Potential reasons for this drop include reduced lipophilicity and thus biomembrane solubility, or the lack of an important conformational constraint on the macrolides. We note that the effect of methylation on biological interactions has strong precedence in the literature^56^.

Finally, given the observed specificity differences between MatB_Sa and MatB_cinna, it would be of interest to attempt to further broaden the substrate tolerance of MatB_Sa by structure-guided mutagenesis. Notably, an engineered version of MatB_Rt (T207G/M306I) showed improved catalytic activity toward ethyl-, propargyl- and allyl-malonic acids, as well some tolerance towards isopropyl-, butyl-, phenyl-, and azido-malonic acids^41^. The two residues corresponding to T207 and M306 in MatB_Sa are V190 and M293 (**Figure S3**), which could be modified in attempts to similarly broaden the substrate scope of MatB_Sa. In parallel, it would be worth targeting for inactivation the identified homologues of SamR0483 in order to reduce the background synthesis of the parental stambomycins, potentially boosting the yields of the novel derivatives. We also envision that an analogous mutasynthesis strategy could be applied to other-diversity oriented pathways but with supplementation using one of the native precursors – redirecting the pathway to production of a single natural compound which should simplify purification and subsequent structure elucidation^57^. Together, such approaches should further expand the enzymatic toolbox available for efficient AT-based polyketide analogue generation.

## Author contributions

K. J. W. and B. A. designed and supervised the project, with initial assistance from C.J. K. J. W. and B. A. obtained the funding at the U.L. and H.B.B. at the MPI. L. S. constructed and fermented (together with L. H., Y. S., and C. J.) the mutasynthesis strains, and analysed and interpreted the LC-MS data. Y. S. synthesized propargyl-malonic acid and interpreted the NMR data. P. G., Y-N. S., and H. B. B purified compounds and performed the NMR measurements. A. C. and S. P. carried out the biological testing. L. S., B. A. and K. J. W. wrote the manuscript, with contributions from all authors.

## Conflicts of interest

There are no conflicts to declare.

## Acknowledgements

We acknowledge financial support from the Université de Lorraine, the Centre National de la Recherche Scientifique (CNRS), and the IMPACT Biomolecules project of the Lorraine Université d’Excellence (LUE) (Investissements d’avenir − ANR 15-004 to L.S., Y.S., C.J., B.A., and K.J.W.). Work in the Bode lab was supported by an ERC Advanced Grant (835108) and the Max-Planck-Society. We are grateful to Prof. Yi Yu (Wuhan University, China) for the chemical synthesis of 6-bromohexyl-malonic acid and Prof. Frank Schulz (Ruhr-Universität Bochum, Germany) for the plasmid encoding MatB_cinna. We are grateful for access to the PASM platform of the Université de Lorraine, and the technical assistance provided by C. Paris.

## Notes

### Competing Interest Statement

The authors have declared no competing interest.

