## Supplementary material for "Exploiting the inherent promiscuity of the acyl transferase of the stambomycin polyketide synthase for the mutasynthesis of analogues"

[h] Senckenberg Gesellschaft für Naturforschung, 60325 Frankfurt am Main, Germany.

[i] Center for Synthetic Microbiology (SYNMIKRO), University of Marburg, 35043 Marburg, Germany

* Corresponding authors

### Materials and Methods

#### Materials

All reagents and chemicals including the fed mono- and diacids were obtained from Sigma-Aldrich. Oligonucleotide primers were synthesized by Eurogentec (France). The propargyl-malonic acid and 6-bromohexyl-malonic acid were chemically synthesized as described in reference[1].

#### Strains and media

Unless otherwise specified, all *Escherichia coli* strains were cultured in LB medium (yeast extract 10 g, tryptone 5 g, NaCl 10 g, distilled water up to 1 L, pH 7.0) or on LB agar plates (LB medium supplemented with 20 g/L agar) at 37 °C. *Streptomyces ambofaciens* ATCC23877 and its derived mutants were grown in TSB (TSB powder 30 g (tryptone 17 g, soy 3 g, NaCl 5 g, $K_2HPO_4$ 2.5 g, glucose 2.5 g), distilled water up to 1 L, pH 7.3) or on TSA plates (TSB medium supplemented with 20 g/L agar), and sporulated on SFM agar plates (NutriSoy flour 20 g, D-mannitol 20 g, agar 20 g, tap water up to 1 L) at 30 °C. All strains were maintained in 20% (v/v) glycerol in 2 mL Eppendorf tubes and stored at −80 °C. For fermentation of *S. ambofaciens* ATCC23877 and its mutants, spores were streaked on TSA with appropriate antibiotics (apramycin 50 μg $mL^{-1}$, kanamycin 50 μg $mL^{-1}$, spectinomycin 50 μg $mL^{-1}$) and after incubation for 48 h at 30 °C, a loop of mycelium was used to inoculate 7 mL of MP5 medium (yeast extract 7 g, NaCl 5 g, $NaNO_3$ 1 g, glycerol 36 mL, MOPS 20.9 g, distilled water up to 1 L, pH 7.4) supplemented with selective antibiotics and sterile glass beads, followed by incubation at 200 rpm and 30 °C for 24–48 h. Finally, the seed culture was centrifuged (3500 *g*) and resuspended in 2 mL fresh MP5 before being inoculated into 50 mL MP5 medium in a 250 mL Erlenmeyer flask, and cultivated at 200 rpm and 30 °C for 4 days. Monoacids and diacids were fed to the media at a final concentration of 10 mM after inoculation of seed culture at 24 h incubation.

#### PCR-targeting based genetic engineering to generate the mutant ATCC/OE484/Δ483

To render the BAC BAB19ZF4 proficient for selection following conjugation, its chloramphenicol resistance gene was replaced , by a spectinomycin resistance gene cassette sourced from pIJ778[2] using a PCR-targeting approach, resulting in BAC3 (**Table S3**). In order to obtain an in-frame deletion of the *samR0483* gene*,* the primer pair D483_For/D483_Rev (**Table S4**) was designed to PCR amplify the "*FRT+aac(3)IV+oriT+FRT*" disruption cassette from pIJ773[2], affording PCR fragment K7Δ483. The PCR fragment was then electro-transformed into *E. coli* BW25113/pKD20/BAC3, giving rise to the mutant BAC BAC3_K7Δ483. Subsequently, BAC3_K7Δ483 was introduced into *E. coli* ET12567/pUZ8002 and then *S. ambofaciens* ATCC23877 *via* intergeneric conjugation. The resulting exconjugants (ATCC/Δ483_*aac(3)IV+oriT*) were selected for apramycin resistance and spectinomycin sensitivity, attesting to successful double crossover. The correct mutations were confirmed by PCR using primer pair F483_For/F483_Rev and sequencing. Thereafter, the cassette "*aac(3)IV+oriT*" was excised by the FLP recombinase encoded by plasmid pUWL-*oriT-flp* as described previously[3], leaving an 81 bp scar sequence (mutant ATCC/Δ483). Successful removal of the cassette was verified by PCR using primer pair F483_For/F483_Rev and DNA sequencing. Finally, the LAL regulator overexpression plasmid pOE484[4] and the parental vector pIB139 which was used as a control, were transferred into mutant ATCC/Δ483, giving rise to the mutants ATCC/OE484/Δ483 and ATCC/pIB139/Δ483, respectively (**Table S2**). (Note, mutant ATCC/OE484/Δ482 was generated in previous work[4]).

#### Complementation of mutant ATCC/OE484/Δ483

The gene *samR0483* and the constitutive promoter *ermEp** (including an RBS) were PCR amplified using primers Comp483_For/Comp483_Rev and ermEp_For/ermEp_Rev (**Table S4**) using ATCC genomic DNA and pIB139 as template, respectively. Two PCR fragments were fused into one fragment (*ermEp*+RBS+samR483*) by overlap extension PCR, prior to cloning into the vector pCR™-Blunt. The fragment was then excised from pCR™-Blunt by FD *Nhe*I and FD *Spe*I digestion, and then cloned into pre-digested (FD *Nhe*I and FD *Spe*I) and dephosphorylated (FastAP) vector pOSV809, giving rise to the construct pOSV809-*ermEp*-samR483.* The vector was then transformed into *E. coli* S17-1 and introduced into mutants ATCC/Δ483/OE484 and ATCC/Δ483/pIB139 via intergeneric conjugation. Kanamycin and apramycin resistant conjugants were selected and verified by PCR with primers pOSV-For/φBT1-attB-Rev and pOSV-Rev/φBT1-attB-For (**Table S4**).

**Construction of mutant ATCC/OE484/Δ483/MatB_cinna**

The gene *matB_cinna* was cloned into an integrative and spectinomycin resistant vector by the group of Prof. Frank Schulz (Ruhr-Universität Bochum, Germany), resulting in the plasmid pRT801_lacZ_PermE_MatB_cinna. The plasmid was then transformed into *E. coli* S17-1 and introduced into mutant ATCC/Δ483/OE484 via intergeneric conjugation. Spectinomycin and apramycin resistant conjugants were selected and verified by PCR with primers pOSV-For/φBT1-attB-Rev and pOSV-Rev/φBT1-attB-For (**Table S4**).

**LC-ESI-HRMS analysis of fermentation metabolites**

The fermentation broth was centrifuged at 4,000 *g* for 10 min. The parental stambomycins and their derivatives were then extracted from the mycelia by first resuspending the cells in 40 mL distilled water, followed by centrifugation (4,000 *g*, 10 min, repeated 3×) to remove the water-soluble components. After decanting the water, the cell pellets were weighed and extracted with MeOH by shaking at 150 rpm for 2 h at room temperature. Thereafter, the MeOH extracts were filtered to remove the cell debris, followed by rotary evaporation to dryness. The obtained extracts were dissolved in MeOH, whose volume was determined according to the initial weight of the mycelia (70 µL MeOH to 1 g of cell pellet). The resulting mycelial extracts were then centrifuged at 16,000 *g* at 4 °C for 20 min and analysed in heated positive electrospray (HESI+) mode by HPLC-HRMS on either a Thermo Scientific Orbitrap LTQXL or an Orbitrap ID-X Tribrid Mass Spectrometer using an Alltima™ C18 column (2.1 × 150 mm, 5 µm particle size). Separation was carried out with Milli-Q water containing 0.1% formic acid (A) and acetonitrile containing 0.1% formic acid (B) using the following elution profile: 0–48 min, linear gradient 5–95% solvent B; 48–54 min, constant 95% solvent B; 54–60 min, constant 5% solvent B. Mass spectrometry operating parameters were: spray voltage, 5 kV; source gases were set respectively for sheath gas, auxiliary gas and sweep gas at 30, 10, and 10 arbitrary units min$^{-1}$; capillary temperature, 275 °C; capillary voltage, 4 V; tube lens, split lens and front lens voltages 155, −28, and −6 V, respectively. Due to the much lower sensitivity of the Orbitrap LTQXL relative to the Orbitrap ID-X Tribrid as described previously[5], we introduced a 10× correction factor to the yields determined using the Orbitrap LTQXL. Quantification was achieved by adding erythromycin (1 mM) as internal standard during the extraction process. Alternatively, the yield of stambomycins from parental and mutasynthesis strains were determined using the stambomycin standard curve generated previoulsy[5], based on the integrated areas of EIC (extracted ion chromatogram) peaks.

NMR spectra were acquired on Bruker AV 500 and AV600 (only used for deoxy-allyl-stambomycin measurements) spectrometers, using the residual solvent peaks as reference. The spectra were processed using MestReNova 15.0.1 software. Coupling constants (*J*) are given in Hz.

**Isolation and purification of mutasynthetic stambomycin analogues**

In order to isolate allyl- and butyl-incorporated stambomycin analogues, 85 × 50 mL flasks of fermentation culture for ATCC/Δ483/OE484/MatB_cinna and 50 × 50 mL flasks culture for ATCC/Δ483/OE484 were cultivated in the presence of 10 mM fed allyl- and butylmalonic acids, respectively. The crude allyl-fed extract (18 mL in MeOH) and butyl-fed extract (12 mL in MeOH) were subjected to Agilent preparative HPLC system and a Waters Xbridge BEH C18 column (19 × 250 mm). The two extracts were separated by using acetonitrile/$H_2O$ containing 0.1% formic acid with the elution profile of 0–2 min 10% acetonitrile, then 2–27 min linear gradient 5–95% acetonitrile at the flow rate of 20 mL $min^{-1}$ to yield four purified compounds deoxy-butyl-stambomycin (36.3 mg), butyl-stambomycin (11.6 mg), C-24-demethyl-stambomycin (3.1 mg), and deoxy-allyl-stambomycin (1.8 mg). Details for the HPLC separation are also summarized in **Table S8**.

**Antibacterial and antiproliferative tests**

The antibacterial activities of the four purified stambomycin analogues were analysed by loading 1 µL of the compounds at 10 mM and at various dilutions (from 1:2 to 1:20) onto a lawn of the Gram-positive bacteria *Bacillus subtilis* ATCC6633 and *Micrococcus luteus*, or the Gram-negative bacterium *Escherichia coli* DH5α. One microliter of DMSO, the solvent used to resuspend the purified analogues, was used as a control. The plates were incubated overnight at 30 °C (*B. subtilis*) or 37° C (*M. luteus* and *E. coli*). For the antiproliferative activities on tumour cells, MTT (3-[4,5-dimethylthiazol-2-yl]-2,5-diphenyltetrazolium bromide) assays were carried out to measure the effects of the stambomycin analogues on the metabolic activity of two cancer cell lines, U87-MG glioblastoma cells (brain cancer) and MDA-MTB-231 cells (breast cancer). Briefly, U87-MG and MDA-MTB-231 cells (200 µL at $1.5.10^4$ cells/mL) were seeded in 96-well plate. Drug treatments started 48h later. After 48 h, cells were incubated with 0.625 mg/ml MTT for 3 h and the absorbance was measured at 560 nm using a microplate reader. To complete, cell viability was assessed by cell counting (number of cells/mL) using TC20 automated cell counter (Bio-Rad). All the assays were performed in triplicate in three independent experiments and the concentrations that induced 50% of cytotoxicity (IC50) were calculated using Prism 5.00 software.

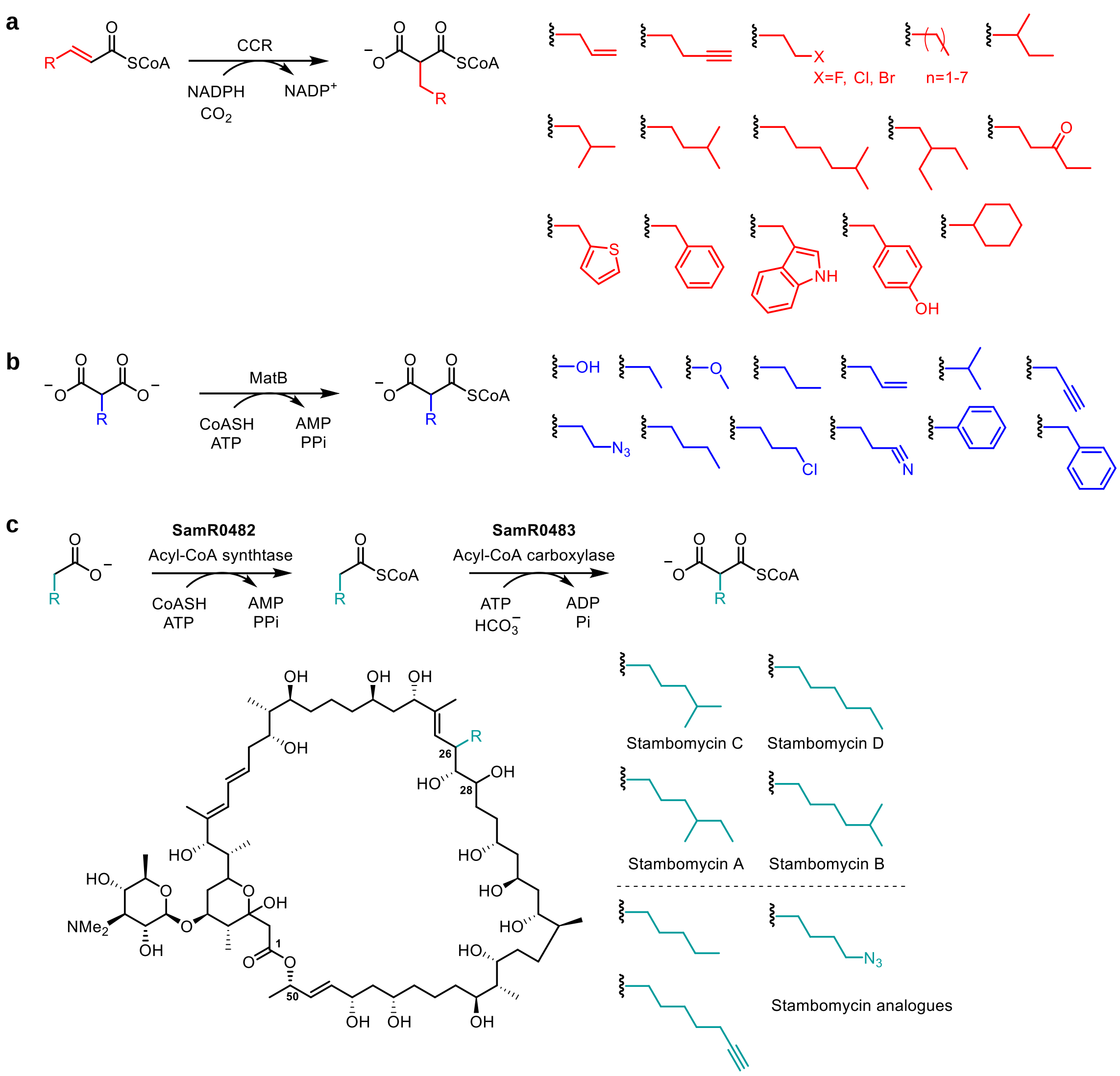


**Figure S1.** Biosynthesis of CoA-linked polyketide synthetase extender units. **a**) CCR enzymes (e.g. AntE[6], SalG[7] and SpnE[8], etc.) and their engineered variants catalyse the reductive carboxylation of CoA-linked α,β-unsaturated acyl-CoA via utilization of one reducing equivalent of NADPH, to afford malonyl-CoA analogous. **b**) MatB (e.g. MatB_Sc[9] from *S. coelicolor*, MatB_cinna[1] from *S. cinnamonesis* and MatB_Rt[10] from *Rhizobium trifolii*, etc.) and derived mutants are capable of generating the CoA derivatives of a broad range of malonate analogues, via an ATP-dependent reaction. **c**) The SamR0482/SamR0483 mediated pathway[11] and the structures of native and PDB-derived novel stambomycins. The two-enzyme mediated pathway affords six atypical malonyl-CoA extender units, by CoA derivatization followed by direct carboxylation of medium chain acyl-carboxylic acids sourced from the cellular pool. It can also act on exogenously-fed carboxylic acids, giving rise to the corresponding stambomycin analogues.

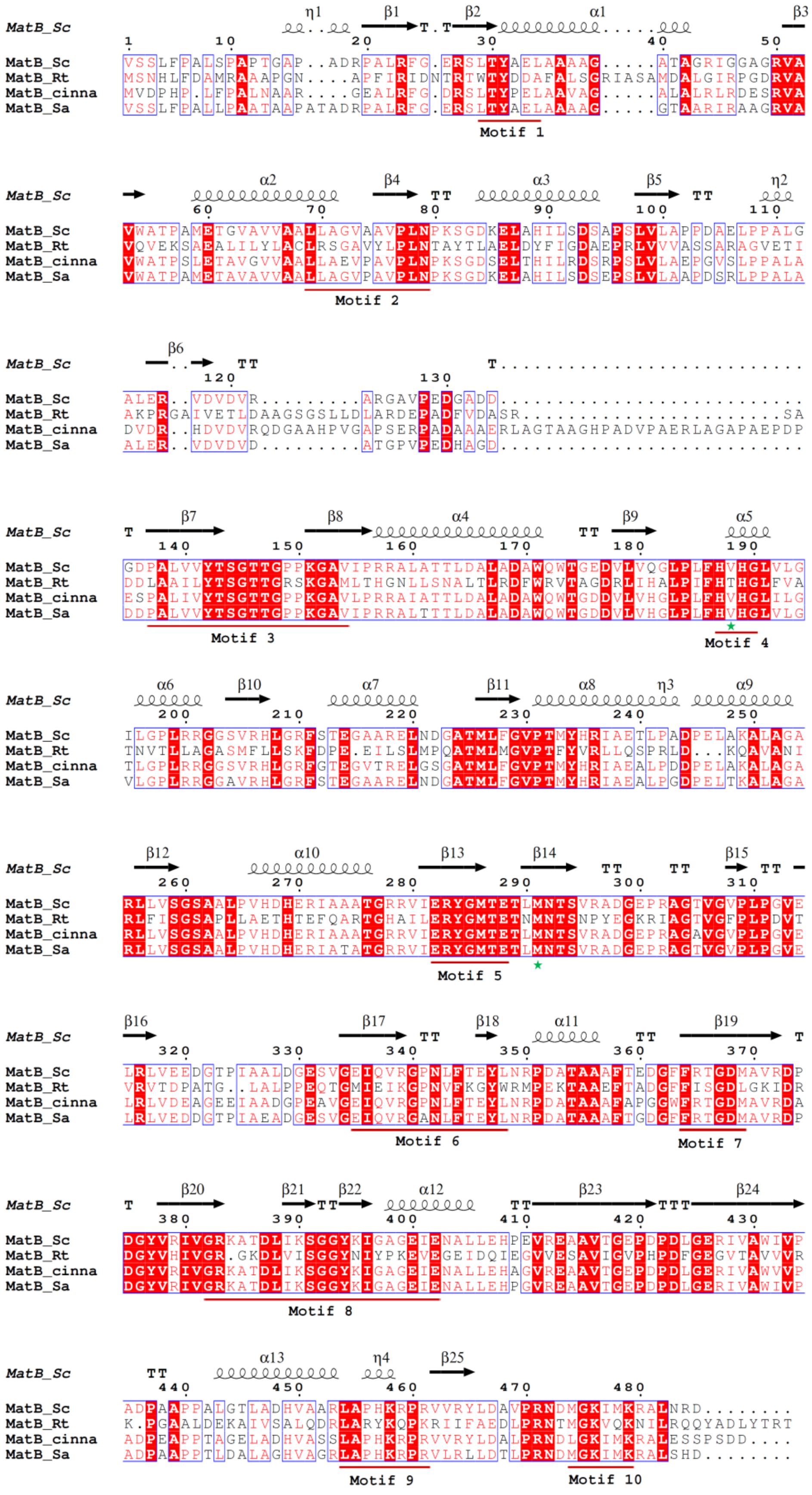
MatB_Sc
MatB_Rt
MatB_cinna
MatB_Sa
Motif 1
Motif 2
Motif 3
Motif 4
Motif 5
Motif 6
Motif 7
Motif 8
Motif 9
Motif 10

**Figure S2.** Sequence alignment of MatB-type enzymes. The secondary structure of MatB_Sc (PDB 3NYQ:A) is shown at the top. Residues which are strictly conserved are indicated in white on a red background, while relatively conserved residues are indicated in red. The red underlines represent conserved core motifs for adenylate-forming enzymes, of which motifs 4 and 5 are known to bind the carboxylated substrates and stabilize the formation of the adenylate intermediates[9]. The green stars refer to two hydrophobic amino acids mutated in MatB_Rt[10], which on the basis of the elucidated structure of MatB_Sc are likely to influence the promiscuity of MatB-type enzymes for α-substituted malonate derivatives. MatB_Sa, WP_053130319, MatB from *S. ambofaciens*; MatB_Sc, 3NYQ:A, MatB from *S. coelicolor*; MatB_cinna, MatB from *S. cinnamonensis* (no available accession number but the amino acid sequence was published previously[12]); MatB_Rt, AAC83455, MatB from *R. trifolii*. MatB_Sa exhibits 90%, 75% and 39% amino acid sequence identity respectively to MatB_Sc, MatB_cinna and MatB_Rt.

**a**

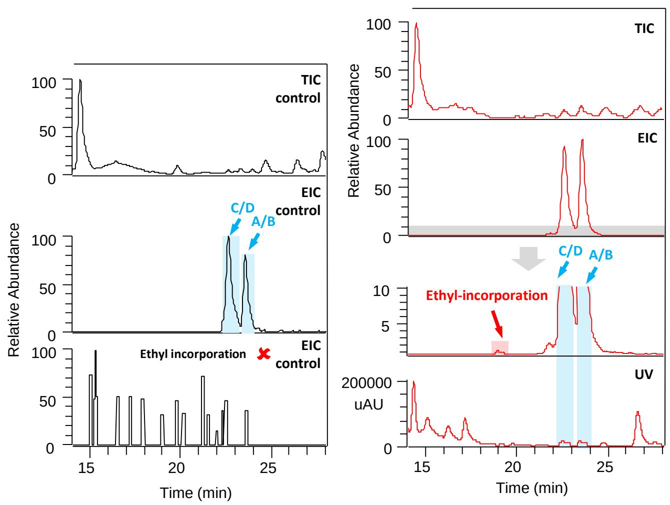


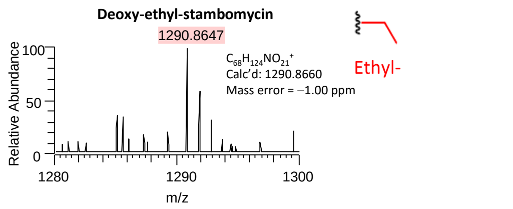


| Mutant ATCC/OE484/Δ483 | Compounds | Integrated area of EIC peaks |
|---|---|---|
| Control (unsupplemented) | Stambomycin C/D + A/B | 17501475 (100%) |
| Supplemented with ethylmalonic acid | Stambomycin C/D + A/B | 149221532 (853%) |
| | Ethyl-stambomycin | n.d. |
| | Deoxy-ethyl-stambomycin (**1**) | 151190 (0.86%) |

**b**

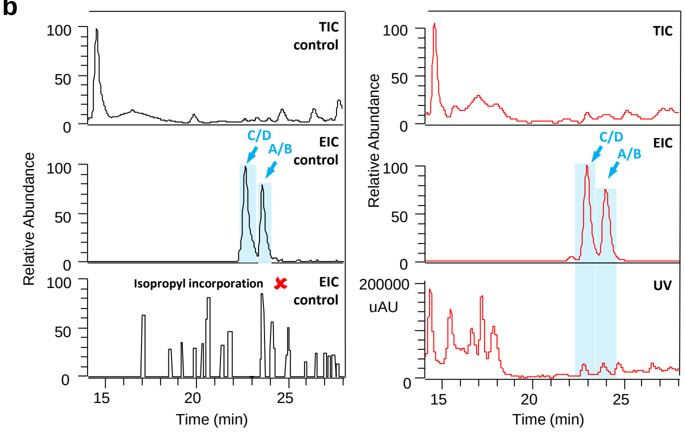


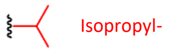


| Mutant ATCC/OE484/Δ483 | Compounds | Integrated area of EIC peaks |
|---|---|---|
| Control (unsupplemented) | Stambomycin C/D + A/B | 17501475 (100%) |
| Supplemented with isopropyl-malonic acid | Stambomycin C/D + A/B | 63227152 (361%) |
| | Isopropyl-stambomycin | n.d. |
| | Deoxy-isopropyl-stambomycin | n.d. |

**c**

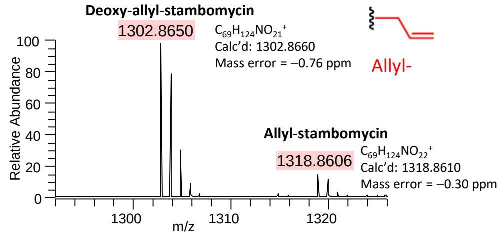


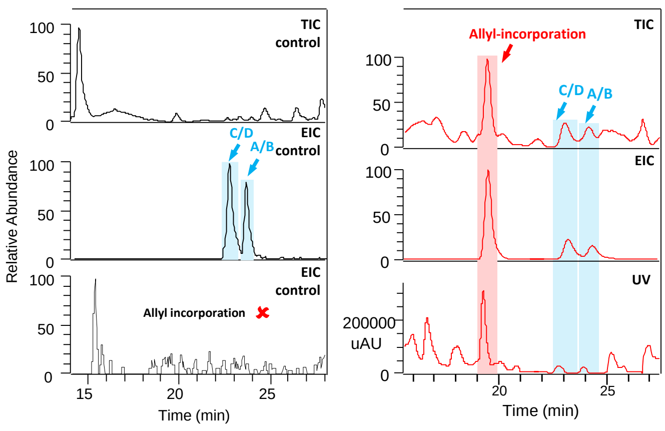


| Mutant ATCC/OE484/Δ483 | Compounds | Integrated area of EIC peaks |
|---|---|---|
| Control (unsupplemented) | Stambomycin C/D + A/B | 17501475 (100%) |
| Supplemented with allyl-malonic acid | Stambomycin C/D + A/B | 33027066 (189%) |
| | Allyl-stambomycin (**2**) | 359513 (2.05%) |
| | Deoxy-allyl-stambomycin (**3**) | 65873199 (378%) |

**d**

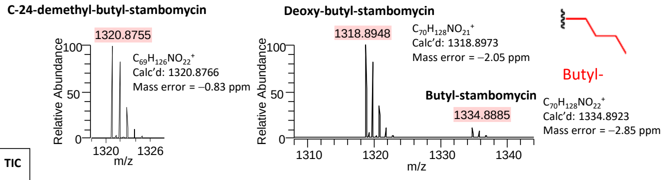


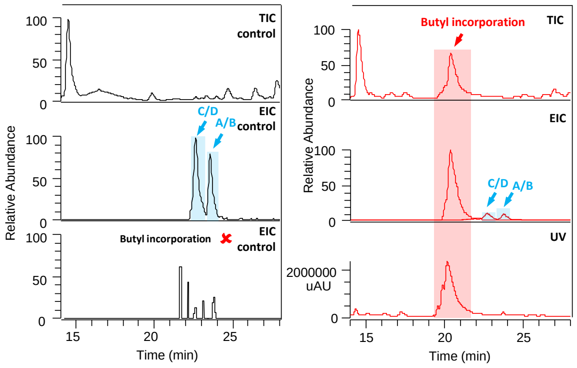


| Mutant ATCC/OE484/Δ483 | Compounds | Integrated area of EIC peaks |
|---|---|---|
| Control (unsupplemented) | Stambomycin C/D + A/B | 17501475 (100%) |
| Supplemented with butyl-malonic acid | Stambomycin C/D + A/B | 33568599 (192%) |
| | C-24-demethyl-butyl-stambomycin (**12**) | 78628078 (449%) |
| | Butyl-stambomycin (**4**) | 757196925 (4326%) |
| | Deoxy-butyl-stambomycin (**5**) | 3608738285 (20619%) |

**e**

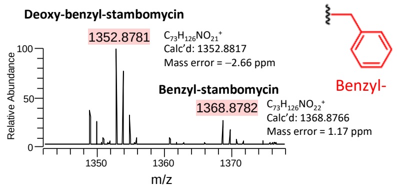


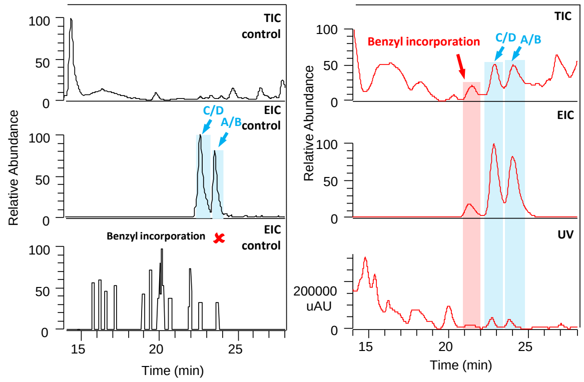


| Mutant ATCC/OE484/Δ483 | Compounds | Integrated area of EIC peaks |
|---|---|---|
| Control (unsupplemented) | Stambomycin C/D + A/B | 17501475 (100%) |
| Supplemented with benzyl-malonic acid | Stambomycin C/D + A/B | 83601192 (478%) |
| | Benzyl-stambomycin (**6**) | 4814150 (27.5%) |
| | Deoxy-benzyl-stambomycin (**7**) | 9595426 (54.8%) |

**f**

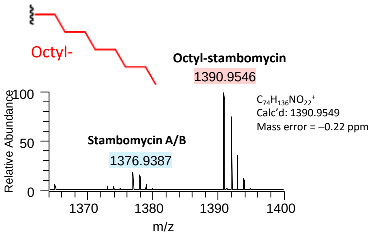


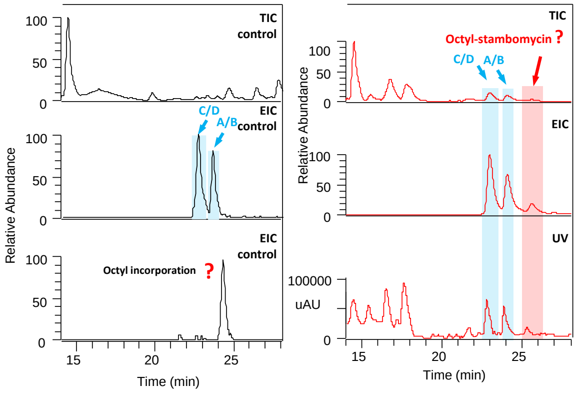


| Mutant ATCC/OE484/Δ483 | Compounds | Integrated area of EIC peaks |
|---|---|---|
| Control (unsupplemented) | Stambomycin C/D + A/B | 17501475 (100%) |
| Supplemented with octyl-malonic acid | Stambomycin C/D + A/B | 147914000 (845%) |
| | Octyl-stambomycin (**8**) | 13572281 (77.5%) |
| | Deoxy-octyl-stambomycin | n.d. |

**g**

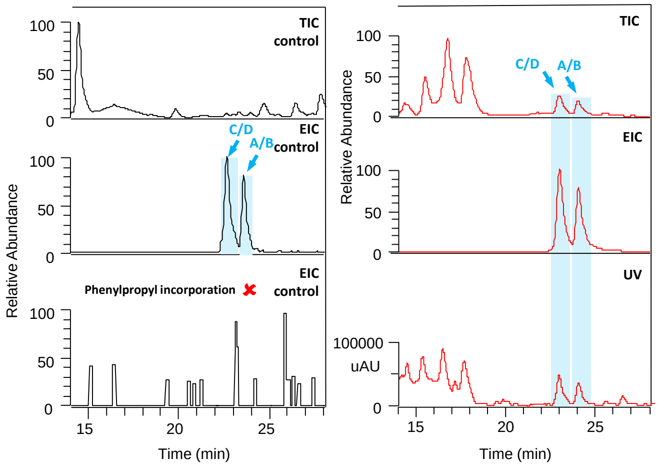


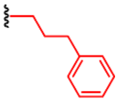


| Mutant ATCC/OE484/Δ483 | Compounds | Integrated area of EIC peaks |
|---|---|---|
| Control (unsupplemented) | Stambomycin C/D + A/B | 17501475 (100%) |
| Supplemented with phenylpropyl-malonic acid | Stambomycin C/D + A/B | 103429282 (591%) |
| | Phenylpropyl-stambomycin | n.d. |
| | Deoxy-phenylpropyl-stambomycin | n.d. |

**h**

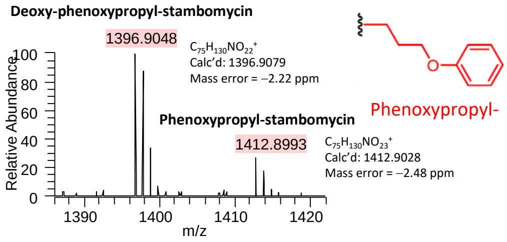


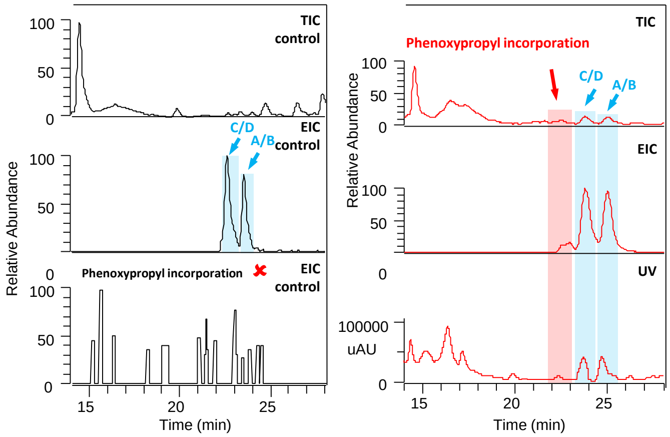


| Mutant ATCC/OE484/Δ483 | Compounds | Integrated area of EIC peaks |
|---|---|---|
| Control (unsupplemented) | Stambomycin C/D + A/B | 17501475 (100%) |
| Supplemented with phenoxypropyl-malonic acid | Stambomycin C/D + A/B | 40699658 (233%) |
| | Phenoxypropyl-stambomycin **(9)** | 1916650 (10.9%) |
| | Deoxy-phenoxypropyl-stambomycin **(10)** | 2366663 (13.5%) |

**i**

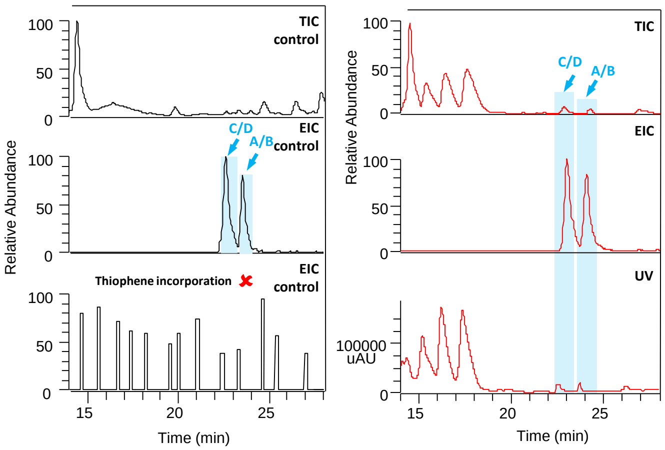


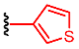


| Mutant ATCC/OE484/Δ483 | Compounds | Integrated area of EIC peaks |
|---|---|---|
| Control (unsupplemented) | Stambomycin C/D + A/B | 17501475 (100%) |
| Supplemented with thiophene-malonic acid | Stambomycin C/D + A/B | 14692173 (83.9%) |
| | Thiophene-stambomycin | n.d. |
| | Deoxy-thiophene-stambomycin | n.d. |

**j**

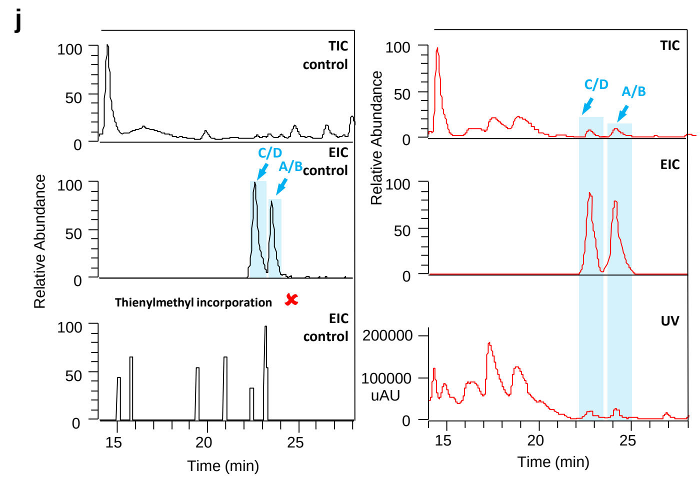


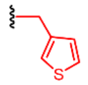

Thienylmethyl-

| Mutant ATCC/OE484/Δ483 | Compounds | Integrated area of EIC peaks |
|---|---|---|
| Control (unsupplemented) | Stambomycin C/D + A/B | 17501475 (100%) |
| Supplemented with thienylmethyl-malonic acid | Stambomycin C/D + A/B | 17404928 (99.4%) |
| | Thienylmethyl-stambomycin | n.d. |
| | Dexoy-thienylmethyl-stambomycin | n.d. |

**k**

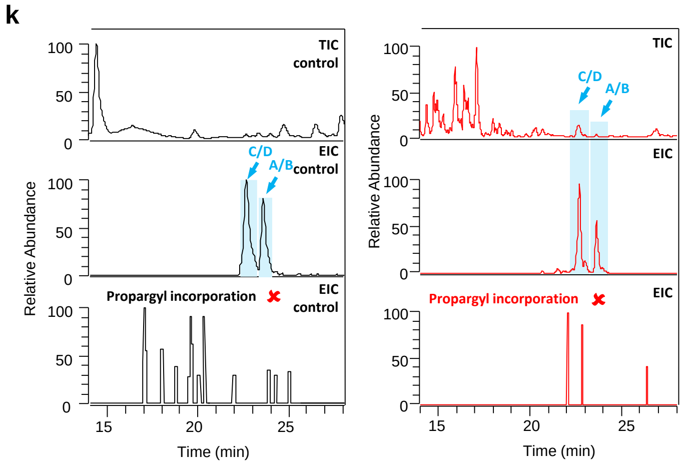


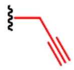

Propargyl-

| Mutant ATCC/OE484/Δ483 | Compounds | Integrated area of EIC peaks |
|---|---|---|
| Control (unsupplemented) | Stambomycin C/D + A/B | 17501475 (100%) |
| Supplemented with propargyl-malonic acid | Stambomycin C/D + A/B | 80079185 (458%) |
| | Propargyl-stambomycin | n.d. |
| | Deoxy-propargyl-stambomycin | n.d. |

**l**

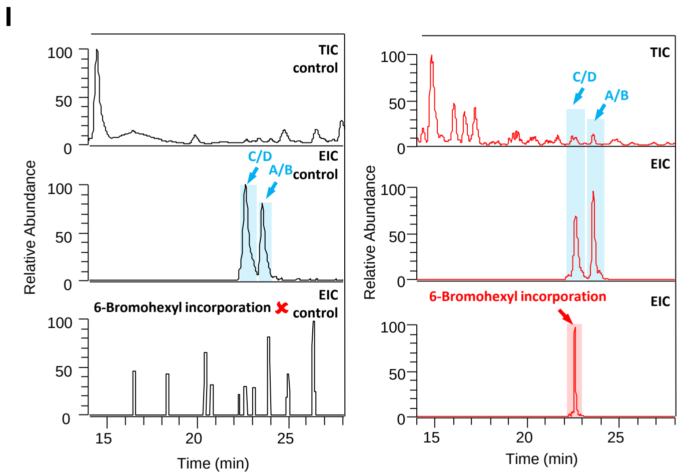


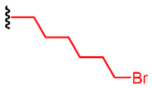


6-Bromohexyl-

| Mutant ATCC/OE484/Δ483 | Compounds | Integrated area of EIC peaks |
|---|---|---|
| Control (unsupplemented) | Stambomycin C/D + A/B | 17501475 (100%) |
| Supplemented with 6-bromohexyl-malonic acid | Stambomycin C/D + A/B | 106923327 (611%) |
| | 6-Bromohexyl-stambomycin (lacking 2H) **(11)** | 4817471 (27.5%) |
| | Deoxy-6-bromohexyl-stambomycin (lacking 2H) | n.d. |

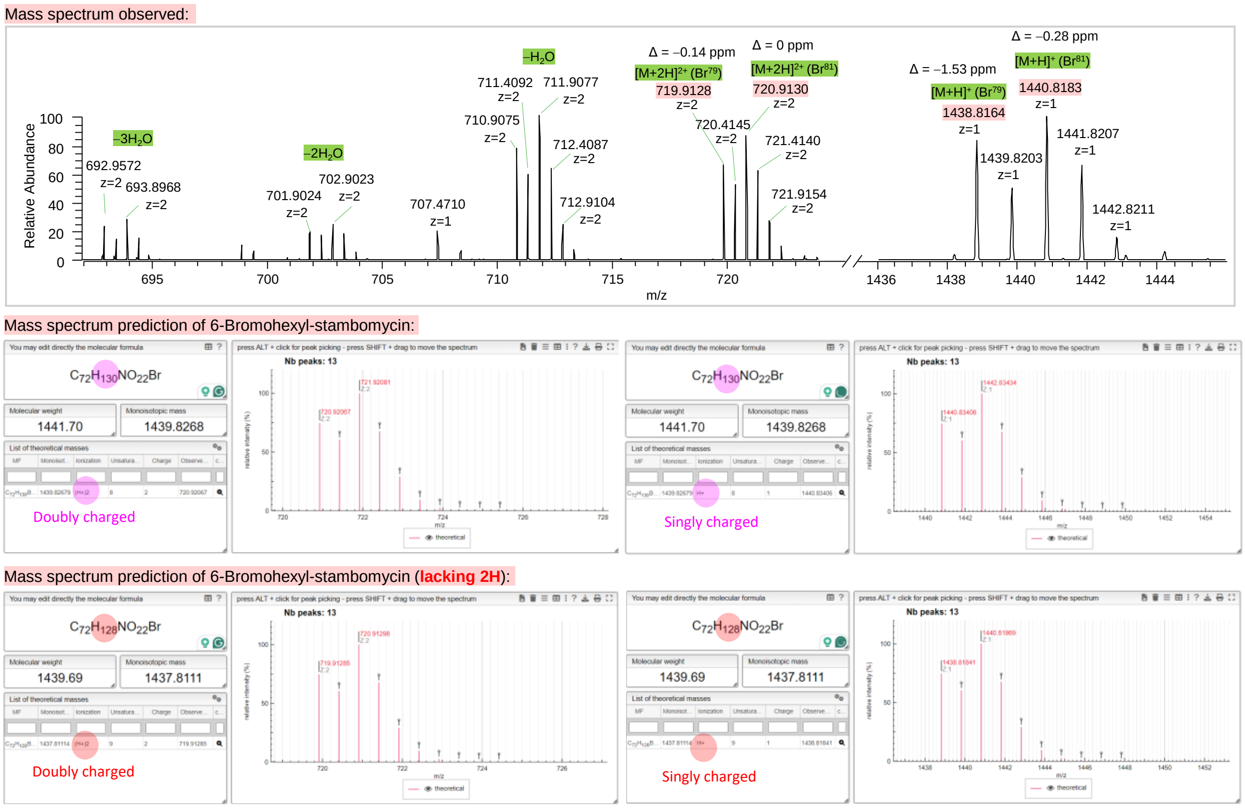


**Figure S3.** Analysis by mass spectrometry and UV detection of crude extract from the mutasynthesis strain ATCC/OE484/Δ483 supplemented with various malonic acid alternatives in comparison to the control ATCC/OE484/Δ483 without supplementation. In each case are shown: the TIC, EIC and $UV_{238}$ of crude fed-extracts, the mass spectrum of the corresponding stambomycin analogues, and the production levels of both the analogues and the native stambomycins A–D based on the integrated area of EIC peaks (average of at least two independent fermentations). The relative yield of stambomycins/analogues in the fed strain is also shown, which was calculated based on the production of stambomycins (C/D + A/B) by the mutasynthesis strain ATCC/OE484/Δ483 in the absence of supplementation (set to 100%). " n.d." indicates compounds which were not detected. (Note: the measured masses for the product resulting from incorporation of 6-bromohexyl-malonic acid exhibit a strong M+2 peak, and the characteristic bromine isotopic pattern. However, the observed mass is 2 Da lower than expected, which we propose arises from over-oxidation at the C-28 position.)

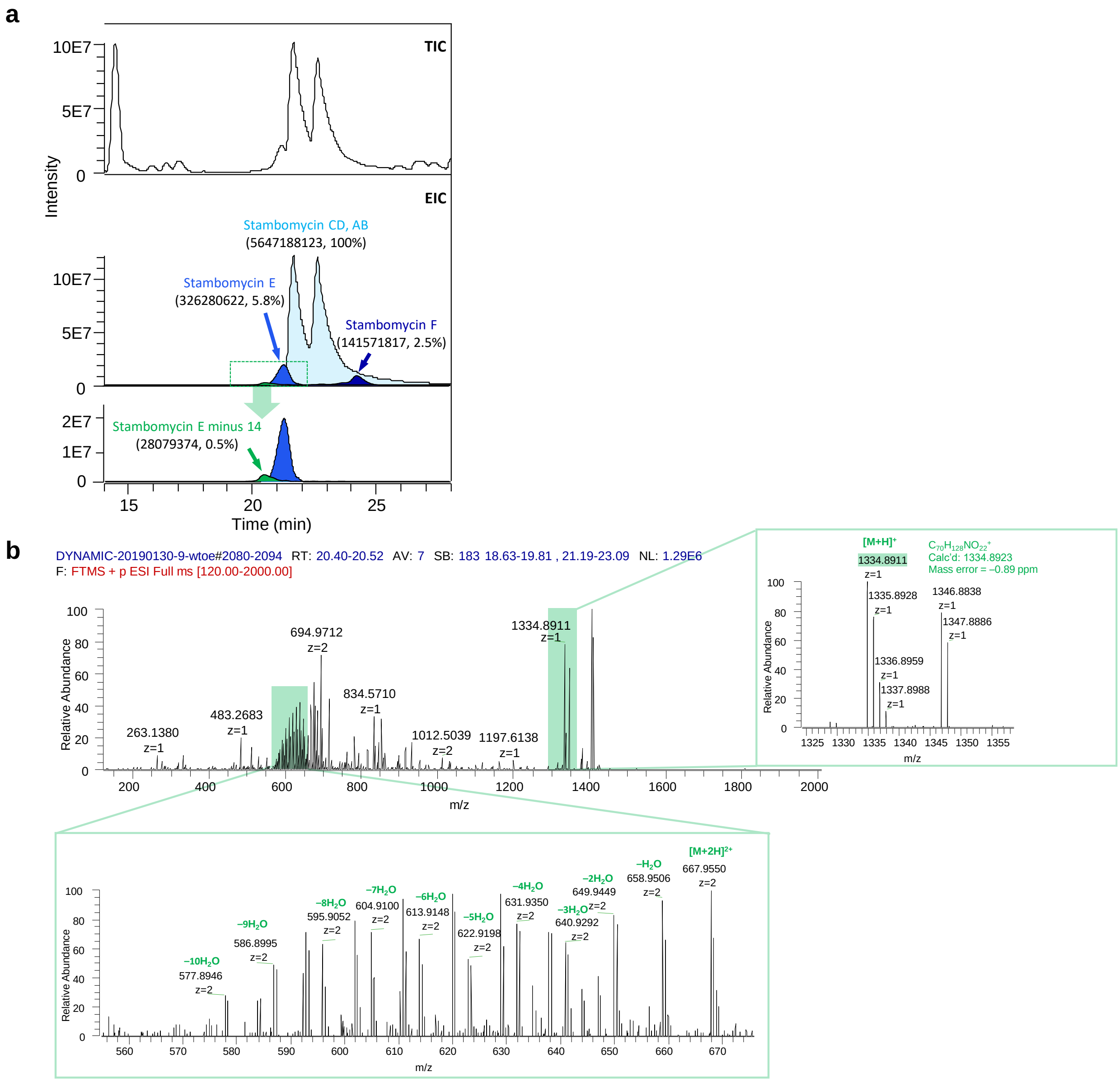


**Figure S4.** Analysis by mass spectrometry of the crude extract from the parental ATCC/OE484. **a**) TIC of the crude extract of ATCC/OE484 and the EIC of stambomycins A–F (representative of two replicates). The production of stambomycins E and F are only 5.8% and 2.5% of stambomycins A–D, as judged on the basis of relative peak areas. We also detected a mass consistent with stambomycin E minus 14, which we attribute to selection by module 13 of malonyl-CoA extender unit instead of the typical methylmalonyl-CoA. Nevertheless, malonyl-CoA remains favoured by $AT_{13}$ as the yield of stambomycin E (5.8%) is 12-times higher than the putative demethyl-stambomycin E (0.5%) in the parental strain. **b**) Mass spectra of dimethyl-stambomycin E, showing the characteristic multiple water losses[5].

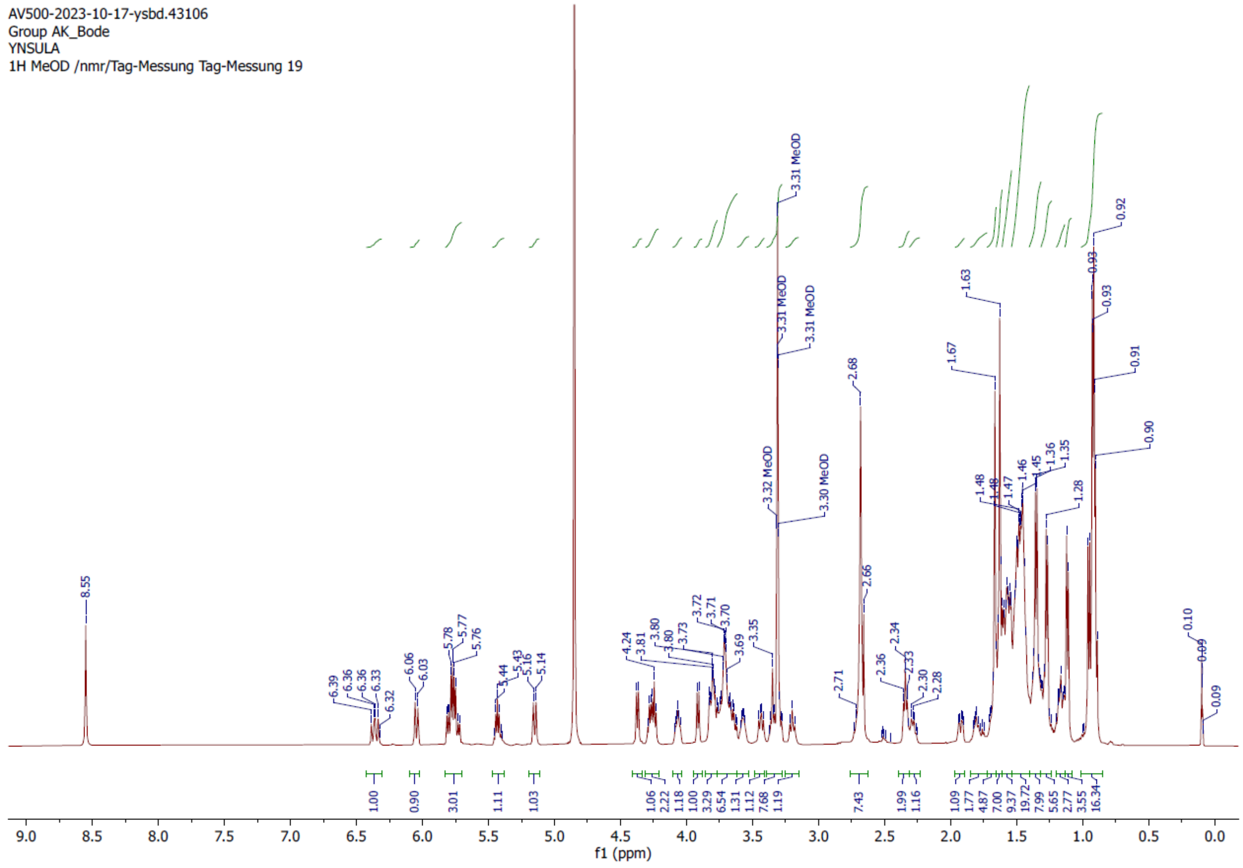


**Figure S5.** $^{1}$H NMR spectrum of deoxy-butyl-stambomycin (**5**).

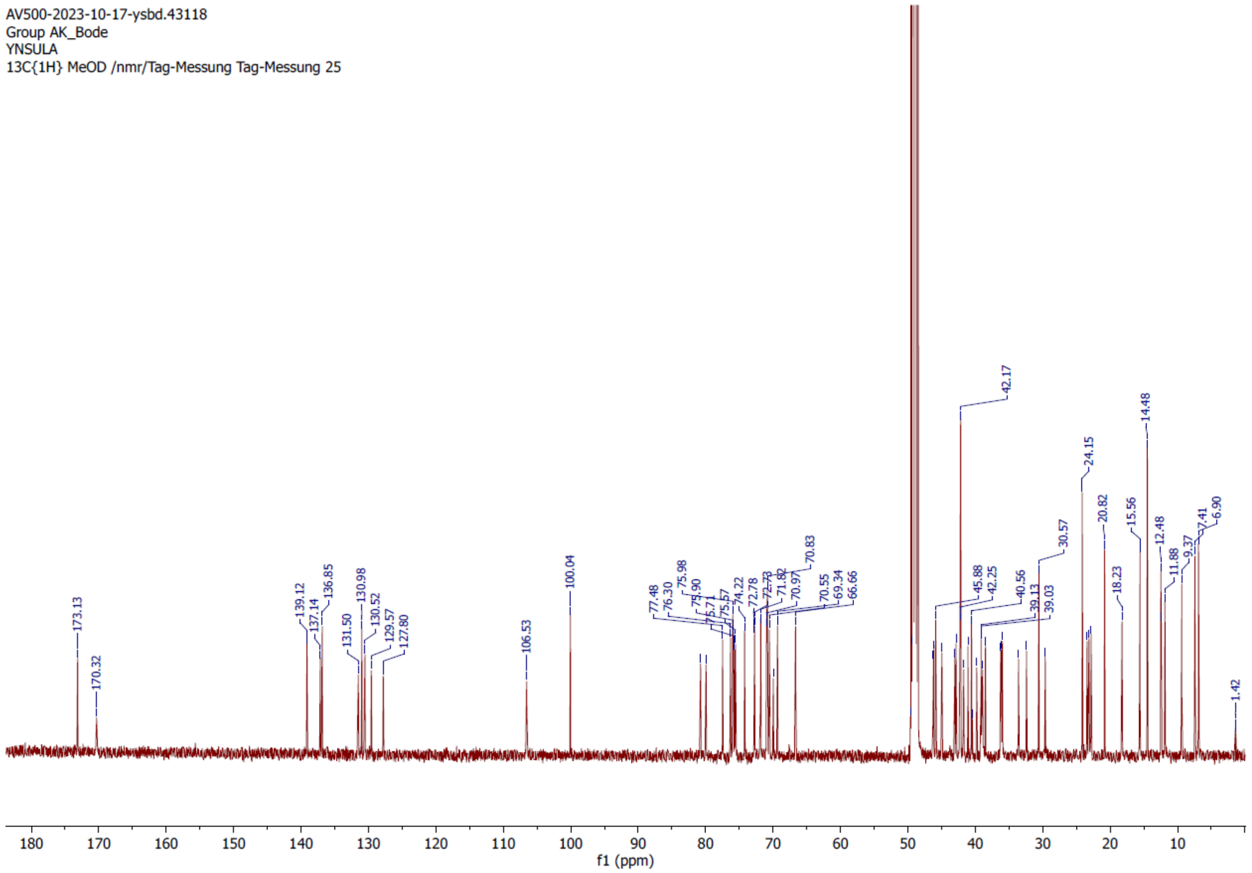


**Figure S6.** $^{13}$C NMR spectrum of deoxy-butyl-stambomycin (**5**).

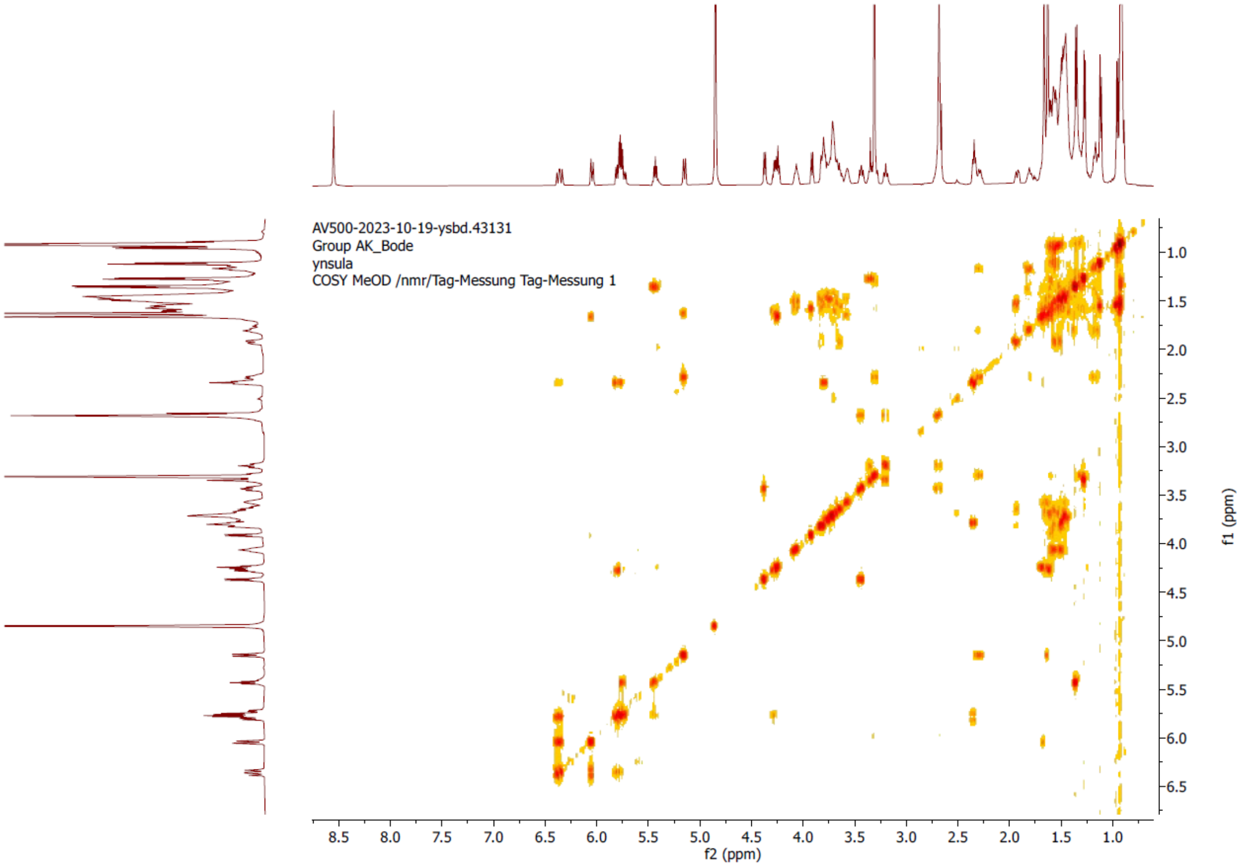


**Figure S7.** $^{1}H$-$^{1}H$ COSY spectrum of deoxy-butyl-stambomycin (**5**).

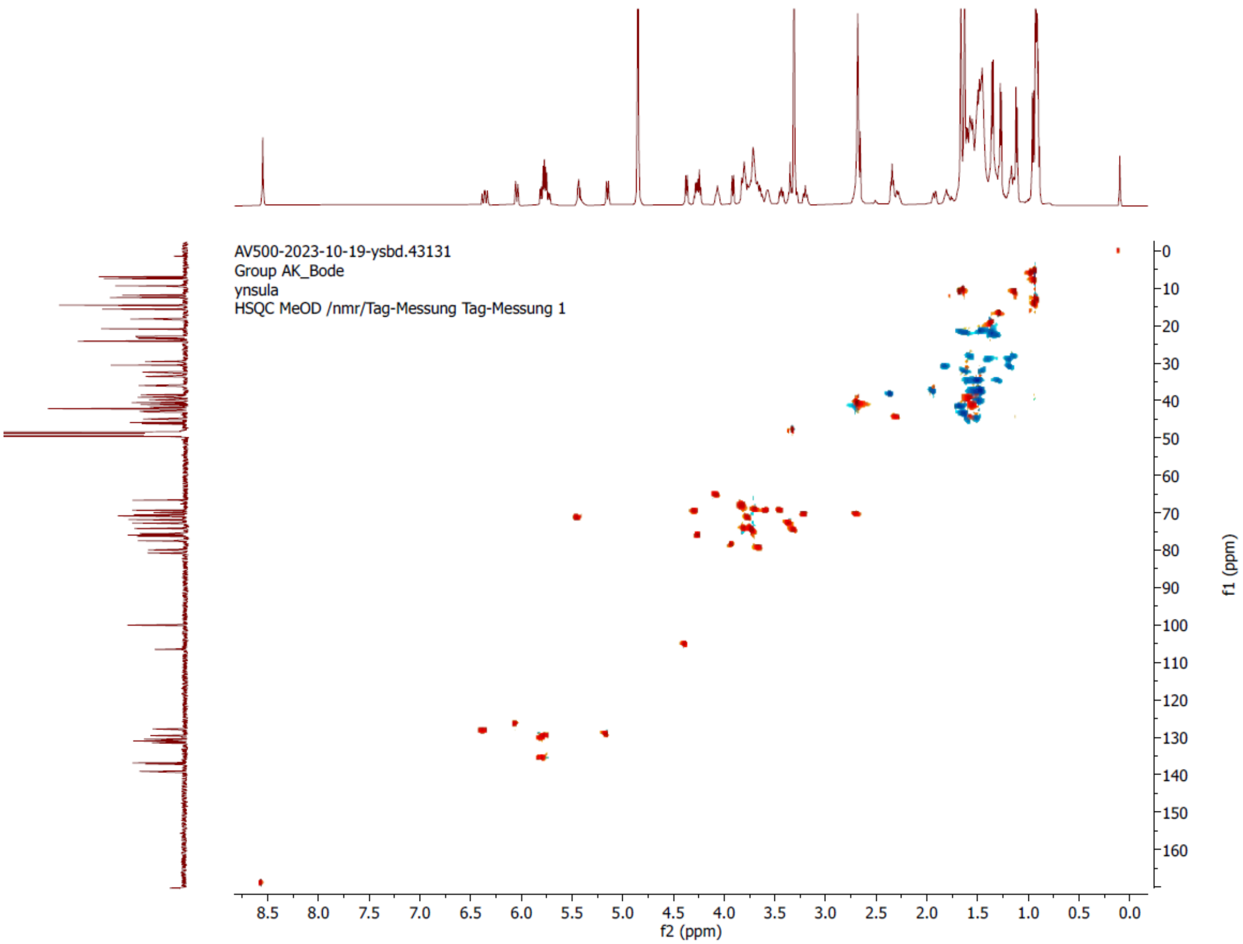


**Figure S8.** [$^{1}$H, $^{13}$C]-HSQC spectrum of deoxy-butyl-stambomycin (**5**).

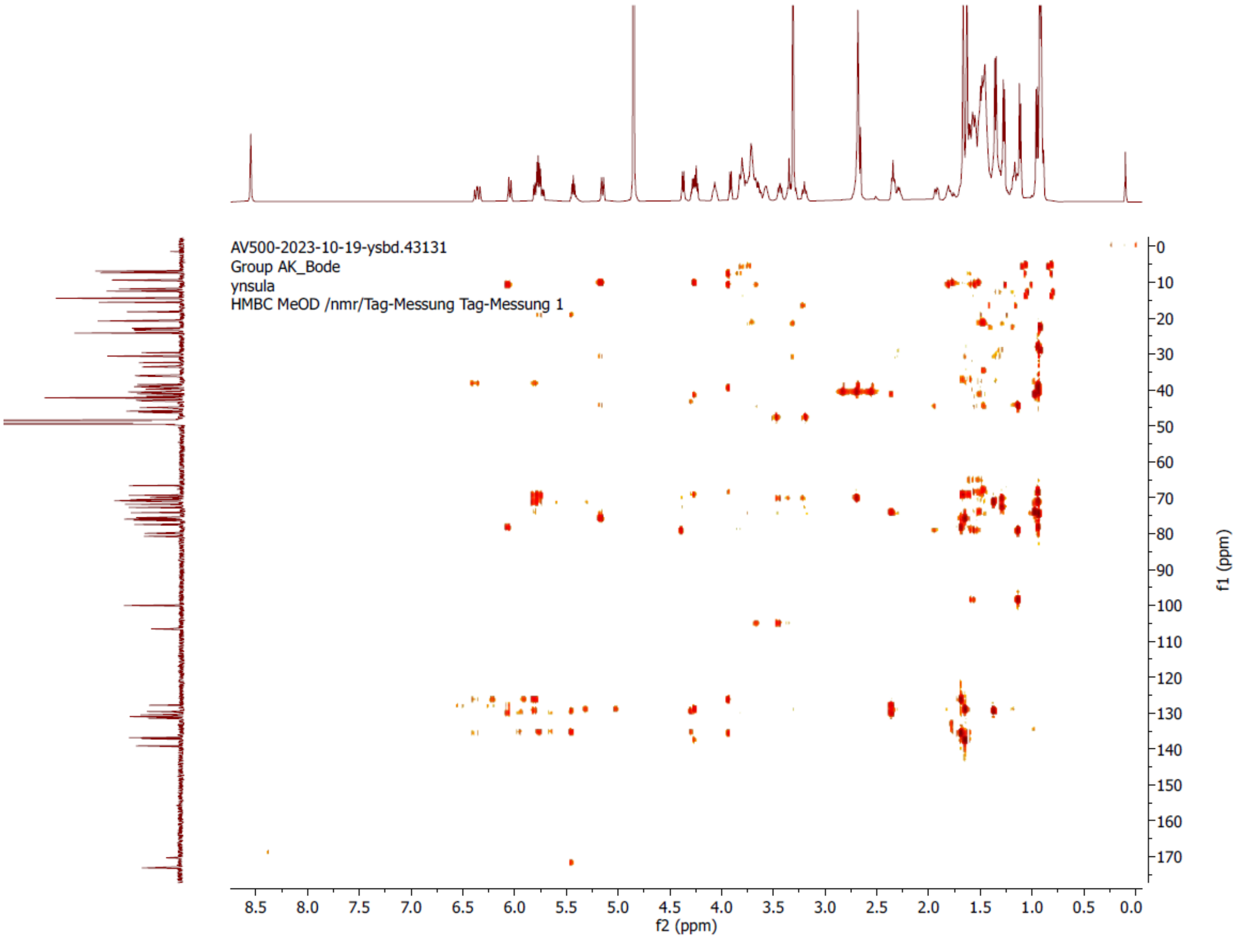


**Figure S9.** [$^{1}$H, $^{13}$C]-HMBC spectrum of deoxy-butyl-stambomycin (**5**).

Deoxy-butyl-stambomycin

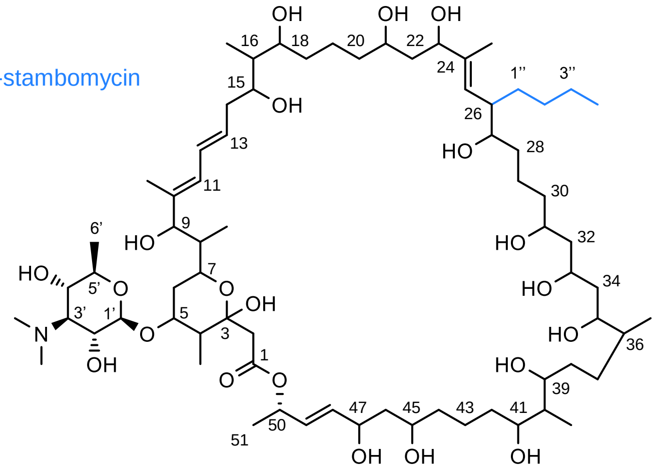

| Position | δC (ppm) | δH (ppm) | HMBC | Position | δC (ppm) | δH (ppm) | HMBC |
|---|---|---|---|---|---|---|---|
| 1 | 173.1 | | | 24-Me | 11.8 | 1.63 | |
| 2 | 43.0 | 2.70 | 10.0 | 25 | 129.5 | 6.36 | 15.0, 10.9 |
| 3 | 100 | | | 26 | 42.2 | 2.64 | |
| 4 | 45.8 | 2.29 | | 1″ | 32.4 | 1.15, 1.81 | |
| 4-Me | 12.4 | 1.11 | | 2″ | 24.1 | 1.30, 1.35 | |
| 5 | 80.7 | 3.65 | | 3″ | 36.2 | 1.58 | |
| 6 | 39.1 | 1.46, 1.90 | | 4″ | 15.5 | 0.91 | |
| 1′ | 106 | 4.37 | 7.4 | 27 | 76.3 | 3.70 | |
| 2′ | 70.5 | 3.68 | 10.6, 9.6 | 28 | 23.1 | 1.43 | |
| 3′ | 71.8 | 3.19 | | 29 | 29.6 | 1.13, 1.56 | |
| 3′-NMe2 | 42.1 | 2.66 | | 30 | 35.9 | 1.46 | |
| 4′ | 70.8 | 3.43 | 10.3, 7.4 | 31 | 69.3 | 3.81 | |
| 5′ | 74.2 | 3.35 | 9.4 | 32 | 46.2 | 1.47 | |
| 6′ | 18.2 | 1.28 | 5.9 | 33 | 66.6 | 4.07 | |
| 7 | 69.9 | 3.81 | | 34 | 39.7 | 2.35 | |
| 8 | 40.5 | 1.57 | | 35 | 72.7 | 3.75 | |
| 8-Me | 9.3 | 0.93 | | 36 | 41.0 | 1.57 | |
| 9 | 79.9 | 3.91 | 7.3 | 36-Me | 14.4 | 0.91 | |
| 10 | 137.1 | | | 37 | 30.5 | 1.18, 1.38 | |
| 10-Me | 12.4 | 1.66 | | 38 | 38.9 | 1.47, 1.94 | |
| 11 | 127.8 | 6.04 | 10.8 | 39 | 75.9 | 3.30 | 7.3 |
| 12 | 130.5 | 5.15 | 15.0, 10.2 | 40 | 41.7 | 1.48 | |
| 13 | 131.5 | 5.78 | 15.4, 8.3 | 40-Me | 6.9 | 0.91 | |
| 14 | 40.4 | 2.66 | | 41 | 75.9 | 3.70 | |
| 15 | 75.7 | 3.72 | | 42 | 33.5 | 143, 1.60 | |
| 16 | 42.8 | 1.62 | | 43 | 22.8 | 1.43 | |
| 16-Me | 7.4 | 0.96 | | 44 | 39.0 | 1.54, 1.91 | |
| 17 | 75.5 | 3.79 | | 45 | 71.8 | 2.68 | |
| 18 | 36.1 | 1.56, 1.46 | | 46 | 46.1 | 1.57 | |
| 19 | 23.4 | 1.59, 1.34 | | 47 | 70.9 | 4.27 | 12.7, 6.7 |
| 20 | 38.4 | 1.45 | | 48 | 136.8 | 5.77 | 15.4, 8.3 |
| 21 | 70.8 | 3.57 | | 49 | 130.9 | 5.73 | |
| 22 | 44.9 | 1.6 | | 50 | 72.7 | 5.44 | 6.4 |
| 23 | 77.4 | 4.23 | 6.7 | 51 | 20.8 | 1.35 | 6.4 |
| 24 | 139.1 | | | | | | |

**Figure S10.** Structure and summary of the NMR data for deoxy-butyl-stambomycin (**5**). The $^{13}$C NMR spectrum exhibits signals accounting for 70 carbons. Comparison of the $^{1}$H and $^{13}$C NMR data acquired on deoxy-butyl-stambomycin with that reported for stambomycin C[4], revealed that the two compounds share the same macrocyclic structure, except for the alkyl chain present at position C-26. The HMBC correlations of H-50 ($\delta_C$ 5.44) with the C-1 ($\delta_C$ 173.1), C-51 ($\delta_C$ 72.7), C-49 ($\delta_C$ 130.9) and C-48 ($\delta_C$ 136.8), show that a closed macrocycle is present. Connections between carbons and their respective protons were assigned based on the HSQC spectrum. The coupling constant are indicated in the table above when possible. Together, these data confirmed the identity of the metabolite as deoxy-butyl-stambomycin.

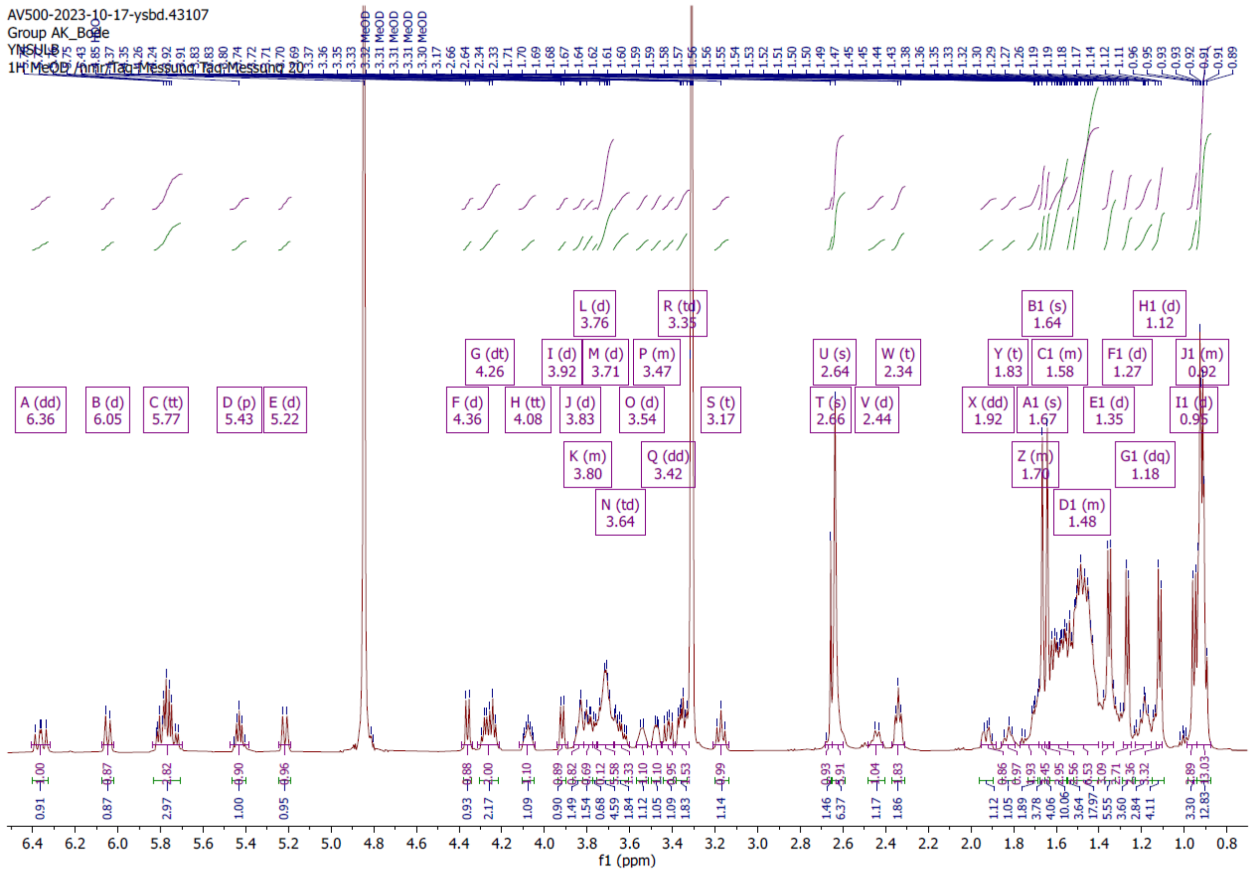


**Figure S11.** $^{1}$H NMR spectrum of butyl-stambomycin (**4**).

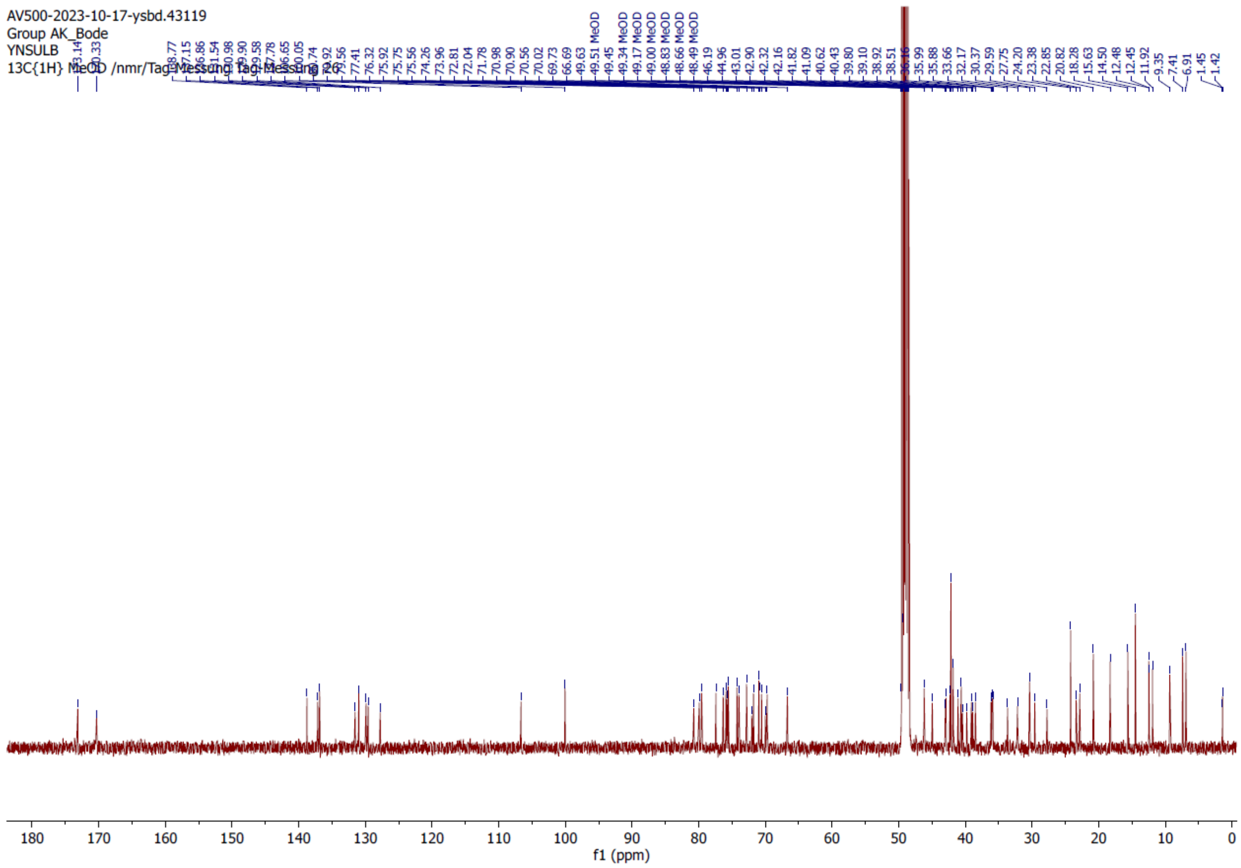


**Figure S12.** $^{13}C$ NMR spectrum of butyl-stambomycin (**4**).

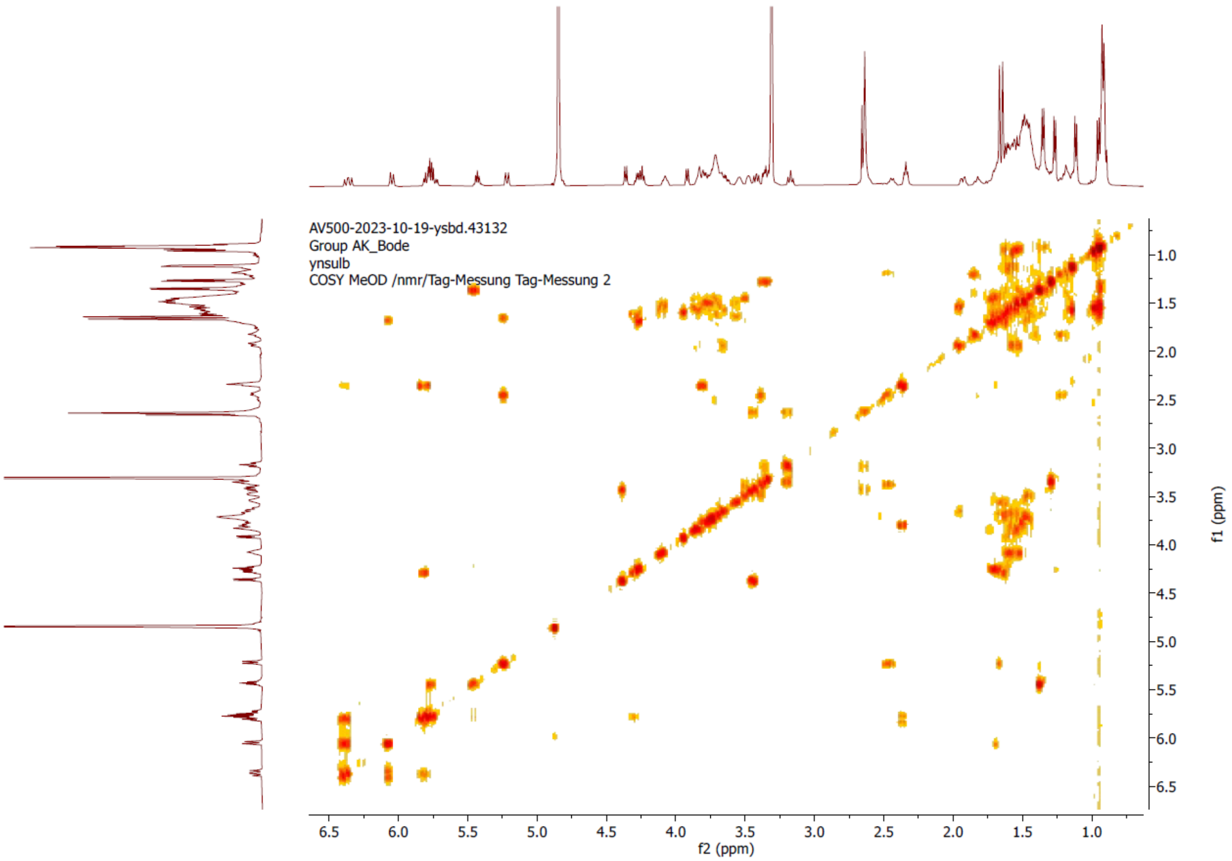


**Figure S13.** $^{1}$H-$^{1}$H COSY spectrum of butyl-stambomycin (**4**).

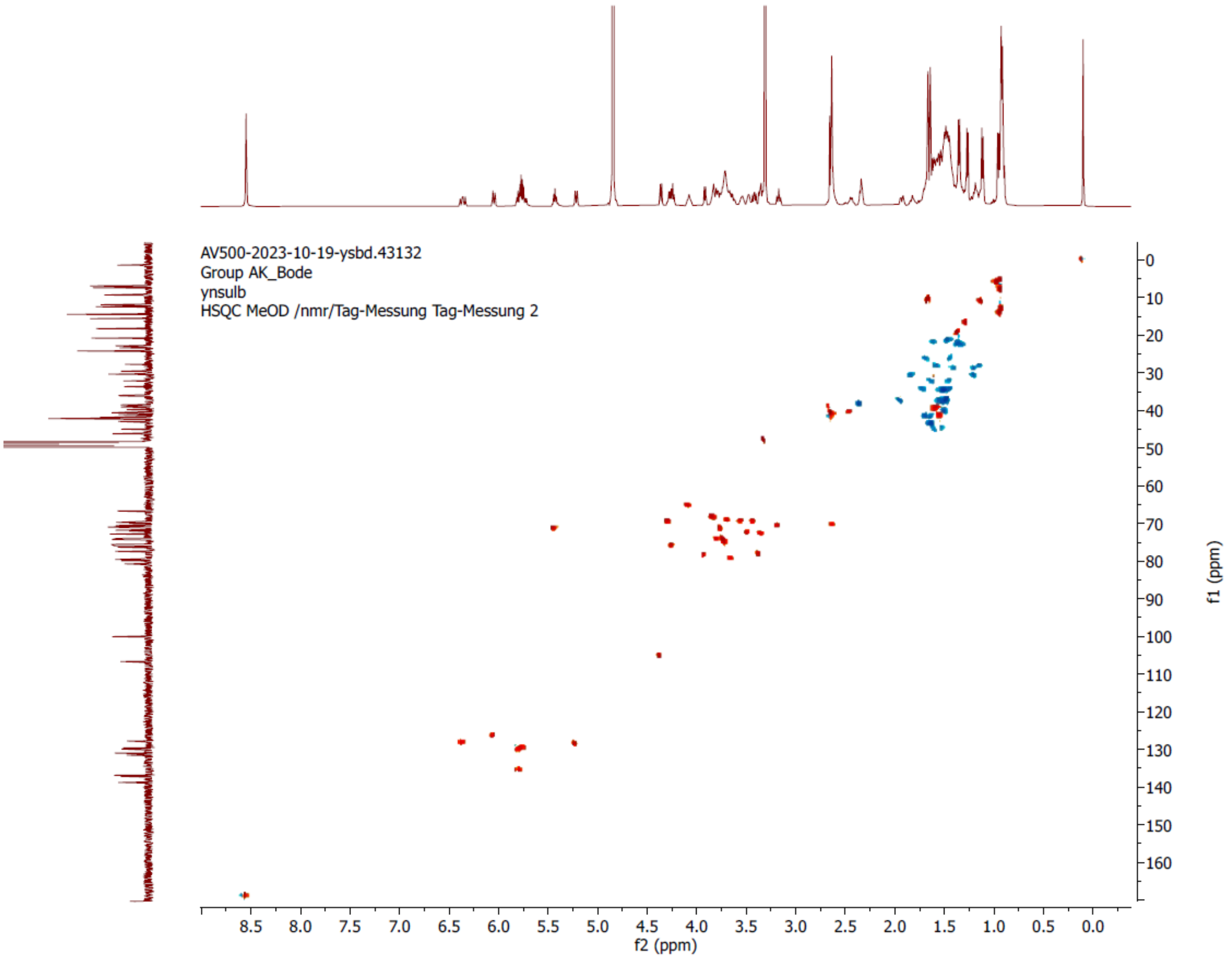


259

260 **Figure S14.** [$^1$H, $^{13}$C]-HSQC spectrum of butyl-stambomycin (**4**).

261

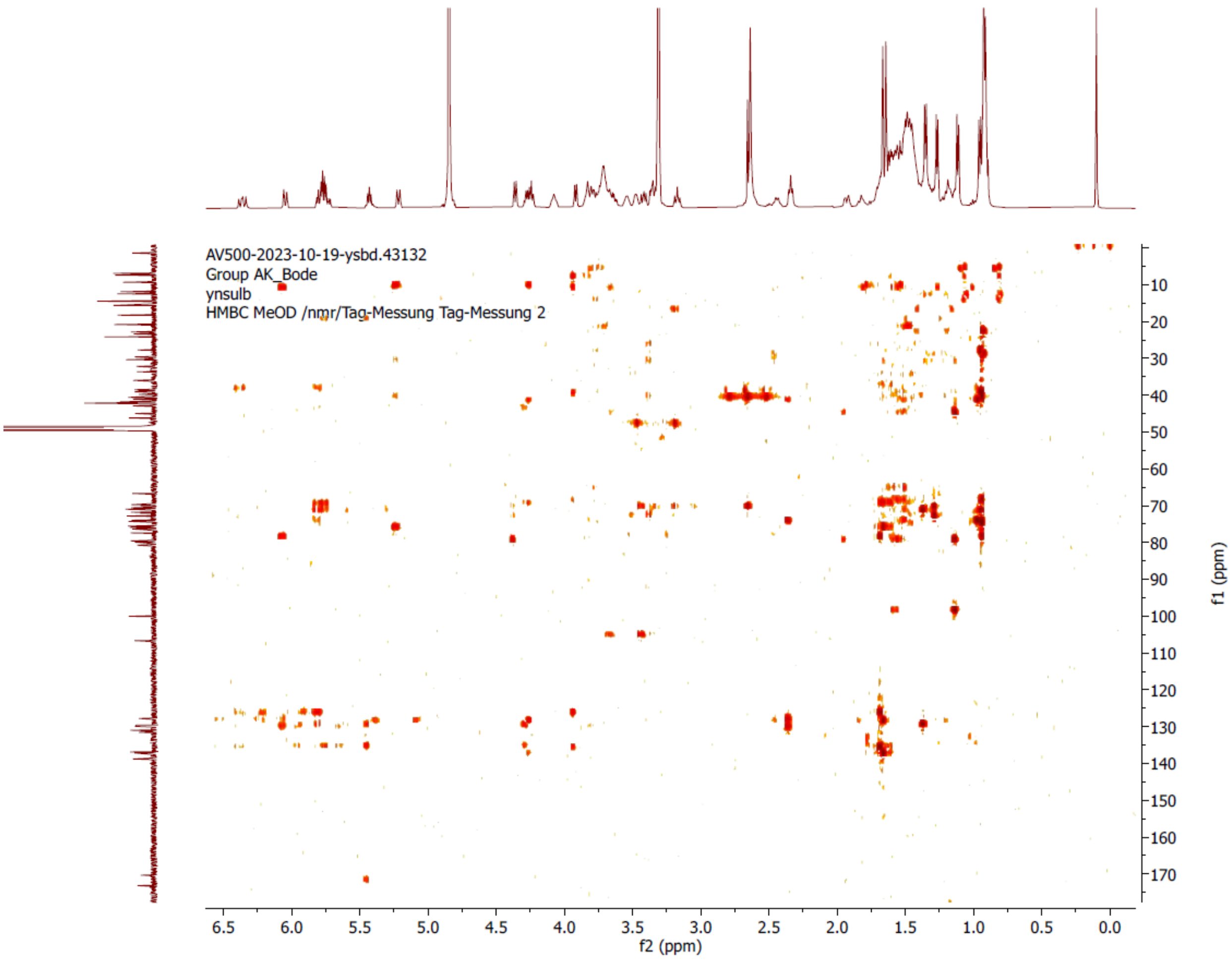


**Figure S15.** [$^{1}$H, $^{13}$C]-HSQC HMBC spectrum of butyl-stambomycin (**4**).

Butyl-stambomycin

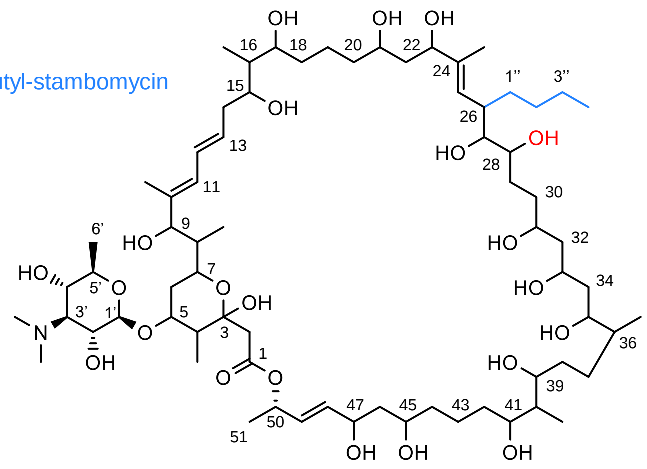


| Position | δC (ppm) | δH (ppm) | HMBC | Position | δC (ppm) | δH (ppm) | HMBC |
|---|---|---|---|---|---|---|---|
| 1 | 173.1 | | | 24-Me | 11.9 | 1.64 | |
| 2 | 43.0 | 1.62, 1.67 | | 25 | 129.9 | 5.22 | 10.4 |
| 3 | 100.0 | | | 26 | 40.4 | 2.65 | |
| 4 | 46.1 | 1.51, 1.60 | | 1″ | 30.3 | 1.18, 1.38 | |
| 4-Me | 12.4 | 1.12 | | 2″ | 24.2 | 1.34 | |
| 5 | 80.7 | 3.64 | | 3″ | 36.1 | 1.47, 1.71 | |
| 6 | 39.1 | 1.47 | | 4″ | 14.5 | 0.91 | |
| 1′ | 106.6 | 4.36 | 7.4 | 27 | 79.5 | 3.35 | |
| 2′ | 70.5 | 3.68 | 7.4, 10.7 | 28 | 75.9 | 3.71 | |
| 3′ | 72.0 | 3.16 | 9.4, 9.4 | 29 | 27.7 | 1.43, 1.68 | |
| 3′-$NMe_2$ | 42.1 | 2.44 | | 30 | 35.9 | 1.47, 1.71 | |
| 4′ | 70.0 | 3.82 | | 31 | 70.9 | 3.55 | |
| 5′ | 73.9 | 3.47 | 10.3, 7.4 | 32 | 46.1 | 1.51, 1.60 | |
| 6′ | 18.2 | 1.27 | | 33 | 66.6 | 4.08 | |
| 7 | 69.7 | 3.81 | | 34 | 41.8 | 1.48, 2.44 | |
| 8 | 41.0 | 1.57 | | 35 | 72.8 | 3.74 | |
| 8-Me | 9.3 | 0.91 | | 36 | 40.6 | 1.57 | |
| 9 | 79.9 | 3.92 | 7.2 | 36-Me | 15.6 | 0.94 | |
| 10 | 137.1 | | | 37 | 29.5 | 1.13, 1.57 | |
| 10-Me | 12.4 | 1.66 | | 38 | 33.6 | 1.44, 1.62 | |
| 11 | 128.7 | 6.05 | 10.9 | 39 | 76.3 | 3,75 | |
| 12 | 129.5 | 6.36 | 15.0, 10.9 | 40 | 42.1 | 1.48 | |
| 13 | 131.5 | 5.77 | 15.5, 7.4 | 40-Me | 6.9 | 0.91 | |
| 14 | 39.8 | 2.34 | 6.9 | 41 | 75.7 | 3.71 | |
| 15 | 75.5 | 3.77 | | 42 | 32.1 | 1.19, 1.82 | |
| 16 | 42.9 | 1.57 | | 43 | 23.3 | 1.59 | |
| 16-Me | 7.4 | 0.95 | | 44 | 38.9 | 1.46, 1.94 | |
| 17 | 74.2 | 3.77 | | 45 | 70.9 | 3.55 | |
| 18 | 35.8 | 1.47, 1.71 | | 46 | 44.9 | 1.63 | |
| 19 | 22.8 | 1.45 | | 47 | 71.7 | 2.61 | |
| 20 | 38.5 | 1.46 | | 48 | 136.8 | 5.77 | 15.5, 7.4 |
| 21 | 70.9 | 3.43 | | 49 | 130.9 | 5.77 | 15.5, 7.4 |
| 22 | 42.3 | 2.61 | | 50 | 72.8 | 5.42 | 6.6, 6.4 |
| 23 | 77.4 | 4.26 | 6.6 | 51 | 20.8 | 1.35 | 6.6 |
| 24 | 138.7 | | | | | | |

**Figure S16.** Structure and summary for the NMR data of butyl-stambomycin (**4**). The $^{1}H$ NMR data obtained on butyl-stambomycin are highly similar to those for butyl-deoxystambomycin, with the exception of the deshielded proton attached to C-28. Furthermore, the $^{13}C$ NMR spectrum of butyl-stambomycin also include 70 carbon signals, except that the C-28 carbon is relatively deshielded ($\delta_C$ 76.3, $\delta_H$ 3.75), consistent with the presence of the C-28 hydroxyl. The data corresponding to the alkyl chain at C-26 are similar to those of butyl-deoxystambomycin, allowing assignment of the side chain as *n*-butyl. Together, these data confirmed the identity of the compound as butyl-stambomycin.

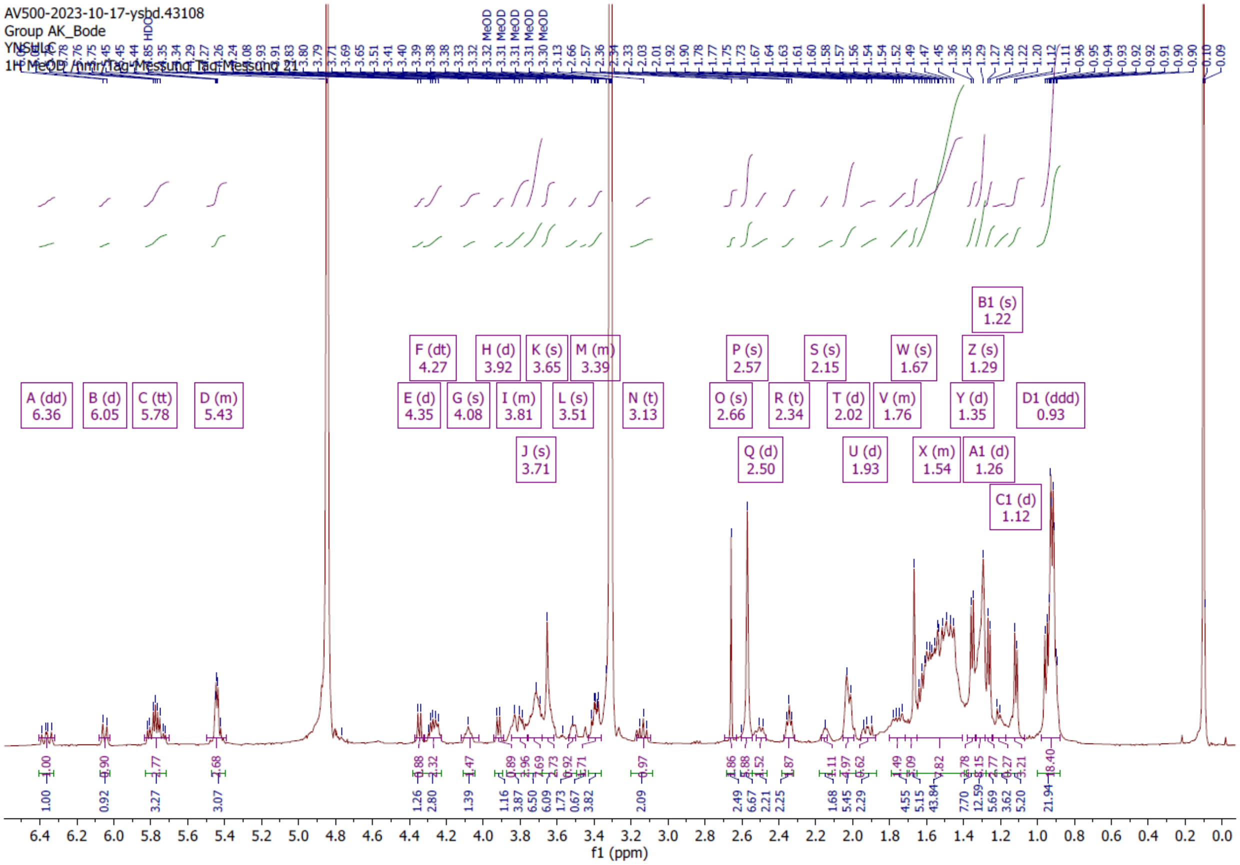


**Figure S17.** [1]H NMR spectrum of C-24-demethyl-butyl-stambomycin (**12**).

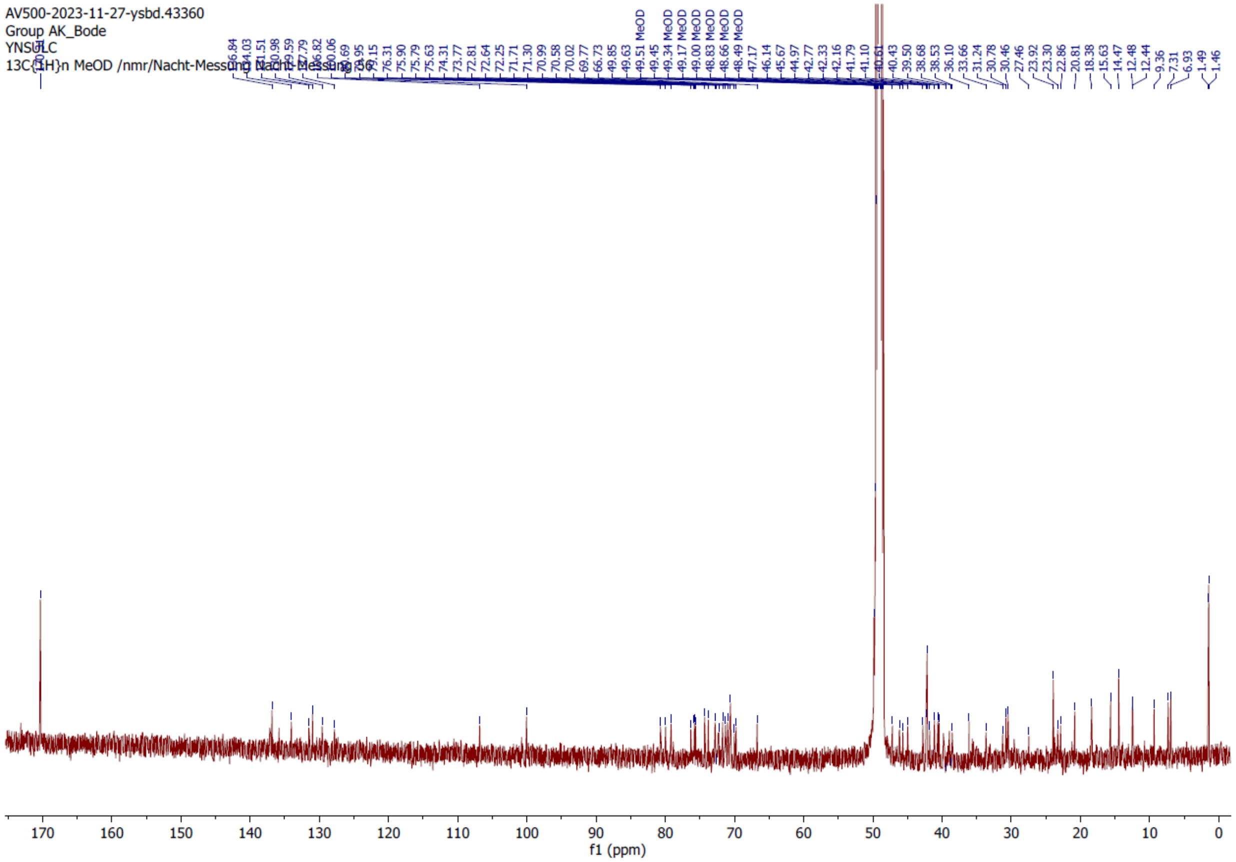


**Figure S18.** $^{13}$C NMR spectrum of C-24-demethyl-butyl-stambomycin (**12**).

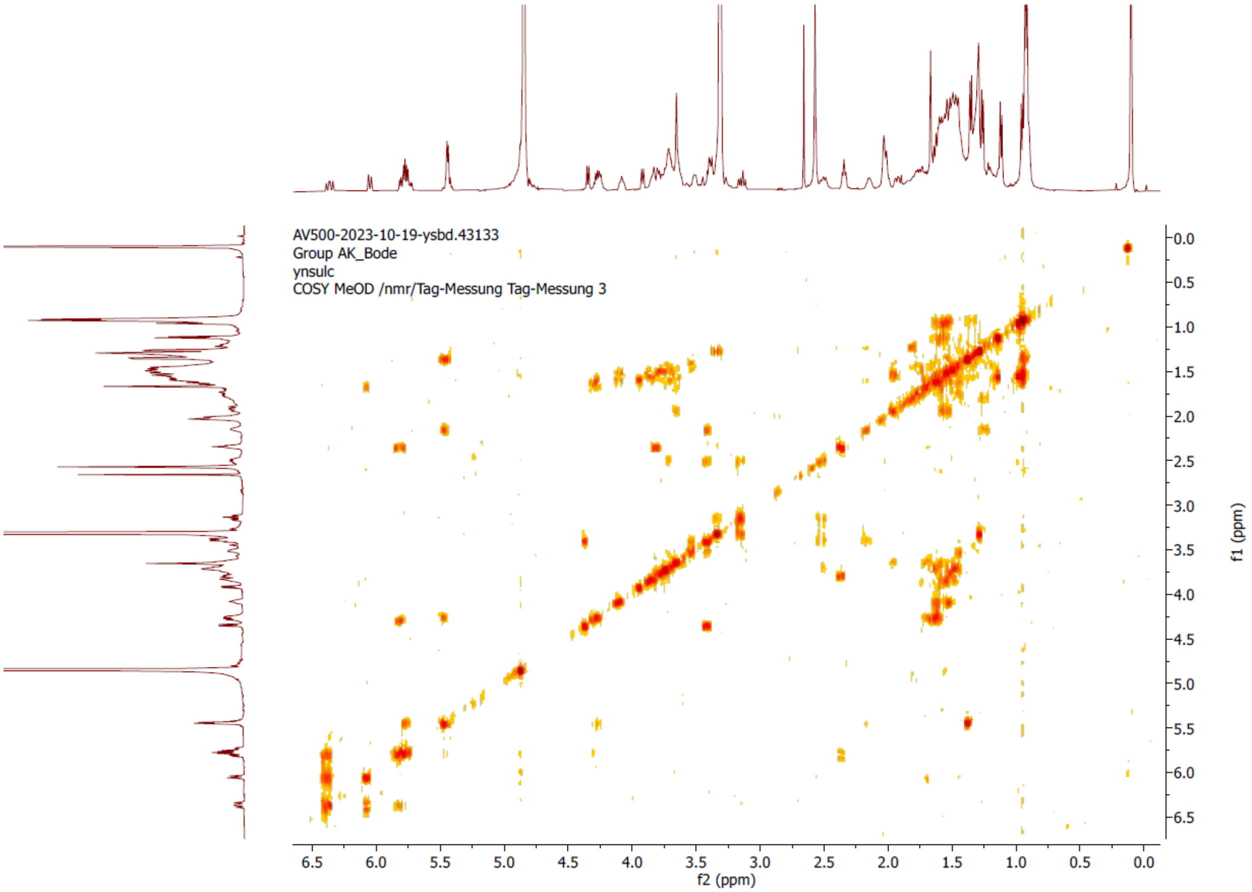


**Figure S19.** $^{1}H$-$^{1}H$ COSY spectrum of C-24-demethyl-butyl-stambomycin (**12**).

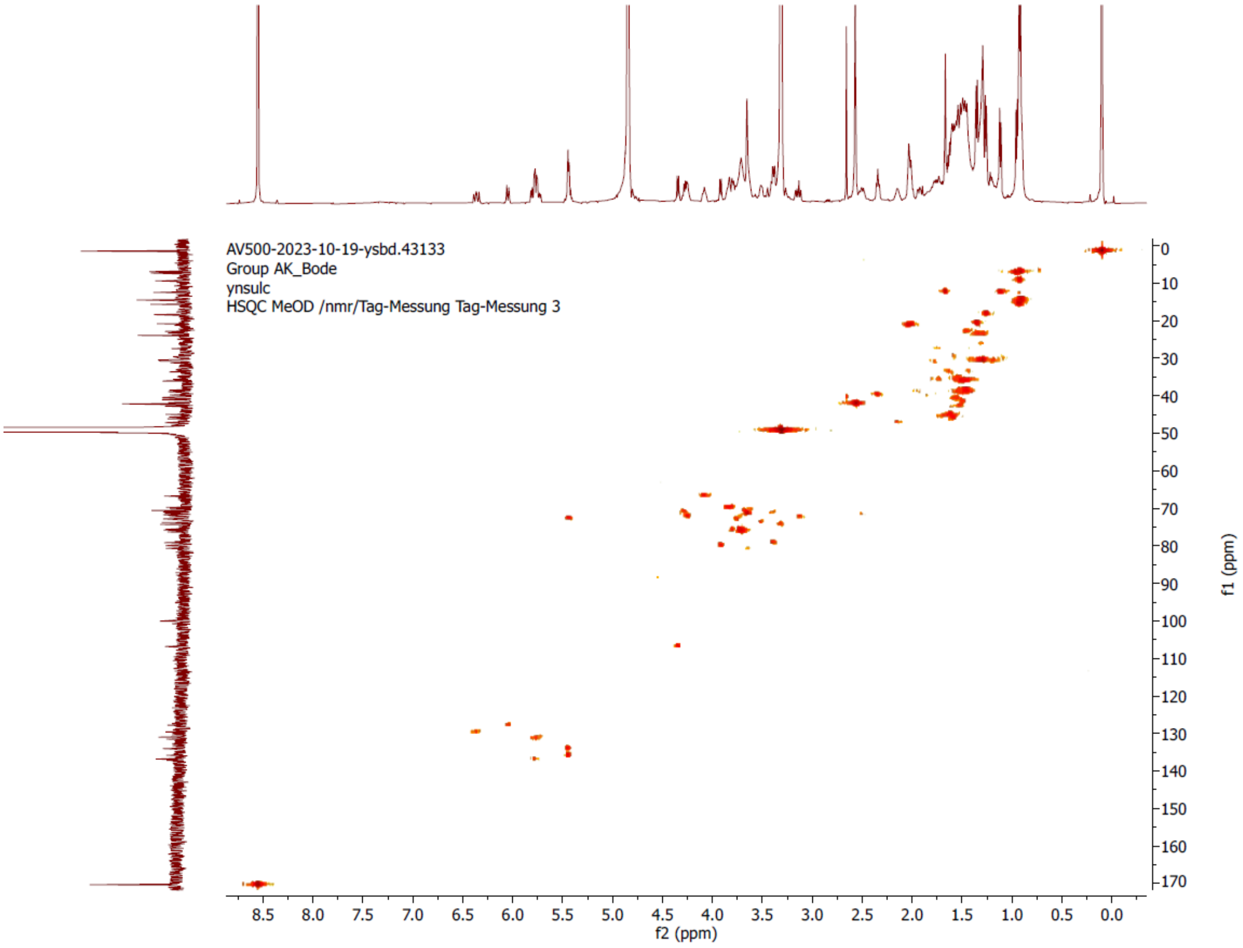


**Figure S20.** [$^{1}$H,$^{13}$C]-HSQC spectrum of C-24-demethyl-butyl-stambomycin (**12**).

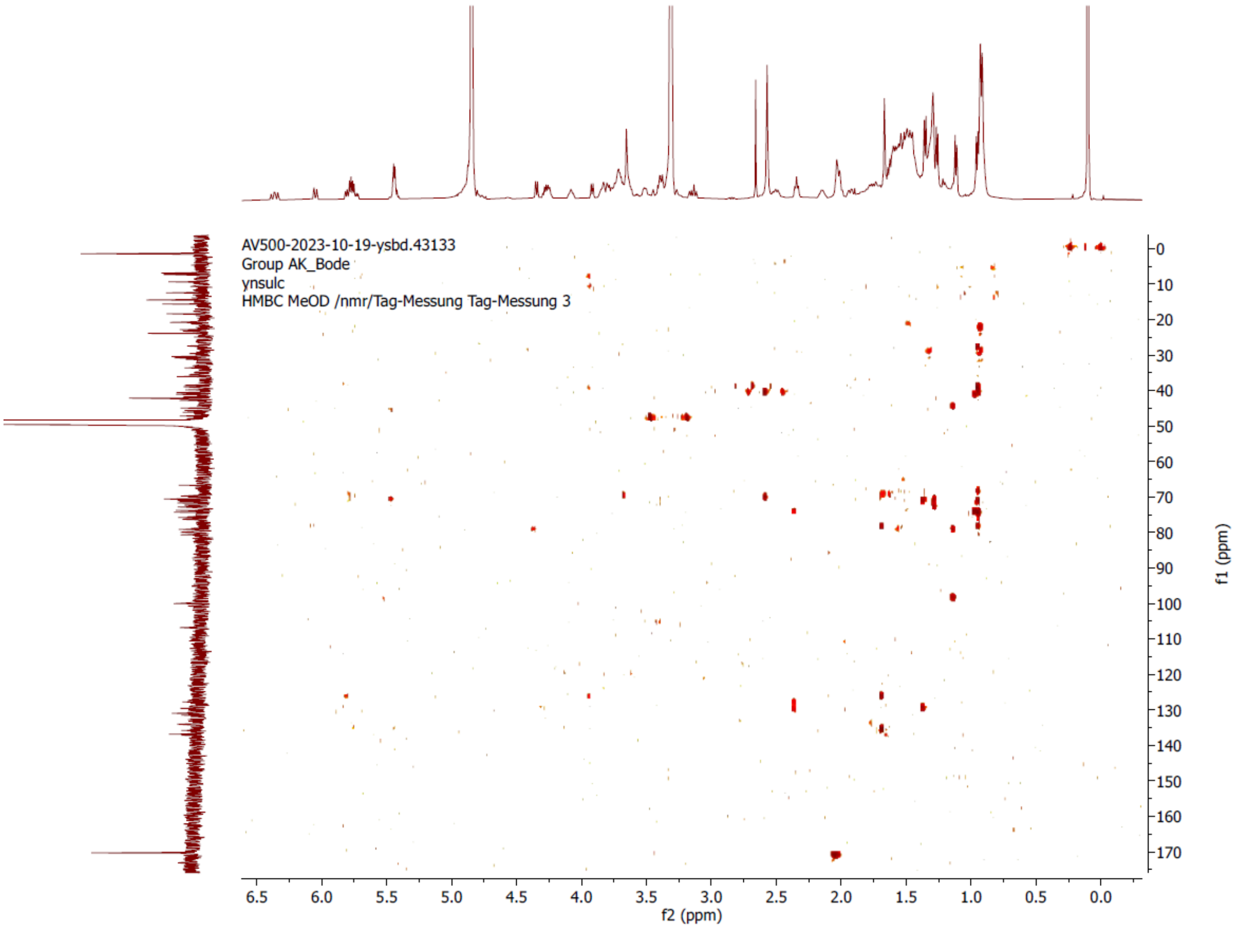


**Figure S21.** [$^{1}$H,$^{13}$C]-HMBC spectrum of C-24-demethyl-butyl-stambomycin (**12**).

C-24-demethyl-butyl-stambomycin

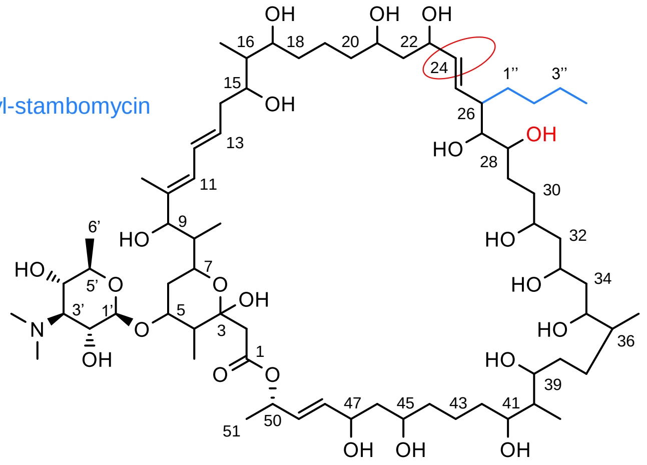


| Position | δC (ppm) | δH (ppm) | HMBC | Position | δC (ppm) | δH (ppm) | HMBC |
|---|---|---|---|---|---|---|---|
| 1 | 171.6 | | | 25 | 130.9 | 5.78 | 15.4, 8.0 |
| 2 | 44.9 | 1.60-1.68 | | 26 | 40.4 | 1.54 | |
| 3 | 100.0 | | | 1″ | 30.4 | 1.29 | |
| 4 | 46.1 | 1.54 | | 2″ | 23.9 | 1.34 | |
| 4-Me | 12.4 | 1.12 | | 3″ | 36.1 | 1.54 | |
| 5 | 80.6 | 3.65 | | 4″ | 14.4 | 0.93 | |
| 6 | 38.9 | 1.46 | | 27 | 79.1 | 3.39 | |
| 1′ | 106.8 | 4.35 | 7.5 | 28 | 75.9 | 3.39 | |
| 2′ | 70.5 | 3.66 | | 29 | 27.4 | 1.42, 1.76 | |
| 3′ | 71.7 | 3.64 | | 30 | 35.4 | 1.47, 1.71 | |
| 3′-$NMe_2$ | 42.1 | 2.50 | | 31 | 70.9 | 4.27 | |
| 4′ | 70.0 | 3.81 | | 32 | 45.6 | 1.54 | |
| 5′ | 72.8 | 3.71 | | 33 | 66.7 | 4.08 | |
| 6′ | 18.3 | 1.26 | | 34 | 41.7 | 1.54 | |
| 7 | 69.7 | 3.81 | | 35 | 72.8 | 3.81 | |
| 8 | 41.1 | 1.57 | | 36 | 40.6 | 1.54 | |
| 8-Me | 9.3 | 0.93 | | 36-Me | 15.6 | 0.93 | |
| 9 | 79.9 | 3.92 | 7.3 | 37 | 30.7 | 1.29 | |
| 10 | 136.8 | | | 38 | 33.6 | 1.54, 1.67 | |
| 10-Me | 12.4 | 1.67 | | 39 | 75.7 | 3.68 | |
| 11 | 127.7 | 6.05 | 10.8 | 40 | 42.1 | 1.54 | |
| 12 | 129.5 | 6.36 | 15.0, 11.0 | 40-Me | 6.9 | 0.93 | |
| 13 | 134.0 | 5.43 | | 41 | 73.7 | 3.51 | |
| 14 | 39.8 | 2.34 | 6.9 | 42 | 31.2 | 1.76 | |
| 15 | 75.6 | 3.70 | | 43 | 23.3 | 1.34 | |
| 16 | 42.7 | 1.53 | | 44 | 38.6 | 1.54 | |
| 16-Me | 7.3 | 0.93 | | 45 | 71.3 | 3.71 | |
| 17 | 74.3 | 3.39 | | 46 | 47.1 | 2.15 | |
| 18 | 35.6 | 1.54 | | 47 | 72.2 | 3.13 | |
| 19 | 22.8 | 1.54 | | 48 | 135.8 | 5.43 | |
| 20 | 38.5 | 1.54 | | 49 | 131.5 | 5.78 | 15.4, 8.0 |
| 21 | 70.9 | 3.65 | | 50 | 72.2 | 3.13 | 9.4, 9.4 |
| 22 | 42.3 | 1.54 | | 51 | 20.8 | 1.35 | 6.4 |
| 23 | 76.3 | 3.72 | 7.6 | | | | |
| 24 | 137.1 | 5.78 | 15.4, 8.0 | | | | |

**Figure S22.** Structure and summary for the NMR data of C-24-demethyl-butyl-stambomycin (**12**). The NMR data obtained on this compound are overall similar to those for butyl-stambomycin and deoxy-butyl-stambomycin, except that C-24-demethyl-butyl-stambomycin is missing a methyl signal at ($\delta_H$ 1.64) corresponding to the methyl at C-24, and the chemical shift of an olefinic proton ($\delta_H$ 5.22 in the case of butyl-stambomycin and deoxy-butyl-stambomycin) has shifted to ($\delta_H$ 5.43). Additionally, coupling constants of 15.4 and 8.0 were observed for H-25, consistent with the presence of protons on both H-24 and H-26. Together, these data confirmed the identity of the metabolite as C-24-demethyl-butyl-stambomycin.

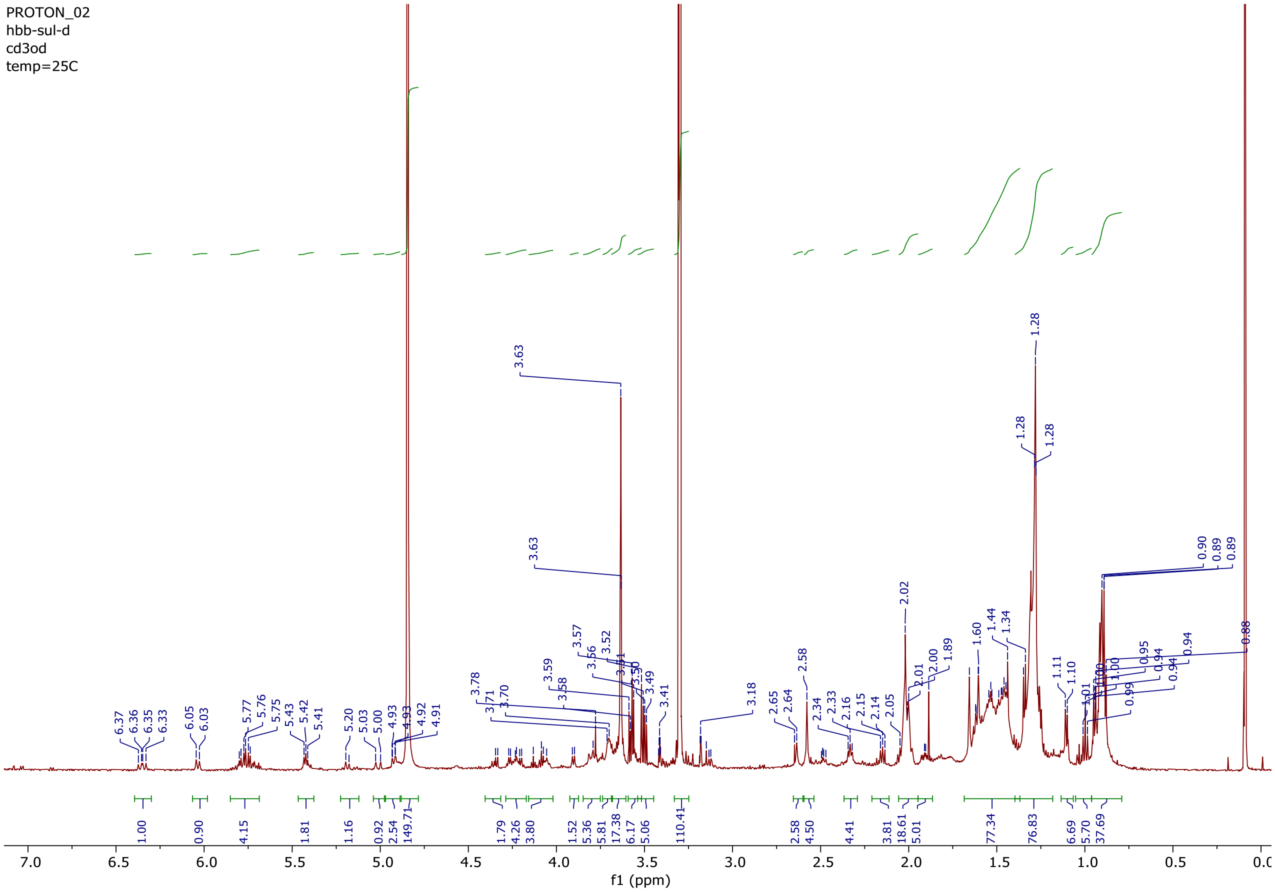


**Figure S23.** $^{1}$H NMR spectrum of deoxy-allyl-stambomycin (**3**).

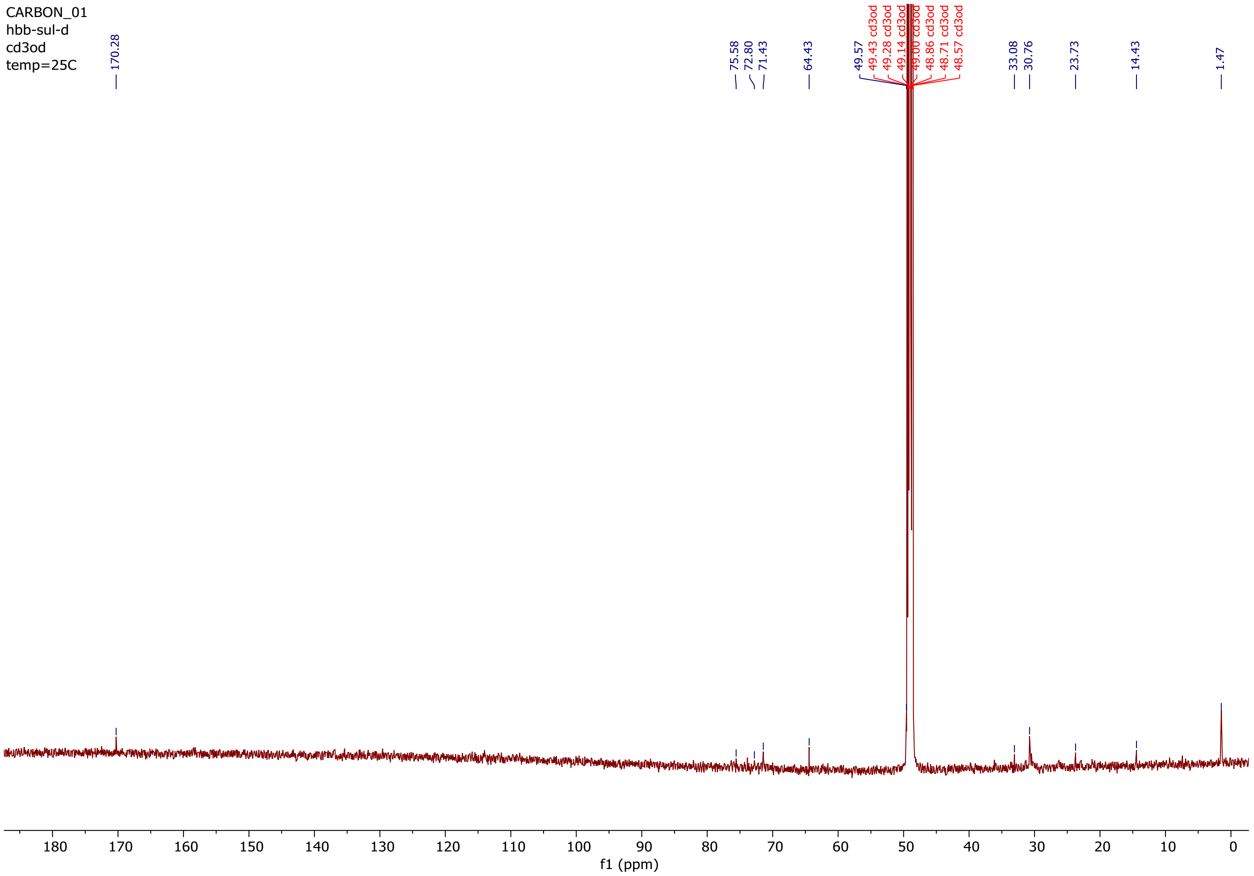


**Figure S24.** $^{13}C$ NMR spectrum of deoxy-allyl-stambomycin (**3**).

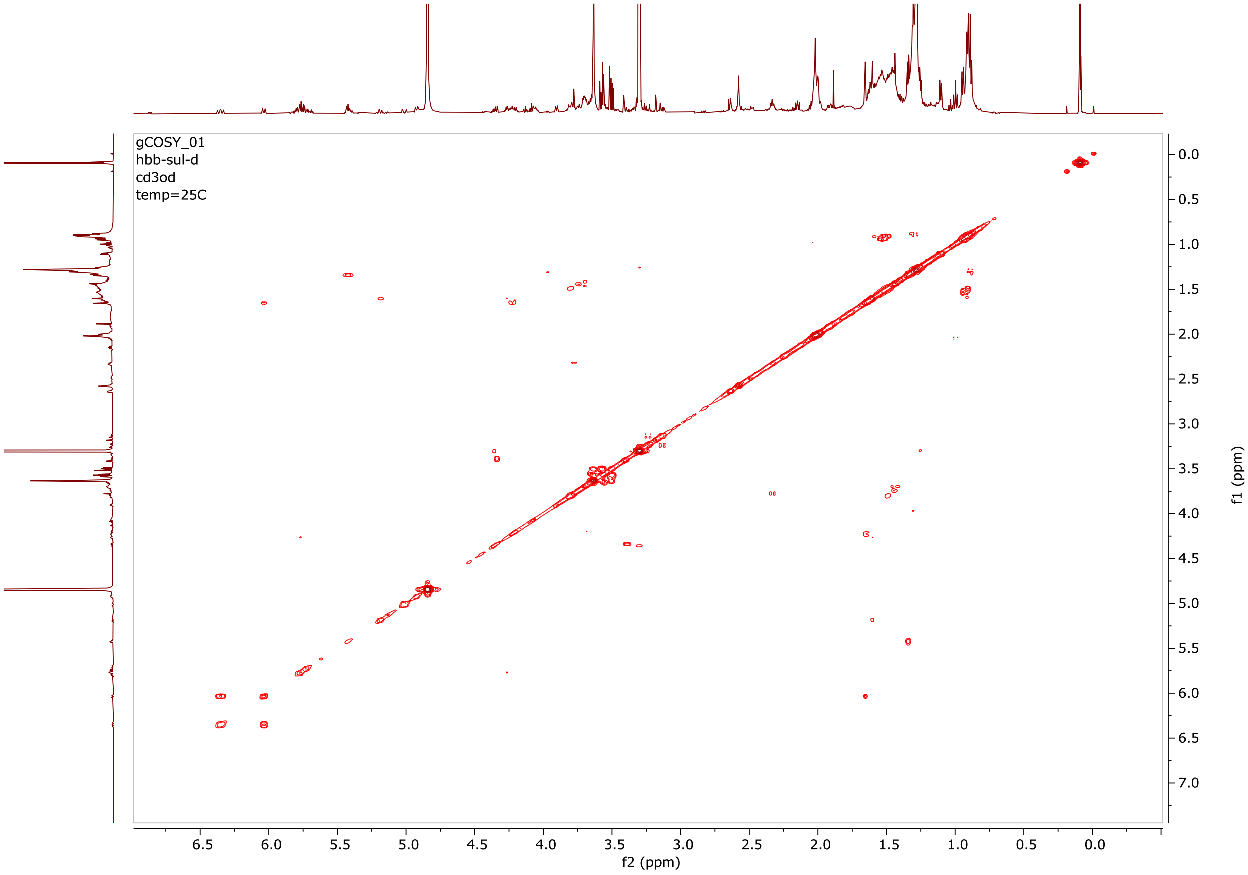


**Figure S25.** $^{1}$H-$^{1}$H COSY spectrum of deoxy-allyl-stambomycin (**3**).

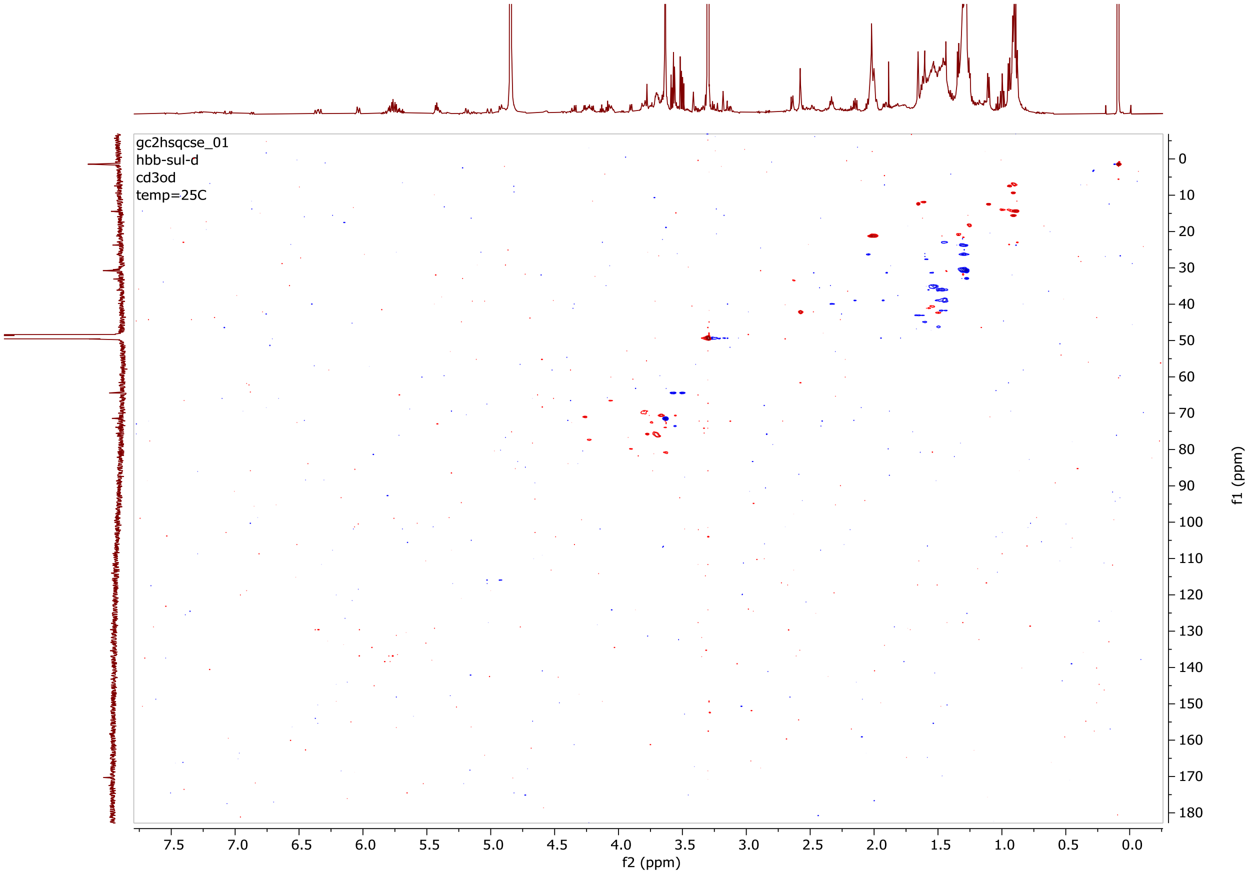


**Figure S26.** [$^{1}$H,$^{13}$C]-HSQC spectrum of deoxy-allyl-stambomycin (**3**).

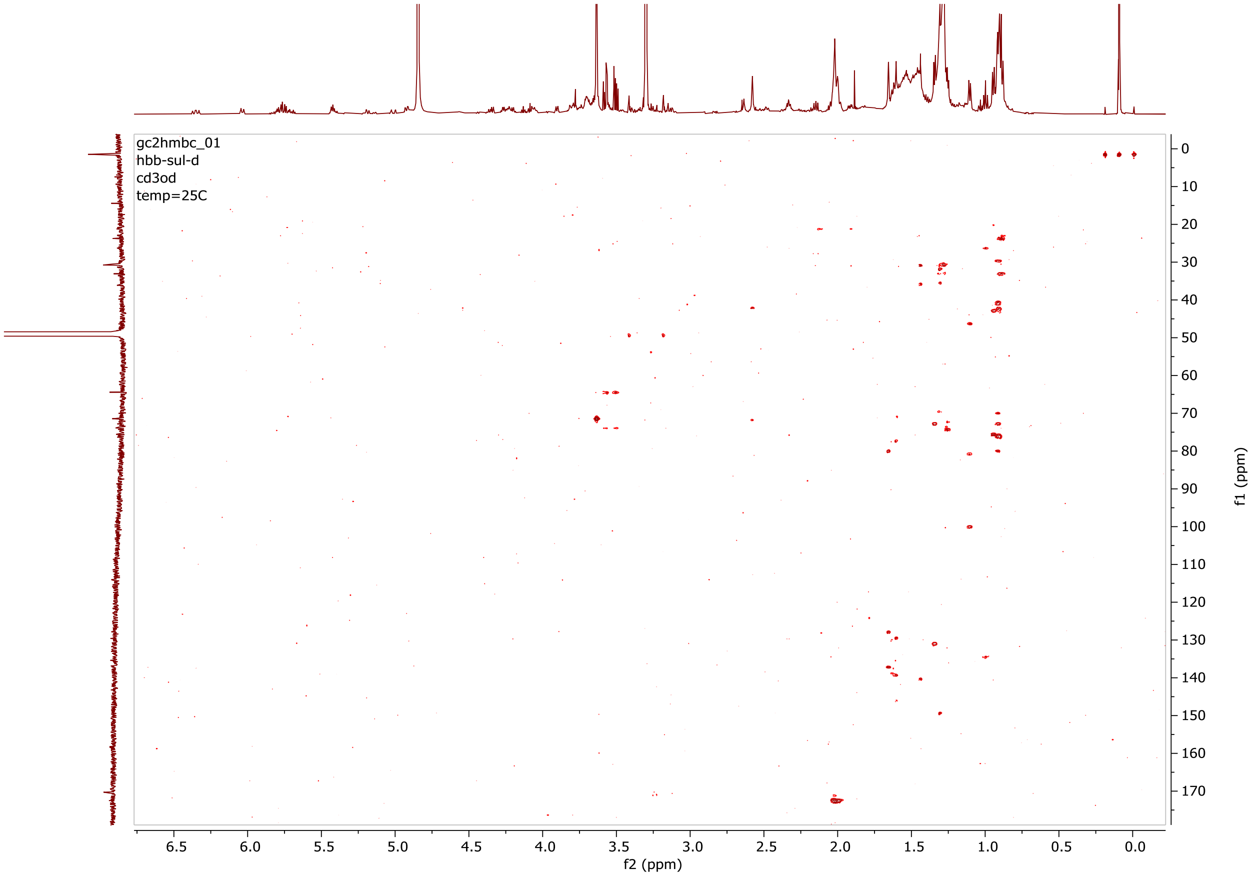


**Figure S27.** [$^{1}$H,$^{13}$C]-HMBC spectrum of deoxy-allyl-stambomycin (**3**).

Deoxy-allyl-stambomycin

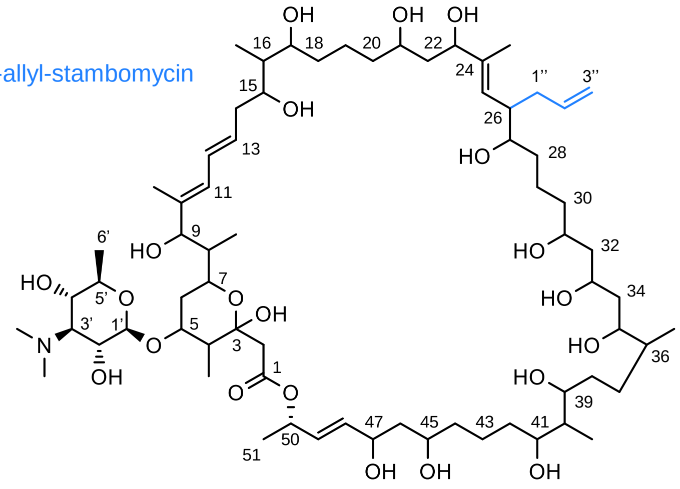


| Position | δC (ppm) | δH (ppm) | HMBC | Position | δC (ppm) | δH (ppm) | HMBC |
|---|---|---|---|---|---|---|---|
| 1 | 172.2 | | | 24-Me | 11.8 | 1.60 | |
| 2 | 43.0 | 1.62 | | 25 | 129.6 | 6.34 | 15.5, 10.9 |
| 3 | 100.5 | | | 26 | 42.1 | 2.57 | |
| 4 | 45.0 | 1.60 | | 1″ | 33.1 | 1.28 | |
| 4-Me | 12.4 | 1.10 | | 2″ | 134.4 | 5.2 | |
| 5 | 80.8 | 3.63 | | 3″ | 115.9 | 4.93 | 10.0 |
| 6 | 39.7 | 1.80 | | 27 | 76.2 | 3.69 | |
| 1′ | 106.7 | 4.33 | | 28 | 22.9 | 1.46 | |
| 2′ | 70.8 | 3.55 | 11.1, 6.0 | 29 | 30.7 | 1.28 | |
| 3′ | 71.6 | 3.64 | | 30 | 35.0 | 1.53 | |
| 3′-NMe$_2$ | 42.2 | 2.57 | | 31 | 69.3 | 3.79 | |
| 4′ | 71.4 | 3.64 | | 32 | 46.2 | 1.50 | |
| 5′ | 73.4 | 3.63 | | 33 | 67.4 | 3.61 | |
| 6′ | 18.2 | 1.26 | | 34 | 38.9 | 1.93 | |
| 7 | 69.8 | 3.81 | | 35 | 72.7 | 3.73 | |
| 8 | 41.6 | 1.47 | | 36 | 41.0 | 1.57 | |
| 8-Me | 9.2 | 0.91 | | 36-Me | 14.7 | 0.89 | |
| 9 | 79.9 | 3.90 | 7.4 | 37 | 30.7 | 1.28 | |
| 10 | 136.8 | | | 38 | 38.9 | 1.46, 1.94 | |
| 10-Me | 12.4 | 1.10 | | 39 | 75.9 | 3.70 | |
| 11 | 127.8 | 6.02 | 10.9, 14.9 | 40 | 41.7 | 1.57 | |
| 12 | 130.6 | 5.74 | | 40-Me | 6.7 | 0.89 | |
| 13 | 131.5 | 5.78 | | 41 | 72.3 | 3.31 | |
| 14 | 41.6 | 1.44 | | 42 | 33.7 | 159 | |
| 15 | 75.8 | 3.70 | | 43 | 21.2 | 2.01 | |
| 16 | 42.8 | 1.53 | | 44 | 39.0 | 1.46, 1.93 | |
| 16-Me | 7.3 | 0.94 | | 45 | 71.8 | 3.64 | |
| 17 | 76.7 | 3.69 | | 46 | 46.2 | 1.50 | |
| 18 | 36.0 | 1.57, 1.47 | | 47 | 70.4 | 3.67 | |
| 19 | 23.7 | 1.30 | | 48 | 136.8 | 5.77 | |
| 20 | 38.5 | 1.44 | | 49 | 130.9 | 5.74 | |
| 21 | 70.8 | 3.55 | | 50 | 72.7 | 5.42 | 6.3 |
| 22 | 45.0 | 1.60 | | 51 | 20.8 | 1.34 | 6.3 |
| 23 | 77.1 | 4.24 | | | | | |
| 24 | 139.1 | | | | | | |

**Figure S28.** Structure and summary for the NMR data for deoxy-allyl-stambomycin (**3**). Globally, the NMR data resemble those for deoxy-butyl-stambomycin, with the exception of those corresponding to the side chain attached to C-26. Specifically, we observe differences for the chemical shifts of C-2″ ($\delta_C$ 134.4) and C-3″ ($\delta_C$ 115.9) corresponding to the double bond of the allyl group. Although many of the expected $^{13}C$ signals are missing from $^{13}C$ NMR spectrum, close inspection of the cross peaks of the [$^1H$,$^{13}C$]-HSQC and [$^1H$,$^{13}C$]-HMBC spectra allowed their assignment. Together, these data confirmed the identity of the compound as deoxy-allyl-stambomycin.

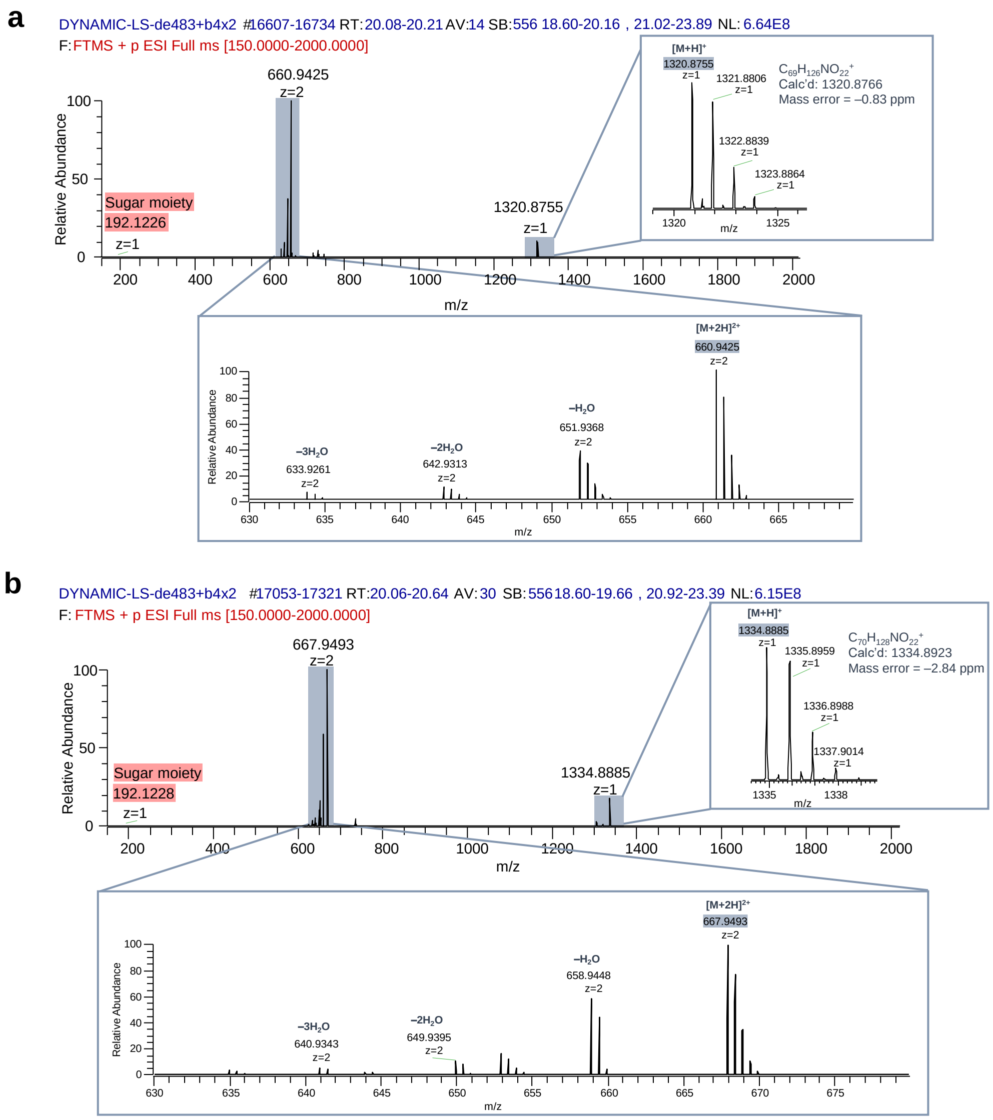

a
DYNAMIC-LS-de483+b4x2 #16607-16734 RT: 20.08-20.21 AV: 14 SB: 556 18.60-20.16 , 21.02-23.89 NL: 6.64E8
F: FTMS + p ESI Full ms [150.0000-2000.0000]
660.9425
z=2
1320.8755
z=1
Sugar moiety
192.1226
z=1
Relative Abundance
m/z
[M+H]+
1320.8755
z=1
1321.8806
z=1
1322.8839
z=1
1323.8864
z=1
C69H126NO22+
Calc'd: 1320.8766
Mass error = –0.83 ppm
[M+2H]2+
660.9425
z=2
–H2O
651.9368
z=2
–2H2O
642.9313
z=2
–3H2O
633.9261
z=2
b
DYNAMIC-LS-de483+b4x2 #17053-17321 RT: 20.06-20.64 AV: 30 SB: 556 18.60-19.66 , 20.92-23.39 NL: 6.15E8
F: FTMS + p ESI Full ms [150.0000-2000.0000]
667.9493
z=2
1334.8885
z=1
Sugar moiety
192.1228
z=1
Relative Abundance
m/z
[M+H]+
1334.8885
z=1
1335.8959
z=1
1336.8988
z=1
1337.9014
z=1
C70H128NO22+
Calc'd: 1334.8923
Mass error = –2.84 ppm
[M+2H]2+
667.9493
z=2
–H2O
658.9448
z=2
–2H2O
649.9395
z=2
–3H2O
640.9343
z=2

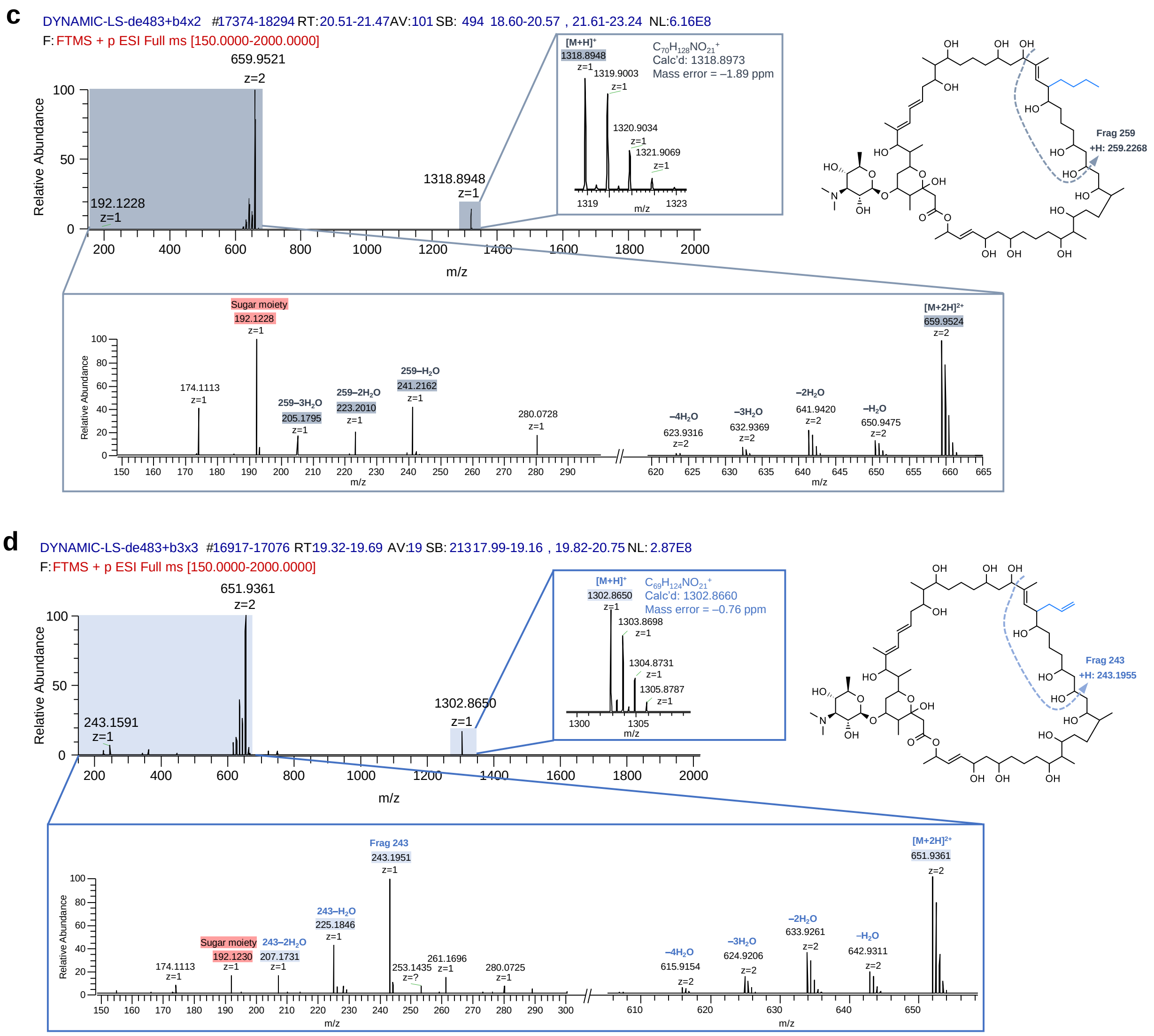


**Figure S29.** MS analysis of stambomycin analogues (**a**) C-24-demethyl-butylstambomycin (**12**), (**b**) butyl-stambomycin (**4**), (**c**) deoxy-butyl-stambomycin (**5**) and (**d**) deoxy-allyl-stambomycin (**3**). Similar to stambomycins A–F as shown previously[5], fragments were also identified which are diagnostic for the C-26 side chains (deoxy-allyl-stambomycin: fragment 243 and derived water loss fragments; deoxy-butyl-stambomycin: fragment 259 and derived water loss fragments). Together, these data provide further support for the structural assignments.

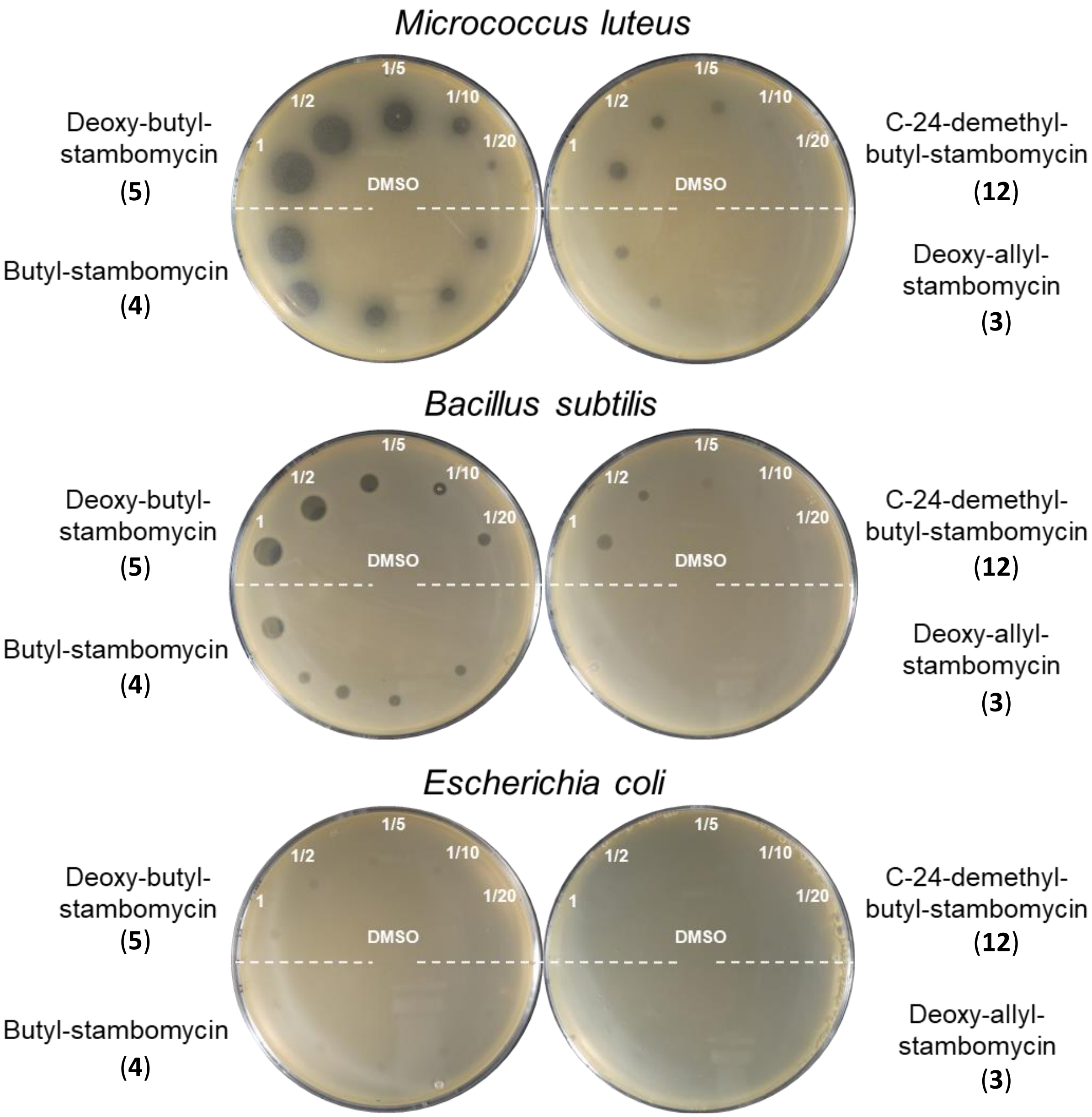


**Figure S30.** Antibacterial activities of the four purified stambomycin analogues (**3**, **4** ,**5** and **12**). For each analogue, 1 µL of sample (from a 10 mM stock solution in DMSO) either undiluted or at various dilutions as shown, was plated onto a confluent lawn of indicator strain (the Gram-positive bacteria *M. luteus* and *B. subtillis* or the Gram-negative bacterium *E. coli*). Pure DMSO (1 µL) was used as a control (plated in the centre).

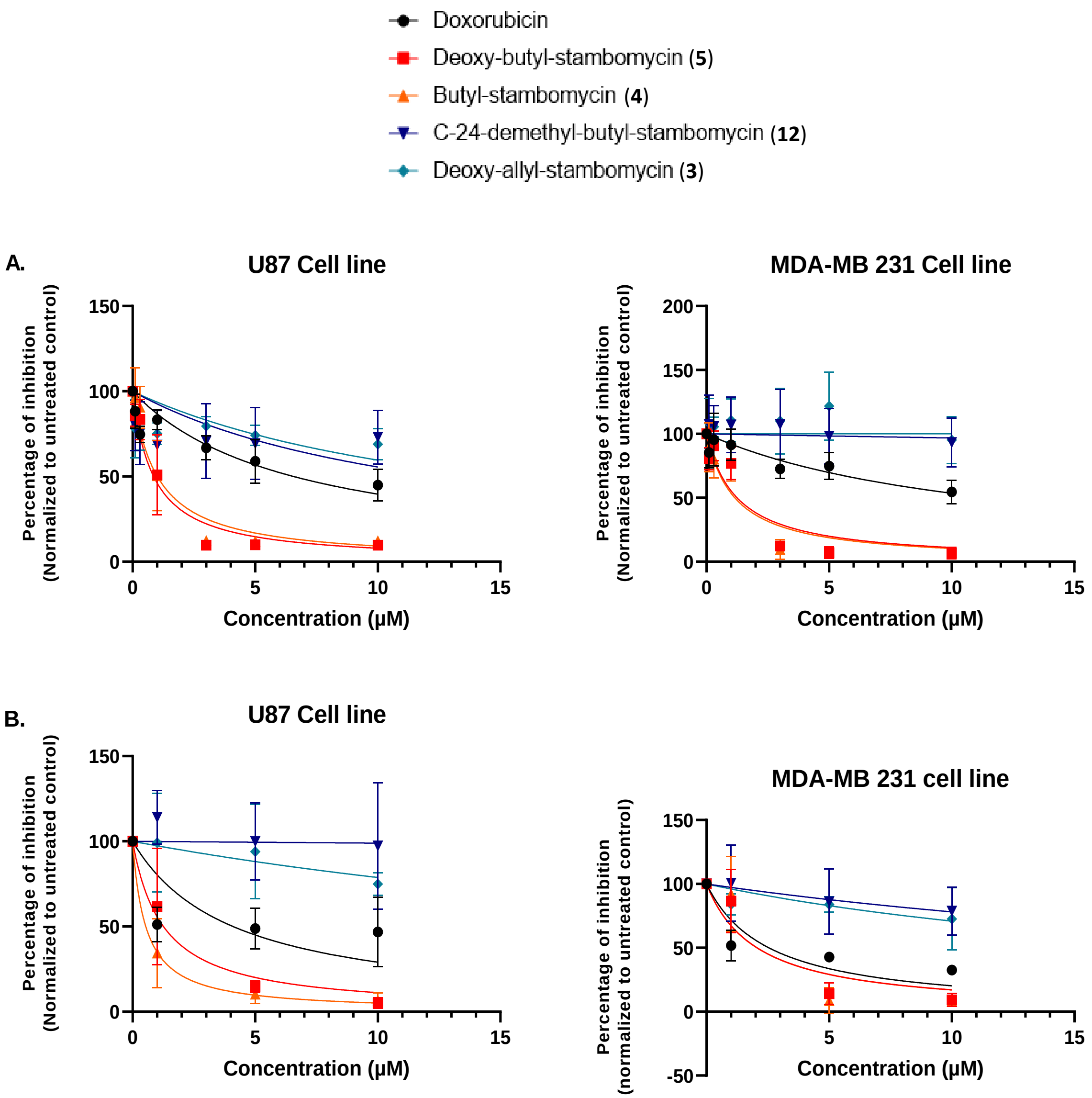


**Figure S31.** Antiproliferative activities of the four purified stambomycin analogues (**3**, **4** ,**5** and **12**) against two human cancer cell lines, U87-MG glioblastoma cells and MDA-MTB-231 cells. **A.** MTT assays and **B.** Cell counting. In both cases, the percentage of inhibition is normalised relative to the untreated sample. The clinical anticancer agent doxorubicin was used as a control. All the assays were performed in triplicate in three independent experiments.

**Table S1.** Homologs of SamR0482 and SamR0483 in *S. ambofaciens* ATCC23877 revealed by BlastP analysis.

| AA identities | Genome coordinates | Features | Accession number |
|---|---|---|---|
| Homologs of SamR0482 in *S. ambofaciens* ATCC23877 | | | |
| 29.8% | SAM23877_0423 | AntF (acyl-CoA ligase, involved in antimycin biosynthesis pathway) | AKZ53472.1 |
| 34.5% | SAM23877_6654 | Cgc22 (acyl-CoA synthetase, involved in congocidine biosynthesis pathway) | AKZ59699.1 |
| 27.8% | SAM23877_2750 | FadD (Long-chain-fatty-acid-CoA ligase) | AKZ55799.1 |
| 28.1% | SAM23877_3675 | AlkK (Medium-chain-fatty-acid-CoA ligase) | AKZ56720.1 |
| Homologs of SamR0483 in *S. ambofaciens* ATCC23877 | | | |
| 52% | SAM23877_0214 and SAM23877_7459 | AlpX (putative carboxyl transferase, involved in kinamycin biosynthesis) | AKZ53263.1 |
| 59% | SAM23877_4747 | PccB (propionyl-CoA carboxylase β chain) | AKZ57792.1 |
| 52% | SAM23877_5290 | PccB (propionyl-CoA carboxylase β chain) | AKZ58335.1 |
| 31% | SAM23877_2811 | Propionyl-CoA carboxylase | AKZ55860.1 |
| 29% | SAM23877_3346 | Methyl crotonyl-CoA carboxylase carboxyl transferase | AKZ56393.1 |
| 26% | SAM23877_2503 | Acetyl-coenzyme A carboxylase carboxyl transferase | AKZ55552.1 |

**Table S2.** List of strains and mutants.

| **Strains/mutants** | **Description and use (resistance)** | **Reference** |
|---|---|---|
| *Streptomyces* | | |
| ATCC/OE484 | *S. ambofaciens* ATCC23877 carrying the integrative plasmid pIB0484 (strain overexpressing the LAL regulator SamR0484) (Apra$^{R}$) | [4] |
| ATCC/OE484/Δ482 | PCR targeting mutant of ATCC/OE484 with the gene *samR0482* (SAM23877_7109, AKZ60152.1) replaced by a "*FRT+aadA+oriT+FRT*" cassette (Spec$^{R}$, Apra$^{R}$) | [13] |
| ATCC/OE484/Δ483 | PCR-targeting mutant of ATCC/OE484 with the gene *samR0483* (SAM23877_7108, AKZ60151.1) replaced by a scar (Apra$^{R}$) | This study |
| ATCC/pIB139/Δ483 | PCR-targeting mutant of ATCC/pIB139 with the gene *samR0483* replaced by a scar (Apra$^{R}$) | This study |
| ATCC/OE484/Δ483::*samR0483* | ATCC/OE484/Δ483 complemented with the conjugative and integrative plasmid pOSV809-PermE_*samR0483* (Kan$^{R}$, Apra$^{R}$) | This study |
| ATCC/pIB139/Δ483::*samR0483* | ATCC/pIB139/Δ483 complemented with the conjugative and integrative plasmid pOSV809_PermE_ *samR0483* (Kan$^{R}$, Apra$^{R}$) | This study |
| ATCC/OE483/Δ483/MatB_cinna | ATCC/OE484/Δ483 complemented with the conjugative and integrative plasmid pRT801_lacZ_PermE_MatB_cinna (Spec$^{R}$, Apra$^{R}$) | This study |
| *E. coli* | | |
| DH5α | General cloning strain | [14] |
| BW25113/pKD20 | PCR-targeting mutagenesis strain containing a λRED recombination plasmid pKD20 (Amp$^{R}$) | [15] |
| ET12567/pUZ8002 (ETU) | Non-methylating strain containing a mobilization plasmid for conjugation with *Streptomyces* (Kan$^{R}$, Cm$^{R}$) | [16] |
| S17-1 | The conjugative donor strain containing the *tra* gene on the chromosome | [17] |

**Table S3.** List of BACs and plasmids.

| BACs/plasmids | Properties and use (resistance) | Reference |
|---|---|---|
| BAC3 | BAB19ZF4 from the genomic library of *S. ambofaciens,* in which the chloramphenicol resistance gene is replaced by a spectinomycin resistance gene (Spec$^R$) | This study |
| BAC3_K7Δ483 | Mutant of BAC3 with gene *samR0483* replaced by a *"FRT+aadA+oriT+ FRT"* cassette (Spec$^R$, Apra$^R$) | This study |
| pIB139 | Conjugative and integrative vector (φC31 *attP-int, ermEp**) (Apra$^R$) | [18] |
| pOE484 | pIB139+*samR0484* (Apra$^R$) | [4] |
| pIJ778 | Origin of "*FRT+aadA+oriT+FRT*" cassette, (Spec$^R$) | [2] |
| pIJ773 | Origin of "*FRT+aac(3)IV+oriT+FRT*" cassette, (Apra$^R$) | [2] |
| pUWL-*oriT-flp* | Conjugative plasmid for "*FRT+aac(3)IV +oriT +FRT"* cassette excision (recombinase FLP) (Hyg$^R$) | [3] |
| pOSV809 | Conjugative and integrative vector (φBT1 *attP-int*) (Kan$^R$) | [19] |
| pOSV809_PermE_SamR483 | For complementation of the Δ*samR0483* mutant strain (Kan$^R$) | This study |
| pRT801_lacZ_PermE_MatB_cinna | Overexpression of MatB_cinna from *Streptomyces cinnamonensis* (Spec$^R$) | Gift from F Schulz, Ruhr-Universität Bochum, Germany |

**Table S4.** List of primers.

| Oligo name | Sequence | Usage |
|---|---|---|
| Spec_For | GAGTTATCGAGATTTTCAGGAGCTAAGGAAGCTAAAA TGAAGTCTACACGAACCCTTTG | Replacement of the chloramphenicol gene by spectinomycin |
| Spec_Rev | AGTGAGCTAACTCACATTAATTGCGTTGCGCTCACTG CCCTTATTTGCCGACTACCTTGG | Replacement of the chloramphenicol gene by spectinomycin |
| D483_For | GGAGGCAGGGTCGTCGTGTTGGAAGGTAGGGCTGGT ATGATTCCGGGGATCCGTCGACC | Deletion of *samR0483* |
| D483_Rev | GCCCCCGTACAGGAGCGGCCGGCACGACGAACGAC GTCATGTAGGCTGGAGCTGCTTC | Deletion of *samR0483* |
| F483_For | CGGTTCCGTGCCGTCCGATA | Verification of deleted *samR0483* |
| F483_Rev | AGACCTCCAGCGGCAAGGTG | Verification of deleted *samR0483* |
| ermEp_For | TGCGGCCGCTGCTAGCCGAGTGTCCGTTCGAGTGGC GGCTTG | For amplification of *ermEp**+RBS |
| ermEp_Rev | GGTCCTCCTGTGGAGTGGTGTGGATCCTACCAACCG GCACGATTG | For amplification of *ermEp**+RBS |
| Comp483_For | CACCACTCCACAGGAGGACCATGTCGCTCCAGGAGC CTGTCTCGC | For amplification of *samR0483* |
| Comp483_Rev | GCGGCCGCTACTAGTTCACAGGGGAATGTTCCCGTG TTTGCG | For amplification of *samR0483* |
| Ver483_For | CAAATGTAGCACCTGAAGTC | Verification of plasmid pOSV809-PermE_SamR483 |
| Ver483_Rev | GTTCGGCCCCTTTTTTGGCC | Verification of plasmid pOSV809-PermE_SamR483 |
| pOSV_For | GACTTCGCCCATCATGCGCTC | Verification of mutant ATCC/OE483/Δ483/*samR0483* and ATCC/pIB139/ Δ483:: *samR0483* |
| pOSV_Rev | GTGCTCAACGGGAATCCTGCTC | Verification of mutant ATCC/OE483/Δ483/*samR0483* and ATCC/pIB139/ Δ483:: *samR0483* |
| φBT1-attB_For | GACCTTGTGCTTGGTCGTCTTC | Verification of mutant ATCC/ OE483/Δ483/*samR0483* and ATCC/pIB139/Δ483::*samR0483* |
| φBT1-attB_Rev | GTAGATCGACAGGGCCATCCAC | Verification of mutant ATCC/ OE483/Δ483/*samR0483* and ATCC/pIB139/Δ483::*samR0483* |

Key: "NNNNNNN" and "NNNNNNN" represent sequences identical to those at the right or left ends of the disruption cassette in the PCR-targeting system. "GCTAGC" is the restriction recognition site of *Nhe*I, and "ACTAGT" is that for *Spe*I.

**Table S5.** Calculated and observed masses of all metabolites in this study.

| Compound | Chemical formula | Retention Time (min) | Calculated Mass $[M+2H]^{2+}$ | Observed Mass $[M+2H]^{2+}$ (mass error in ppm) | Calculated Mass $[M+H]^{+}$ | Observed Mass $[M+H]^{+}$ (mass error in ppm) |
|---|---|---|---|---|---|---|
| Stambomycin F | $C_{74}H_{135}NO_{22}$ | 24.2 | 694.9815 | n.d. | 1390.9549 | 1390.9517 (–2.30 ppm) |
| Stambomycin A/B | $C_{73}H_{133}NO_{22}$ | 22.8 | 688.9736 | 688.9697 (–5.66 ppm) | 1376.9392 | 1376.9358 (–2.47 ppm) |
| Stambomycin C/D | $C_{72}H_{131}NO_{22}$ | 21.7 | 681.9659 | 681.9620 (–5.72 ppm) | 1362.9236 | 1362.9207 (–2.13 ppm) |
| Stambomycin E | $C_{71}H_{129}NO_{22}$ | 21.3 | 674.9539 | 674.9531 (–1.19 ppm) | 1348.9058 | 1348.9046 (–1.11 ppm) |
| Demethyl-stambomycin E | $C_{70}H_{127}NO_{22}$ | 20.5 | 667.9502 | 667.9493 (–1.35 ppm) | 1334.8923 | 1334.8911 (–0.89 ppm) |
| Ethyl-stambomycin | $C_{68}H_{123}NO_{22}$ | n.d. | 653.9345 | n.d. | 1306.8610 | n.d. |
| Deoxy-ethyl-stambomycin (**1**) | $C_{68}H_{123}NO_{21}$ | 19.1 | 645.9345 | n.d. | 1290.8660 | 1290.8647 (–1.00 ppm) |
| Isopropyl-stambomycin | $C_{69}H_{125}NO_{22}$ | n.d. | 660.9419 | n.d. | 1320.8766 | n.d. |
| Deoxy-isopropyl-stambomycin | $C_{69}H_{125}NO_{21}$ | n.d. | 652.9445 | n.d. | 1304.8817 | n.d. |
| Allyl-stambomycin (**2**) | $C_{69}H_{123}NO_{22}$ | 19.3 | 659.9345 | n.d. | 1318.8610 | 1318.8606 (–0.30 ppm) |
| Deoxy-allyl-stambomycin (**3**) | $C_{69}H_{123}NO_{21}$ | 19.5 | 651.9370 | 651.9361 (–1.38 ppm) | 1302.8660 | 1302.8650 (–0.76 ppm) |
| C-24-demethyl-butyl-stambomycin (**12**) | $C_{69}H_{125}NO_{22}$ | 19.9 | 660.9425 | 660.9420 (–0.75 ppm) | 1320.8766 | 1320.8755 (–0.83 ppm) |
| Butyl-stambomycin (**4**) | $C_{70}H_{127}NO_{22}$ | 20.3 | 667.9502 | 667.9493 (–1.35 ppm) | 1334.8923 | 1334.8885 (–2.84 ppm) |
| Deoxy-butyl-stambomycin (**5**) | $C_{70}H_{127}NO_{21}$ | 20.9 | 659.9527 | 659.9521 (–0.91 ppm) | 1318.8973 | 1318.8948 (–1.89 ppm) |
| Benzyl-stambomycin (**6**) | $C_{73}H_{125}NO_{22}$ | 21.5 | 684.9580 | 684.9417 (–23.8 ppm) | 1368.8766 | 1368.8782 (1.17 ppm) |
| Deoxy-benzyl-stambomycin (**7**) | $C_{73}H_{125}NO_{21}$ | 21.5 | 676.9423 | 676.9438 (2.22 ppm) | 1352.8817 | 1352.8781 (–2.66 ppm) |
| Octyl-stambomycin (**8**) | $C_{74}H_{135}NO_{22}$ | 25.4 | 695.9811 | 695.9805 (–0.86 ppm) | 1390.9549 | 1390.9546 (–0.22 ppm) |
| Deoxy-octyl-stambomycin | $C_{74}H_{135}NO_{21}$ | n.d. | 687.9836 | n.d. | 1374.9599 | n.d. |
| Phenylpropyl-stambomycin | $C_{75}H_{129}NO_{22}$ | n.d. | 698.9555 | n.d. | 1396.9079 | n.d. |
| Deoxy-phenylpropyl-stambomycin | $C_{75}H_{129}NO_{21}$ | n.d. | 690.9602 | n.d. | 1380.9130 | n.d. |
| Phenoxypropyl-stambomycin (**9**) | $C_{75}H_{129}NO_{23}$ | 22.5 | 706.9554 | n.d. | 1412.9028 | 1412.8993 (–2.48 ppm) |
| Deoxy-phenoxypropyl-stambomycin (**10**) | $C_{75}H_{129}NO_{22}$ | 22.5 | 698.9555 | n.d. | 1396.9079 | 1396.9048 (–2.22 ppm) |
| Thiophene-stambomycin | $C_{70}H_{121}NO_{22}S$ | n.d. | 680.9123 | n.d. | 1360.8174 | n.d. |
| Deoxy-thiophene-stambomycin | $C_{70}H_{121}NO_{21}S$ | n.d. | 672.9148 | n.d. | 1344.8225 | n.d. |
| Thienylmethyl-stambomycin | $C_{71}H_{123}NO_{22}S$ | n.d. | 687.9201 | n.d. | 1374.8330 | n.d. |
| Deoxy-thienylmethyl-stambomycin | $C_{71}H_{123}NO_{21}S$ | n.d. | 679.9227 | n.d. | 1358.8381 | n.d. |
| Propargyl-stambomycin | $C_{69}H_{121}NO_{22}$ | n.d. | 658.9263 | n.d. | 1316.8453 | n.d. |
| Deoxy-propargyl-stambomycin | $C_{69}H_{121}NO_{21}$ | n.d. | 650.9289 | n.d. | 1300.8504 | n.d. |
| 6-Bromohexyl-stambomycin (**11**) (lacking 2H) | $C_{72}H_{128}NO_{22}Br$ | 22.6 | 719.9129 ($Br^{79}$) 720.9130 ($Br^{81}$) | 719.9128 ($Br^{79}$) (–0.14 ppm) 720.9130 ($Br^{81}$) (–0 ppm) | 1438.8184 ($Br^{79}$) 1440.8187 ($Br^{81}$) | 1438.8164 ($Br^{79}$) (–1.53 ppm) 1440.8183 ($Br^{81}$) (–0.28 ppm) |
| Deoxy-6-bromohexyl-stambomycin (lacking 2H) | $C_{72}H_{128}NO_{21}Br$ | n.d. | 711.9154 ($Br^{79}$) 712.9155 ($Br^{81}$) | n.d. | 1422.8235 ($Br^{79}$) 1424.8238 ($Br^{81}$) | n.d. |

n.d. = not detected

**Table S6.** Quantification of stambomycins in parental and mutasynthesis strains.

| Strains (culture volume) | Integrated peak areas for stambomycins A, B, C, D (extracted ion (EI) = 1362.9236, 681.9659, 1376.9392, 688.9736) | Volume of final extract in MeOH (μL) | Calculated titre (mg $L^{-1}$) | Average ± deviation (parental level, %) |
|---|---|---|---|---|
| 20190130-wtOE (50 mL) | 4672442552 | 300 | 28.2 | 30 ± 2 (100%) |
| 20190913-wtOE (50 mL) | 6621933694 | 240 | 31.7 | |
| 20190130-de482OE#1 (50 mL) | 3117290008 | 240 | 15.0 | 15.2 ± 0.3 (50.8%) |
| 20190130-de482OE#2(50 mL) | 2550403337 | 300 | 15.4 | |
| 20190130-de483OE (50 mL) | 22253469 | 270 | 0.12 | 0.09 ± 0.04 (0.31%) |
| 20190206-de483OE (50 mL) | 23710170 | 270 | 0.13 | |
| 20190226-de483OE#1 (50 mL) | 10444147 | 220 | 0.049 | |
| 20190226-de483OE#2 (50 mL) | 23826563 | 240 | 0.12 | |
| 20190430-de483OE (50 mL) | 12248726 | 200 | 0.052 | |
| 20190913-de483OE (50 mL) | 12525726 | 330 | 0.088 | |
| 20191108-de483OE::483#1 (50 mL) | 2125375050 | 200 | 8.56 | 9.2 ± 0.9 (30.8%) |
| 20191108-de483OE::483#2 (50 mL) | 2337747862 | 210 | 9.88 | |

**Table S7.** Titres (mg $L^{-1}$) of stambomycins and analogues calculated using the relative amounts shown in **Table 1**. The titre of stambomycins A–D produced by ATCC/OE484/Δ483 was 0.09 mg $L^{-1}$, as shown in **Table S6**.

| **Fed malonic acid derivatives** | **Titres of stambomycins and its analogues (mg $L^{-1}$)** | | |
|---|---|---|---|
| | Stambomycins (A–D) | Analogue (with C-28 -OH) | Analogue (without C-28 -OH) |
| Ethyl- | 0.77 | n.d. | 0.00077 (**1**) |
| Isopropyl- | 0.32 | n.d. | n.d. |
| Allyl- | 0.17 | 0.0018 (**2**) | 0.34 (**3**) |
| Butyl- | 0.17 | 3.9 (**4**) | 18.56 (**5**) |
| Benzyl- | 0.43 | 0.025 (**6**) | 0.049 (**7**) |
| Octyl- | 0.76 | 0.07 (**8**) | n.d. |
| Phenylpropyl- | 0.53 | n.d. | n.d. |
| Phenoxylpropyl- | 0.21 | 0.0098 (**9**) | 0.012 (**10**) |
| Thiophene- | 0.08 | n.d. | n.d. |
| Thienylmethyl- | 0.09 | n.d. | n.d. |
| Propargyl- | 0.41 | n.d. | n.d. |
| 6-Bromoethyl- | 0.55 | 0.025 (**11**) | n.d. |

**Table S8.** Conditions for the purification of mutasynthetic stambomycin analogues.

| **1. Prep HPLC** | |
|---|---|
| Solvents | ACN/$H_2O$ +0.1%FA |
| 0 min | 10% |
| 2 min | 10% |
| 27 min | 50% |
| Column | Waters XBridge BEH C18 19x250mm |
| Detection UV | 205 mAU |
| Retention time of deoxy-butyl-stambomycin (**5**) | 18.5 min |
| Retention time of butyl-stambomycin (**4**) | 17.6 min |
| Retention time of C-24-demethyl-butyl stambomycin (**12**) | 17.1 min |
| Retention time of deoxy-allyl-stambomycin (**3**) | 16.9 min |
| **Amounts of purified compounds** | |
| Deoxy-butyl-stambomycin (**5**) | 36.3 mg |
| Butyl-stambomycin (**4**) | 11.6 mg |
| C-24-demethyl-butyl-stambomycin (**12**) | 3.1 mg |
| Deoxy-allyl-stambomycin (**3**) | 1.8 mg |

**Table S9.** IC$_{50}$ values of the stambomycin analogues for two human cancer cell lines. The U87-MG cell line correspond to glioblastoma cells (brain cancer) and MDA-MTB-231 cells are commonly used to model late-stage breast cancer. The clinical anticancer agent doxorubicin was used as a control. IC$_{50}$ indicates the concentration needed to inhibit the growth of 50% of cells in the population.

| | IC50 (µM) | U87 cell line | | | | | MDA-MB-231 cell line | | | | |
|---|---|---|---|---|---|---|---|---|---|---|---|
| | | Doxorubicin | Deoxy-butyl-stambomycin | Butyl-stambomycin | C-24-demethyl-butyl-stambomycin | Deoxy-allyl-stambomycin | Doxorubicin | Deoxy-butyl-stambomycin | Butyl-stambomycin | C-24-demethyl-butyl-stambomycin | Deoxy-allyl-stambomycin |
| MTT assay | Experiment 1 | 9,39 | 1,09 | 1,29 | 6,66 | 12,67 | 8,90 | 0,98 | 0,73 | > 1 mM | > 1 mM |
| | Experiment 2 | 5,90 | 0,78 | 0,66 | 5,48 | 25,38 | 9,49 | 1,52 | 1,45 | 26,39 | 47,51 |
| | Experiment 3 | 5,17 | 0,76 | 1,15 | 74,47 | 10,55 | 18,58 | 1,23 | 1,31 | 71,35 | > 1 mM |
| | **Mean** | **6,82** | **0,88** | **1,03** | **28,87** | **16,20** | **12,32** | **1,24** | **1,16** | **N.A.** | **N.A.** |
| | **SD** | **2,25** | **0,19** | **0,33** | **39,50** | **8,02** | **5,43** | **0,27** | **0,38** | | |
